# Resolution-aware spectral parameterization reveals robust context-dependent aperiodic EEG dynamics during real-world table tennis

**DOI:** 10.64898/2026.10.01.755924

**Authors:** Arunim Guchait, Chiao-Yun Chen, Neil G. Muggleton

## Abstract

Mobile EEG can link neuroimaging to natural behavior, but effects recorded during movement, and sport in particular, are difficult to interpret because neural activity overlaps with signals from movement, muscle activity, and cable movement artifacts. We analyzed the OpenNeuro Real World Table Tennis dataset to test whether peri-hit spectral activity could be analyzed and how it differed between human-opponent and ball-machine play. Data from 25 participants comprising EEG, noise-layers, EMG and motion recordings was analyzed with pre-hit, impact, and post-hit windows. EEG power bands were residualized using trial-level root mean square from noise-layer, EMG, and motion/miscellaneous channels. Spectral parameterization modeled spectra as an aperiodic component plus periodic peaks. The principal short-window model was resolution-aware; sensitivity analyses reduced model complexity, used a 2.0-s peri-hit subwindow (−1.0 to +1.0 s), or disabled periodic fitting. Results showed residualized alpha and beta power was lower during ball-machine play, including pre-hit posterior alpha power. The clearest context difference was a higher ball-machine aperiodic exponent over posterior and parietal sensors, corresponding to a steeper broadband spectral slope. Posterior post-hit means were 0.451 and 0.606 for human-opponent and ball-machine play, respectively. A focal parietal effect persisted across specifications. The results support artifact-informed, resolution-aware spectral analysis as an effective approach for detecting robust spectral differences between naturalistic visuomotor contexts and suggest that EEG may provide useful information about the processes engaged by different practice environments, without establishing a single cognitive or physiological mechanism or demonstrating a training outcome.

## 1 Introduction

### 1.1 Real-world neuroimaging and the problem of movement

Much of cognitive neuroscience has advanced through experiments in which stimuli, responses, and environmental conditions are carefully controlled. This approach has been essential for isolating particular perceptual, cognitive, and motor processes. However, experimental control is often achieved by restricting movement and simplifying the environment, meaning that the resulting neural measurements may not fully represent cognition when perception and action are continuously coupled to the body and the surrounding world. Mobile brain/body imaging was developed partly in response to this limitation, allowing electrophysiological activity to be examined together with movement and behavior under more natural conditions (Gramann et al., 2014; Makeig et al., 2009). This approach does not replace controlled laboratory neuroscience. Rather, it provides complementary evidence about how neural processes are expressed when participants move through and respond to dynamic environments.

The importance of this issue is particularly apparent for visuomotor behavior. In many conventional experiments, visual perception, response preparation, movement execution, and outcome evaluation are analyzed as distinct trial stages, even though the underlying processes may overlap. During interception, successful action requires the performer to extract information about the moving object, anticipate where and when contact is likely to occur, prepare an appropriate movement, and continue updating that movement as new visual information becomes available (Bootsma & van Wieringen, 1990; Brenner & Smeets, 2018; Fooken et al., 2016). These perceptual and motor operations are coupled across time rather than completed as independent steps, and their relative contributions can change as the target approaches. Sport provides a useful setting for investigating such processes because it combines natural movement with identifiable events around which neural activity can be analyzed.

Table tennis is well suited to this type of investigation. The task requires rapid visual tracking, trajectory prediction, object interception, whole-body coordination, racket control, and performance monitoring. The public Real World Table Tennis dataset was designed to support investigation of these processes and contains recordings from 25 participants completing real table-tennis play with both a human opponent and a ball machine. The dataset includes 120 scalp EEG electrodes, 120 mechanically matched noise-layer electrodes, eight neck electromyography electrodes, and synchronized inertial measurements from the participant and task equipment (Studnicki & Ferris, 2024). This multimodal structure is particularly valuable because it allows event-related scalp activity to be considered alongside signals that may reflect movement, muscle activity, cable motion, and other non-neural contributions.

The possibility of recording during natural movement does not, however, remove the methodological difficulties associated with analysis of mobile-acquired EEG. Whole-body sport produces several forms of physiological and non-physiological contamination of the EEG signal, including neck and facial muscle activity, electrode displacement, cable movement, and changes associated with head, arm, and trunk motion. These signals may overlap with the frequencies and scalp distributions used to interpret neural activity. The dual-layer table-tennis work showed that mechanically matched noise electrodes provide useful information about movement-related artifacts and that incorporating these reference signals can improve the quality of recovered brain components (Studnicki et al., 2022). Consequently, EEG recorded during table tennis cannot be interpreted as though the participant were stationary. Any context comparison requires an analysis strategy that treats artifact-related signals as part of the scientific problem rather than as an issue resolved solely by conventional filtering.

### 1.2 Human-opponent and ball-machine play as distinct visuomotor contexts

The availability of both human-opponent and ball-machine trials provides a structured comparison between two forms of real-world object interception. Both contexts presumably require the participant to track an approaching ball, prepare a stroke, contact the ball with the racket, and continue responding to the developing exchange. The contexts therefore seem comparable in their broad visuomotor requirements. However, the source and structure of the information preceding the incoming ball differ substantially.

During ball-machine play, the ball is produced by a mechanical device. Its speed, direction, spin, and timing can vary, but the participant does not receive advance biological-motion information from the body or racket movement of another person. During human-opponent play, the incoming trajectory is preceded by information from the opponent’s posture, arm movement, racket orientation, movement timing, and strategic behavior. Expert performance in interceptive sports is associated with the ability to extract useful advance information from such kinematic cues, and skilled performers can use information available before object release or contact to anticipate the likely outcome of an action (Abernethy & Zawi, 2007; Mann et al., 2007). Human-opponent play may therefore provide a richer anticipatory context, whereas ball-machine play may place greater emphasis on the observable trajectory and regularities of mechanical delivery.

This distinction should not be reduced to a claim that one context is simply more difficult or more cognitively demanding than the other. Difficulty is not a single task property, and the available dataset does not provide a direct measure capable of establishing that one condition required greater attention, effort, or engagement. Human-opponent and ball-machine play differ simultaneously in predictability, interpersonal timing, biological motion, strategic uncertainty, delivery structure, and potentially movement execution. A neural difference between the processing in different contexts could therefore be associated with one or several of these properties. It could also be influenced by differences in stroke selection, stance, head movement, muscle activation, or remaining differences in movement-related or other recording artifacts that were not fully captured by the nuisance measures. The appropriate comparison is consequently between two forms of perception–action coupling, rather than between a generally easy and a generally difficult condition.

### 1.3 Previous analyses and the contribution of the present study

Studnicki et al. (2022) initially used the dual-layer recordings to characterize movement artifacts during table tennis and to compare approaches for improving EEG cleaning. That work demonstrated the methodological value of the matched noise layer during a discrete but whole-body sporting task. Studnicki and Ferris (2024) subsequently released the recordings as a reusable multimodal dataset, documenting the scalp EEG, noise-layer EEG, neck electromyography, inertial measurements, anatomical MRI, electrode locations, events, and available video resources. These contributions established both the technical basis for recording during table tennis and the infrastructure required for secondary analyses.

The same database has also been used to examine longer-timescale EEG changes across rest, gameplay, and recovery. Iammarino et al. (2026), for example, evaluated the temporal behavior of multiple bandpower-based engagement indexes during different phases of the recording session. Such work addresses whether broad psycho-emotional or engagement-related measures change across sustained sporting activity. The present analysis addresses a different question by focusing on shorter intervals time-locked to participant racket–ball contact and by directly comparing human-opponent with ball-machine play. It also avoids relying exclusively on band ratios, because ratios and canonical band measures may combine changes in narrowband oscillations with changes in the broadband background of the spectrum (Donoghue, Dominguez, et al., 2020).

The contribution of the present study is therefore not the acquisition of the dataset or the demonstration that EEG can be recorded during table tennis. Instead, the analysis tests whether human-opponent and ball-machine contexts are associated with different peri-hit spectral states, whether canonical bandpower differences remain after accounting for available nuisance signals, and whether the dominant context effects are better characterized as periodic or aperiodic spectral changes. The comparison is conducted at the participant level so that the effective inferential sample reflects the 25 paired participants rather than the much larger number of non-independent event epochs.

This comparison also has potential relevance to the study of training. Practice frequently simplifies, constrains, or otherwise alters the situation for which performance is ultimately being trained, and these changes may affect not only behavior but also the perceptual and cognitive processes engaged during practice. Representative learning design emphasizes the importance of preserving functionally relevant information and perception–action coupling when practice is intended to transfer to a target setting (Pinder et al., 2011), while the broader transfer literature shows that similarity between training and later performance cannot be reduced to a single dimension (Barnett & Ceci, 2002). From this perspective, EEG may provide information that complements behavioral performance by indicating whether two superficially related practice contexts are associated with similar or different patterns of neural processing. The present study does not test training transfer directly, but the human-opponent and ball-machine comparison provides a useful setting in which to examine this possibility.

### 1.4 Rationale for separating periodic and aperiodic spectral activity

Electrophysiological analysis has traditionally emphasized canonical frequency bands such as theta, alpha, and beta. This approach is useful because rhythmic neural activity can reflect the organization of the timing of neuronal populations and has of utility in investigation of perception, attention, memory, and sensorimotor processing (Buzsaki & Draguhn, 2004). In the context of table tennis, changes in posterior, parietal, and sensorimotor spectral power could plausibly accompany visual prediction, movement preparation, racket–ball contact, and post-hit evaluation.

Bandpower is not a component-specific measure. Power within a predefined band may increase because the amplitude of a genuine periodic oscillation has increased. It may also change because the center frequency or bandwidth of an oscillatory component has shifted, or because the broader aperiodic component of the spectrum has changed. Electrophysiological spectra contain an arrhythmic, approximately 1/f-like component in addition to periodic peaks. Variation in the offset or exponent of this aperiodic component can alter measured power across several canonical bands simultaneously, even when no selective oscillatory change has occurred (Donoghue, Haller, et al., 2020; He, 2014). This creates a particular interpretive problem for mobile EEG because broadband spectral changes may also be sensitive to movement, muscle activity, and recording quality.

Spectral parameterization provides a way to examine this problem more explicitly. The approach represents a neural power spectrum as the combination of an aperiodic component and putative periodic peaks, allowing the exponent and offset of the aperiodic component to be estimated separately from the center frequency, power, and bandwidth of periodic components (Donoghue, Haller, et al., 2020). This does not make every estimated component uniquely neural, nor does it assign a single physiological meaning to the aperiodic exponent. It does, however, reduce the risk that a broad change in spectral slope will be interpreted as though it necessarily reflects an, for example, alpha- or beta-specific oscillation.

The aperiodic exponent has been related to properties of population-level neural activity, and modelling work has suggested that changes in the slope of electrophysiological spectra may reflect changes in the balance of excitatory and inhibitory synaptic activity under some experimental conditions (Gao et al., 2017). Such an interpretation remains several inferential steps removed from scalp EEG recorded during active sport. An exponent measured at the scalp is affected by source mixing, reference choice, preprocessing, spectral estimation, and non-neural contamination. It should therefore not be treated as a direct measure of cortical excitation–inhibition balance, attention, arousal, or cognitive effort. The more defensible interpretation is that it describes a component of broadband spectral architecture that may vary with task context.

This distinction is also relevant to composite measures such as band ratios and engagement indexes. A ratio can change because either frequency band changes, because a periodic peak moves across a band boundary, or because aperiodic slope alters power across the numerator and denominator. Simulations and empirical analyses have shown that band ratios can therefore conflate several periodic and aperiodic spectral properties (Donoghue, Dominguez, et al., 2020). In a task involving simultaneous cognitive, motor, muscular, and mechanical processes, interpretation based solely on canonical power or ratios would consequently be incomplete.

### 1.5 Temporal specificity and spectral resolution

Event-locked spectral analysis also involves a necessary compromise between temporal specificity and frequency resolution. Short analysis windows are needed to distinguish activity occurring before racket–ball contact from activity around impact and after the hit. Combining these periods into a single long interval would obscure whether a context difference was associated more strongly with anticipatory processing, movement and contact, or subsequent evaluation. However, reducing the duration of the analyzed signal also produces a coarser frequency grid. Welch estimation improves the stability of power spectra by averaging modified periodograms, but the spectral information available from each segment remains constrained by its duration (Welch, 1967). Time-resolved spectral-parameterization work similarly demonstrates that dynamic periodic and aperiodic features can be estimated from short windows, while also showing that temporal localization and spectral precision must be balanced when defining those windows (Wilson et al., 2022).

This limitation is especially important for periodic-peak estimation. A broad aperiodic slope can be estimated across multiple frequency bins, whereas the center frequency and bandwidth of a narrow peak depend more directly on the density of the spectral grid. In the present analysis, the pre-hit, impact, and post-hit windows were 0.8, 0.5, and 0.7 s in duration, producing frequency spacings of approximately 1.24, 1.98, and 1.42 Hz, respectively. A model that permits peaks narrower than the spacing of the frequencies entered into the model may allocate isolated spectral points to putative oscillatory components. Peak-width limits should therefore be selected in relation to the frequency resolution of each spectrum. The specparam guidance recommends that the minimum permitted bandwidth be at least approximately twice the frequency resolution, reducing the possibility that a single frequency point will be fitted as a periodic peak.

Short-window estimates provide temporal specificity at the cost of reduced spectral resolution, with the greatest limitation applying to the characterization of narrow periodic peaks. This trade-off motivated a resolution-aware short-window model in which the lower peak-width limit was adapted to the frequency grid of each event interval. A separate two-second peri-hit model was also used to provide a better-resolved sensitivity analysis, with a frequency spacing of approximately 0.50 Hz. The two-second analysis estimates a broader peri-hit spectral difference and should therefore be treated as a related but distinct temporal estimand rather than as another measurement of exactly the same effect.

### 1.6 Aims and hypotheses

The first aim of the study was to determine whether human-opponent and ball-machine play were associated with different event-locked EEG spectral activity around participant racket–ball contact. The comparison was performed separately for pre-hit, impact, and post-hit windows because the perceptual and motor operations surrounding a hit are unlikely to remain constant across the full peri-action interval. Context effects were expected to be most evident over posterior and parietal sensors, given the importance of visual prediction, spatial processing, and visuomotor transformation during object interception. This was a sensor-level hypothesis and was not intended to imply localization to a specific cortical generator.

The second aim was to determine whether context-related differences in canonical theta, alpha, and beta power remained after accounting for trial-level nuisance signals. Noise-layer root-mean-square activity, neck-electromyography root-mean-square activity, and motion/miscellaneous-channel root-mean-square activity were used to model variance that could plausibly be associated with cable movement, muscle activation, and body or equipment motion. Residual effects were not expected to be artifact-free, because nuisance regression can only account for variation represented by the included predictors. However, persistence after residualization would provide stronger evidence of a context-associated scalp spectral difference than an unadjusted comparison alone.

The third aim was to test whether the clearest context differences were expressed primarily in aperiodic spectral structure rather than being explained solely by canonical bandpower or consistently detected periodic peaks. The aperiodic exponent was treated as the principal spectral-parameterization outcome. A higher or lower exponent was not assigned a predetermined psychological interpretation, because the direction of such an effect could plausibly depend on several interacting task properties. The more specific hypothesis was that spectral parameterization would reveal posterior and parietal context effects that could not be described adequately as selective alpha- or beta-peak differences.

The fourth aim was to assess whether the parietal impact-window contrast remained stable under alternative spectral specifications selected on statistical and scientific grounds. The initial short-window specification was compared with a principal resolution-aware model in which the lower peak-width limit was aligned with the frequency grid. Reduced-complexity max3, better-resolved two-second max6, and two-second aperiodic-only analyses then tested dependence on maximum peak count, temporal resolution, and periodic-peak modelling. These analyses evaluated analytic robustness within the same participant sample rather than independent replication. Agreement across specifications would indicate that the focal context difference was not dependent on one peak-width setting, one level of model complexity, or one treatment of periodic spectral components.

## 2 Materials and Methods

### 2.1 Dataset and participants

The analysis was based on the publicly available Real World Table Tennis dataset hosted on OpenNeuro (Studnicki & Ferris, 2023). The present investigation was a secondary analysis of previously collected and deidentified data.

The public processed dataset contained 25 healthy adults who completed real-world table-tennis play with both a human opponent and a ball machine. All 25 participants were retained in the present workflow and contributed usable participant-hit epochs to the final paired analyses. The mean age was 22.0 years, with a standard deviation of 3.8 years and a range of 18–30 years. The source metadata field for sex recorded 15 participants as male and 10 as female. Participants were right-hand dominant, had normal or corrected-to-normal vision, and reported no musculoskeletal or neurological injuries in the original study documentation (Iammarino et al., 2026; Studnicki & Ferris, 2024).

The original experimental protocol included approximately 60 min of table-tennis play, divided between human-opponent and ball-machine conditions, together with standing baseline periods. Human-opponent trials included cooperative and competitive exchanges. Ball-machine trials included stationary and moving deliveries designed to approximate serve-like and rally-like shots. The comparison therefore involved two related forms of object interception rather than a contrast between active and inactive conditions. Both required visual tracking, stroke preparation, racket–ball contact, and continued movement, although they differed in the source and predictability of the incoming ball and in the availability of biological and strategic information.

Recordings were obtained using a high-density dual-layer EEG system. The principal electrophysiological montage comprised 120 scalp EEG electrodes and 120 mechanically matched noise-layer electrodes. Eight additional electrodes recorded activity from the left and right sternocleidomastoid and trapezius muscles. Synchronized inertial and miscellaneous signals were also available from the participant, paddles, table, net, ball machine, amplifier backpack, and associated synchronization streams. Because the set of auxiliary sensors was not identical for every participant, the total number of recorded channels ranged from 270 to 340, although the numbers of scalp, noise-layer, and neck-EMG channels were constant across the main dataset. Studnicki et al. (2022) and Studnicki and Ferris (2024) provide full descriptions of the hardware, task protocol, and organization of the public release.

The raw dual-layer EEG signals were originally acquired at 500 Hz using four BrainVision LiveAmp 64 amplifiers. The synchronized and merged EEGLAB-format files made available for the present analysis had already been resampled to 250 Hz. This distinction is important because 500 Hz describes the original acquisition rate, whereas 250 Hz was the sampling frequency that determined the number of samples, frequency grids, and spectral resolution of the analyses reported here. The scalp and noise-layer signals used in the current pipeline were analyzed between 1 and 45 Hz. The dataset was organized according to the Brain Imaging Data Structure (BIDS), which provides standardized conventions for organizing neuroimaging data and associated metadata (Gorgolewski et al., 2016).

The original data collection was approved by the University of Florida Institutional Review Board, and written informed consent was obtained from the participants by the original investigators.

### 2.2 Channel classification and preprocessing

Preprocessing was carried out using a custom Python workflow built around MNE-Python, which provides tools for importing, filtering, referencing, epoching, and analysing electrophysiological recordings (Gramfort, 2013). The EEGLAB .set and associated .fdt files were loaded separately for each participant. Channel identities were then classified before any operation that depended on channel type. This step was necessary because the merged files contained scalp EEG, noise-layer EEG, neck electromyography, accelerometry, inertial measurements, synchronization streams, and signals associated with the paddles and table-tennis equipment.

Channels whose labels began with the N-prefix were classified as noise-layer measurements. Channels corresponding to the left and right sternocleidomastoid and trapezius muscles were classified as EMG. Labels containing terms associated with accelerometers, inertial measurement units, gyroscopes, magnetometers, synchronization pulses, paddles, the ball machine, backpack, net, table, or Cometa recording streams were classified as motion or miscellaneous channels. All remaining electrophysiological channels were treated as scalp EEG. This rule-based classification was checked against participant-level channel summaries so that auxiliary channels were not inadvertently included in the neural scalp montage.

After classification, the 120 scalp channels were assigned the MNE channel type eeg, the eight neck-muscle channels were assigned emg, and the noise-layer and motion/miscellaneous channels were treated as auxiliary measurements. The noise-layer electrodes were therefore not interpreted as additional scalp EEG channels or included as neural outcomes. Their role was to provide reference information concerning mechanically and environmentally induced signal variation. In the dual-layer system, each electrically isolated noise electrode was mechanically coupled to a corresponding scalp electrode so that cable movement, electrode movement, and other non-biological perturbations could be represented without recording the scalp potential measured by the paired EEG electrode. Previous analysis of these recordings showed that matched noise-layer signals captured artifact structure that was more closely related to scalp-channel contamination than were general measures of head and body acceleration, supporting their use as nuisance or reference measurements during mobile EEG analysis (Studnicki et al., 2022).

Discontinuities in the EEGLAB recordings were handled before filtering and epoch creation. Annotations whose descriptions identified boundary events were converted into BAD_boundary segments, with two seconds of padding added before and after each boundary. This conservative interval was intended to prevent filtering transients or discontinuity-related spectral structure from entering event-locked epochs. Epochs that overlapped these annotations were subsequently rejected automatically.

Scalp EEG and noise-layer channels were filtered between 1 and 45 Hz. Applying the same frequency range to the scalp and noise-layer recordings allowed nuisance signals to be represented within the range used for the principal scalp spectral analyses. The neck-EMG and motion/miscellaneous channels were preserved as auxiliary signals rather than subjected to the same scalp-EEG processing sequence, because their purpose was to provide nuisance information rather than neural bandpower outcomes. Following filtering, an average reference was applied to the 120 scalp EEG channels only. Noise-layer, EMG, and motion channels were excluded from calculation of the scalp reference.

No additional independent-component rejection was performed within this secondary workflow. This decision should not be interpreted as implying that ICA is unsuitable for mobile EEG. The original dataset includes ICA-related materials, and the earlier dual-layer investigation demonstrated advantages of incorporating noise-channel information into component-based artifact-removal procedures (Studnicki et al., 2022). However, the present analysis followed a deliberately conservative sensor-level approach in which scalp signals were retained and trial-level nuisance information was modelled explicitly. Avoiding additional component rejection also reduced the possibility that differences between conditions could be introduced by subjective or condition-dependent component-selection decisions. It does not, however, imply that the resulting signals were artifact-free.

### 2.3 Event construction and epoching

Event information was read from the participant-specific BIDS event files and reorganized into a common trial-level table. The initial event table was restricted to participant-centered events labelled Subject_hit and Subject_receive, because these events represented the principal perception–action sequence experienced by the participant. Non-task, synchronization, and status-like rows were excluded from the event analysis. Cooperative and competitive table-tennis trials were mapped to the human-opponent context. Moving-serve, stationary-serve, moving-hit, and stationary-hit trials were mapped to the ball-machine context. The term human-opponent is used throughout the manuscript to provide a consistent condition label, even though the cooperative trials did not necessarily involve competitive opposition. It denotes that the ball was returned by another person rather than generated by the machine.

The main event-locked analysis was based on participant racket–ball contacts, corresponding to the source event label Subject_hit. For each participant, eligible participant-hit rows were retained only when they could be assigned unambiguously to the human-opponent or ball-machine context. Events were ordered by their sample index, rows with negative sample values were removed, and duplicate sample events were removed when present. The remaining events were converted to MNE event arrays with separate numerical event codes for human-opponent and ball-machine hits. Participant identifier, trial identifier, onset, sample index, task context, original trial type, event role, adjacent event roles, and timing relative to neighbouring events were retained in the epoch metadata.

Continuous data were segmented from −1.5 to +1.0 s relative to each participant hit. No baseline correction was applied. Baseline subtraction can alter the apparent relationship between conditions when the baseline itself contains context-related activity, and the present objective was to characterize the spectral structure within explicitly defined peri-hit intervals rather than relative change from a separate reference period. Annotation-based rejection was enabled so that any epoch overlapping a padded BAD_boundary segment was excluded.

Three shorter intervals were extracted from each retained epoch. The pre-hit interval extended from −1.0 to −0.2 s, the impact interval from −0.2 to +0.3 s, and the post-hit interval from +0.3 to +1.0 s. These labels provide operational descriptions of time relative to racket–ball contact. They should not be interpreted as pure or mutually exclusive cognitive stages, because visual prediction, movement preparation, muscular activity, sensory feedback, and preparation for the subsequent exchange may overlap across the intervals.

MNE-Python includes both endpoints when an epoch is cropped by default. At the 250-Hz sampling frequency, the pre-hit interval therefore contained 201 samples, the impact interval contained 126 samples, and the post-hit interval contained 176 samples. The pre-hit and impact intervals shared the sample at −0.2 s, while the impact and post-hit intervals shared the sample at +0.3 s. Each shared sample represented 4 ms of data. The intervals were consequently not mathematically disjoint, although the overlap was restricted to a single boundary sample at each transition.

A broader two-second peri-hit interval, extending from −1.0 to +1.0 s, was also extracted for the better-resolved spectral sensitivity analyses. Because both endpoints were included, this interval contained 501 samples. The two-second interval averaged across the pre-hit, impact, and post-hit periods and was not used as a replacement for the temporally specific short-window analyses.

### 2.4 Sensor grouping, bandpower extraction, and nuisance residualization

Scalp channels were assigned to six broad sensor groups using a rule-based mapping of their channel labels. Frontal and anterior-frontal labels were assigned to the frontal group. Central and frontocentral labels were assigned to the sensorimotor group. Parietal and centroparietal labels were assigned to the parietal group, while occipital and parieto-occipital labels were assigned to the posterior group. Temporal and frontotemporal labels were assigned to the temporal group. Channels that could not be assigned clearly to these categories were retained in another group rather than being discarded.

These groupings were intended to provide interpretable summaries of the high-density scalp recordings. They should not be treated as equivalent to source-localized cortical regions. Scalp potentials are affected by volume conduction, referencing, sensor placement, and the spatial mixing of neural and non-neural sources. The terms frontal, parietal, posterior, temporal, and sensorimotor therefore refer to sensor distributions rather than direct anatomical localization.

Canonical bandpower was estimated for theta, alpha, and beta activity. Theta was defined as 4–7 Hz, alpha as 8–13 Hz, and beta as 14–30 Hz. Frequencies above 30 Hz were not included as canonical outcomes because high-frequency scalp power recorded during whole-body movement is particularly susceptible to myogenic contamination.

For each retained epoch, the scalp data were cropped separately to the pre-hit, impact, and post-hit intervals. Power spectral density was estimated between 2 and 40 Hz using Welch’s method. Welch estimation divides a signal into windowed segments, estimates a modified periodogram for each segment, and averages the resulting estimates to reduce periodogram variance (Welch, 1967). In the bandpower pipeline, the FFT length was set to the smaller of 128 samples or the number of available samples in the cropped interval. The Welch segment length equalled the FFT length, and adjacent segments overlapped by 50%. A Hamming window was used through the MNE-Python Welch implementation.

Power values were transformed using the base-10 logarithm before aggregation. For each epoch, time interval, band, and sensor region, mean log power was calculated across the frequency bins within the relevant canonical band and across the scalp channels assigned to the region. This produced one trial-level scalp-bandpower value for every available window *×* band *×* region combination.

Three trial-level nuisance measures were then calculated for each event window. Noise-layer root mean square (RMS) was calculated across the matched noise-layer channels and time points. EMG RMS was calculated across the eight neck-muscle channels and time points. Motion/miscellaneous RMS was calculated from the available auxiliary movement and miscellaneous channels. These measures were intended to summarize broad trial-to-trial variation in cable or electrode movement, neck-muscle activity, and body or equipment motion. They were not interpreted as experimental outcomes.

Residualization was performed before participant-level condition means were calculated. For each window *×* band *×* sensor-region combination, trial-level log bandpower was entered as the dependent variable in an ordinary least-squares regression. Standardized noise-layer RMS, EMG RMS, and motion/miscellaneous RMS were entered simultaneously as nuisance predictors. Experimental context was deliberately excluded from the nuisance model because the human-opponent versus ball-machine difference was the effect subsequently tested. Including context as a nuisance regressor would have removed part of the variation that the analysis was intended to estimate.

The residual from each nuisance model was retained as the nuisance-residualized bandpower value for that trial. Residuals were then averaged separately for each participant and context. The human-opponent and ball-machine values were consequently compared as paired participant-level observations rather than as independent trials. This aggregation prevented the large number of event epochs from being treated as the effective inferential sample and ensured that participants with more recorded hits did not contribute additional independent degrees of freedom.

Residualization cannot demonstrate that the remaining scalp differences were entirely neural. Linear regression can account only for variation represented by the included nuisance features, and nonlinear or unmeasured artifact contributions may remain. The procedure instead provides a more conservative comparison than unadjusted bandpower by asking whether context-related differences remain after variance associated with directly recorded nuisance signals has been removed. The resulting outcomes are therefore referred to as residualized bandpower during human and machine play, not as aperiodic-adjusted power. No aperiodic exponent, offset, or fitted aperiodic spectrum was included in the bandpower residualization model.

### 2.5 Power-spectral estimation and spectral resolution

A separate spectral-estimation procedure was used to generate the spectra entered into the spectral-parameterization models. For each participant, context, event window, and sensor region, linear-power Welch spectra were estimated from the retained scalp epochs. Spectra were averaged across the epochs belonging to the same context and across the channels assigned to the same sensor region before spectral parameterization. Each fitted model therefore represented one participant *×* context *×* window *×* region spectrum rather than one model for every individual trial.

The spectra supplied to the spectral-parameterization model remained in linear-power units. The implementation expects frequency values and power spectra in linear space and applies the transformations required during fitting. Supplying spectra that had already been logarithmically transformed would not reproduce the intended model.

Short-window spectra were estimated using a Hamming window, with both the FFT length and Welch segment length set to the complete number of samples in each cropped interval. No additional zero-padding was used. The pre-hit spectrum used a 201-sample FFT and segment with 100-sample overlap; the impact spectrum used 126 samples with 63-sample overlap; and the post-hit spectrum used 176 samples with 88-sample overlap. The corresponding frequency-bin spacings, calculated as the sampling frequency divided by the FFT length, were 1.2438 Hz, 1.9841 Hz, and 1.4205 Hz, respectively.

For the two-second peri-hit sensitivity models, the FFT length and segment length were both 501 samples. Because the segment occupied the complete two-second window, the overlap was set to zero. This produced a frequency-bin spacing of 0.4990 Hz. In all spectral-parameterization analyses, models were fitted between 3 and 40 Hz.

### 2.6 Spectral-parameterization model hierarchy

Spectral parameterization was performed with specparam version 2.0.0rc7. The model represents each power spectrum as an aperiodic component together with zero or more periodic peaks (Donoghue et al., 2020). The aperiodic component was fitted in fixed mode. Unless periodic fitting was disabled, the upper peak-width limit was 8 Hz, the peak threshold was two standard deviations above the aperiodic-removed spectrum, and the minimum peak height was 0.05. The allowable peak-width range determined the periodic bandwidths that could be fitted, while the peak-count setting determined the maximum number of periodic components.

The initial short-window specification used fixed peak-width limits of 1–8 Hz, allowed a maximum of six periodic peaks, and was fitted separately to the pre-hit, impact, and post-hit spectra. A formal audit of the window-specific frequency grids showed that the lower peak-width limit was smaller than the available short-window frequency resolution, particularly for the impact interval, which had a frequency spacing of 1.9841 Hz. Because periodic components narrower than the spectral grid cannot be represented reliably, the principal short-window analysis used resolution-aware lower peak-width limits. The initial specification was retained as a sensitivity comparator so that the stability of the context effect could be evaluated against the scientifically motivated change in model settings.

The resolution-aware max6 model was used as the principal short-window analysis. Its lower peak-width limit was set to approximately twice the frequency-bin spacing of each event window. The resulting minimum widths were 2.488 Hz for pre-hit spectra, 3.968 Hz for impact spectra, and 2.841 Hz for post-hit spectra. The upper width limit remained 8 Hz, and a maximum of six peaks was permitted. This specification retained the flexibility of the initial specification while preventing the fitting of periodic components that were narrower than could be meaningfully represented by the spectral grid.

The resolution-aware max3 model used the same window-specific peak-width limits but reduced the maximum number of periodic peaks from six to three. This model was included as a reduced-complexity sensitivity analysis. Agreement between max6 and max3 would indicate that the focal aperiodic effect was not dependent on allowing a relatively large number of periodic components. Differences in model complexity were also examined using residual-based Bayesian information criterion summaries.

The two-second peri-hit max6 model was fitted from −1.0 to +1.0 s around participant contact. Its 0.4990-Hz frequency spacing permitted a lower peak-width limit of 0.998 Hz, with an upper limit of 8 Hz and a maximum of six peaks. This model provided a better-resolved sensitivity analysis of the broad peri-hit spectrum. It did not estimate the same temporal quantity as the short impact-window model because it averaged activity across the pre-hit, impact, and post-hit intervals.

The two-second aperiodic-only model used the same peri-hit interval, frequency grid, and fitting range, but periodic fitting was disabled by setting the maximum number of peaks to zero. This model was intended as an aperiodic-only sensitivity analysis rather than as the preferred overall representation of the spectrum. Its purpose was to determine whether a comparable participant-level context difference remained when no spectral variation could be assigned to periodic peaks. A similar condition effect under max6 and aperiodic-only specifications would indicate that the result was not contingent on one particular treatment of oscillatory activity, although the aperiodic-only model would be expected to provide a poorer description of spectra that contained genuine periodic peaks.

The principal fitted parameters were the aperiodic exponent and offset. The exponent was designated as the primary spectral-parameterization outcome because it described broadband spectral slope after the fitted periodic components had been considered. The offset was retained as a secondary aperiodic outcome. Periodic alpha and beta peaks were recorded when present, including center frequency, power, and bandwidth, but these outcomes were treated as model-dependent diagnostics because peak detection varied with window duration, frequency resolution, and model complexity. Model R^2^, model error, and peak-detection rates were similarly treated as quality-control measures rather than as interchangeable observations of neural effect magnitude.

### 2.7 Statistical analysis

All inferential analyses were conducted at the participant level. Each participant contributed one human-opponent and one ball-machine estimate for a given feature, window, and sensor region. Condition differences were consistently calculated as ball machine minus human opponent, meaning that positive aperiodic-exponent values indicated a steeper exponent during ball-machine play, whereas negative residualized-bandpower values indicated lower residualized power during ball-machine play.

The aperiodic exponent from the resolution-aware max6 model formed the primary short-window inferential family. This family contained 18 comparisons, comprising three temporal windows and six sensor regions. A two-sided paired-samples t test and a two-sided Wilcoxon signed-rank test were calculated for every comparison. The paired t test estimated the mean participant-level difference under a parametric model, while the Wilcoxon test provided a complementary rank-based analysis that was less dependent on normality of the paired differences.

The 18 paired-t probabilities were corrected together using the Benjamini–Hochberg false-discovery-rate procedure. The 18 Wilcoxon probabilities were corrected as a separate family. Adjusted probabilities are reported as FDR q values. The aperiodic offset was treated as a secondary feature and corrected within its own 18-comparison paired-t and Wilcoxon families. The Benjamini–Hochberg procedure was selected to limit the expected proportion of false discoveries among rejected hypotheses while retaining greater sensitivity than familywise-error procedures in a fixed multicomparison setting (Benjamini & Hochberg, 1995).

The residualized-bandpower family contained 54 comparisons, corresponding to three temporal windows, three frequency bands, and six sensor regions. The 54 paired-t probabilities were corrected as one family, and the 54 Wilcoxon probabilities were corrected separately. This family definition was fixed by the complete window *×* band *×* region structure and was not limited to the comparisons that produced the smallest unadjusted probabilities.

Cohen’s dz was calculated for each participant-level paired comparison by dividing the mean of the paired differences by the standard deviation of those differences. This form of standardized effect size was used because observations from the two contexts were not independent. The sign of dz followed the ball-machine-minus-human-opponent contrast. Effect sizes were interpreted together with the observed difference, uncertainty interval, inferential statistics, and participant-level direction rather than as isolated labels such as small, medium, or large (Lakens, 2013).

Direction consistency was calculated by counting the participants whose difference had the same sign as the group mean. The number and percentage showing the machine-greater-than-human or machine-lower-than-human direction were reported alongside the paired tests. This measure was included because a statistically detectable mean difference may still be difficult to interpret when it is produced by only a small subset of participants.

The impact-window parietal contrast was selected as the focal robustness target because it combined an anatomically interpretable sensor region with the short interval having the coarsest spectral grid. This choice provided a stringent test of whether the context effect depended on model settings under the most demanding short-window resolution condition. The same participant-level contrast was evaluated under the initial short-window specification, the resolution-aware max6 model, and the resolution-aware max3 model. The complete model hierarchy was not prospectively preregistered; the additional specifications are therefore treated as sensitivity analyses within the same dataset rather than as independent confirmatory tests (Nosek et al., 2018).

Uncertainty around the focal mean difference and Cohen’s dz was assessed with 50,000 paired bootstrap resamples. The participant-level paired differences were sampled with replacement, and each resample retained the pairing of the original condition estimates. Confidence limits were obtained from the empirical bootstrap distribution. Bootstrap resampling provides a way to characterize sampling uncertainty without relying entirely on the analytic normal approximation for the effect estimate (Efron & Tibshirani, 1993).

A leave-one-participant-out analysis was also performed. Each participant was omitted in turn, after which the focal condition difference, paired tests, effect size, direction count, and relevant FDR-adjusted values were recalculated using the remaining 24 participants. Robustness was summarized as the number of omitted-participant analyses that retained a positive machine-minus-human difference and the number that retained FDR significance. This procedure was intended to determine whether the focal result depended disproportionately on one participant.

Participant-level concordance between spectral models was assessed using Pearson correlation, Spearman rank correlation, the mean and median difference between model-specific effects, mean absolute effect difference, maximum absolute effect difference, and the number of participants whose effect direction changed. These measures addressed related but distinct questions. Correlation assessed whether participants were ordered similarly across models, whereas absolute-difference and direction-agreement measures tested whether the numerical effects themselves remained comparable.

The relative complexity of the resolution-aware max6 and max3 models was examined using the Bayesian information criterion (BIC). Lower BIC values were interpreted as indicating the better balance between fit and model complexity. Differences were calculated as max3 BIC minus max6 BIC, so negative values favored max3 and positive values favored max6. BIC summaries were evaluated across model cells and for the focal parietal comparison. These summaries were treated as model diagnostics rather than as participant-level hypothesis tests because many cells contained identical or near-identical fitted structures (Schwarz, 1978).

The two-second max6 result was additionally examined for possible dependence on condition differences in spectral-model quality. For each participant, the ball-machine-minus-human difference in aperiodic exponent was related to the corresponding difference in R^2^ and model error. Spearman correlations were accompanied by permutation probabilities. Ordinary least-squares regressions were estimated with HC3 heteroskedasticity-consistent intervals, and Theil–Sen estimates were included as a robust comparison. Cook’s distance was calculated to identify observations with disproportionate regression influence, using 4/n as the screening threshold. These analyses were considered exploratory because aperiodic exponent and fit quality were outputs of the same spectral model and were therefore not statistically independent constructs (Cook, 1977; MacKinnon & White, 1985).

A quality-neutral intercept analysis was used to estimate the expected context difference when the modelled between-condition difference in R^2^ or error equalled zero. The intercept was accompanied by an HC3 confidence interval and a separate 50,000-resample bootstrap interval. A positive intercept under equal modelled fit quality would be consistent with the effect not being explained entirely by one context receiving a systematically better spectral fit. It would not, however, prove the absence of all signal-quality influence.

Finally, the two-second max6 and two-second aperiodic-only estimates were compared using paired inferential tests, bootstrap intervals, leave-one-participant-out analyses, participant-level correlations, mean absolute differences, and effect-direction agreement. These analyses tested whether the broad peri-hit context difference depended on assigning spectral structure to periodic peaks. The short impact-window and two-second peri-hit estimates were not treated as repeated estimates of an identical parameter. The former was specific to the 0.5-s interval around contact, whereas the latter averaged across a two-second period containing anticipatory, impact-related, and post-hit activity.

All tests were two-sided unless explicitly stated otherwise. Statistical interpretation considered the observed condition difference, Cohen’s dz, bootstrap interval, adjusted q values, participant-level direction consistency, and sensitivity across model specifications. No result was interpreted solely because an adjusted probability fell below .05.

### 2.8 Software

The main analysis used Python-based scientific computing tools. MNE-Python was used for loading, preprocessing, epoching, and spectral estimation of EEG (Gramfort, 2013). NumPy and pandas were used for numerical operations and tabular data handling (Harris et al., 2020; McKinney, 2010). SciPy was used for paired inferential statistics (Virtanen et al., 2020), and statsmodels was used for ordinary least-squares residualization and false discovery rate correction workflows (Seabold & Perktold, 2010). Spectral parameterization was performed using specparam, with compatibility retained for legacy FOOOF-style implementations where needed (Donoghue, Haller, et al., 2020). Matplotlib was used for visualization (Hunter, 2007). The analysis code is available at https://gitlab.com/brain-behav-lab/tabletennis-eeg-analysis.

## 3 Results

### 3.1 Dataset retention and participant-level balance

All 25 participants in the public dataset were successfully preprocessed and contributed usable participant-hit epochs to the final analysis. A total of 37,556 candidate participant racket-ball contact events were identified. Following rejection of epochs overlapping the padded boundary annotations, 36,811 epochs were retained, corresponding to an overall retention rate of 98.02%. Participant-level retention was consistently high, ranging from 97.7% to 98.3%. The number of retained epochs per participant ranged from 1,072 to 2,220, indicating that all participants contributed substantial numbers of events to the participant-level condition estimates.

The retained dataset contained 19,212 human-opponent epochs and 17,599 ball-machine epochs. These values represented 52.19% and 47.81% of the complete retained sample, respectively. Every participant contributed epochs from both contexts, although the relative condition balance differed across participants. Human-opponent epochs accounted for between 46.48% and 61.38% of the retained epochs within an individual participant, while ball-machine epochs accounted for between 38.62% and 53.52%. The inferential analyses were nevertheless based on one aggregated estimate per participant and condition, meaning that differences in the number of retained trials did not alter the participant-level degrees of freedom.

Table 1 summarizes the dataset, preprocessing, event-locking, and epoch-retention outcomes. The participant-specific condition counts and retention percentages are shown in Supplementary Figure S1 and reported in full in Supplementary Table S1.

**Table 1.**
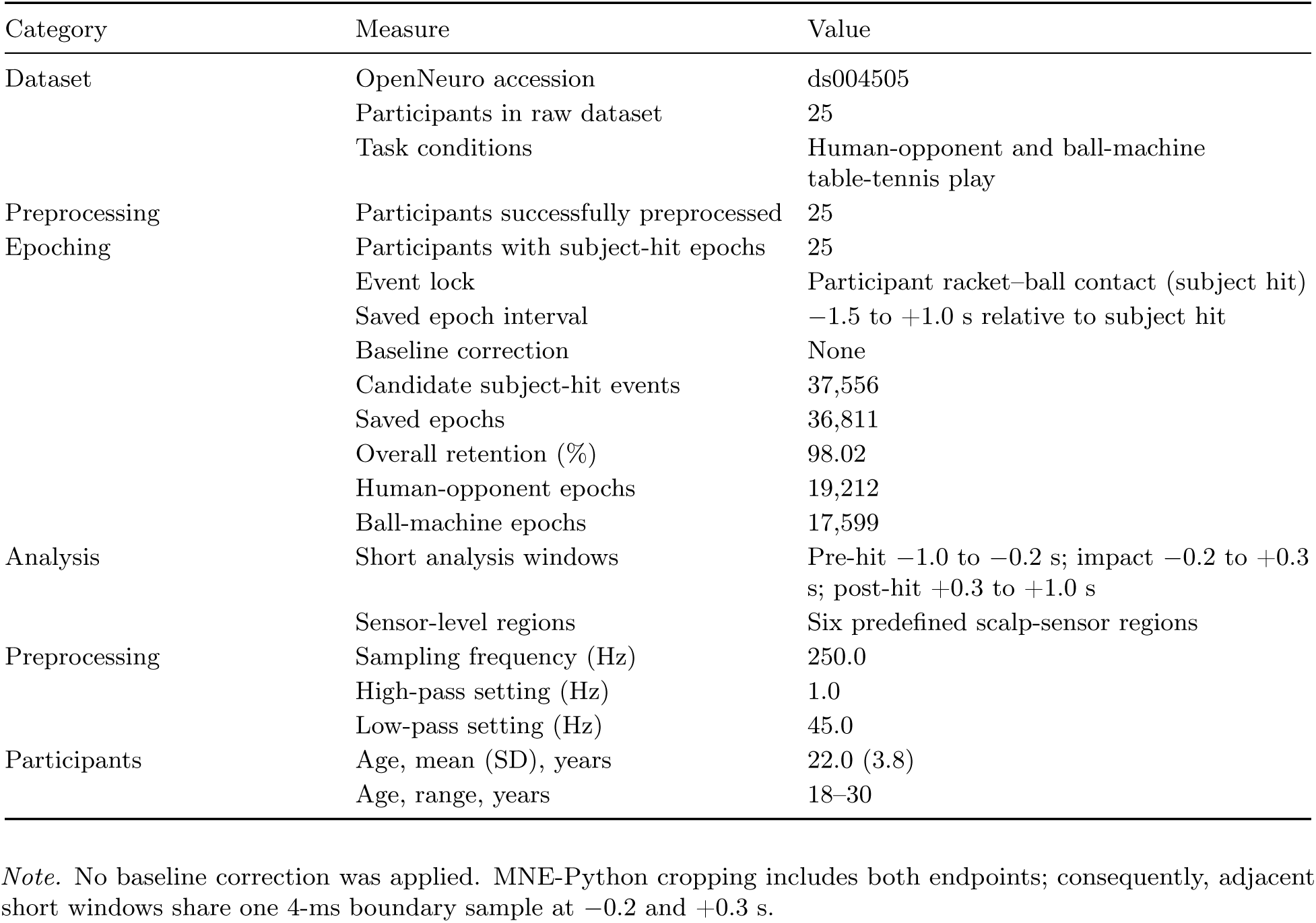
Dataset, preprocessing, and epoch-retention summary. Summary of the OpenNeuro dataset, participant characteristics, preprocessing, subject-hit epoching, condition counts, and sensor-level analysis structure.

### 3.2 Primary short-window aperiodic-exponent results

The principal resolution-aware max6 model produced a consistent spatial pattern in which the aperiodic exponent was generally higher during ball-machine than human-opponent play over posterior, parietal, temporal, and other sensors, whereas frontal and sensorimotor differences were smaller and did not survive correction. Of the 18 fixed short-window comparisons, 11 survived both the paired-t and Wilcoxon FDR corrections. Adjusted values reported below are Benjamini-Hochberg FDR q values (Benjamini & Hochberg, 1995). The complete regional and temporal pattern is illustrated in Figure 1.

**Figure 1.**
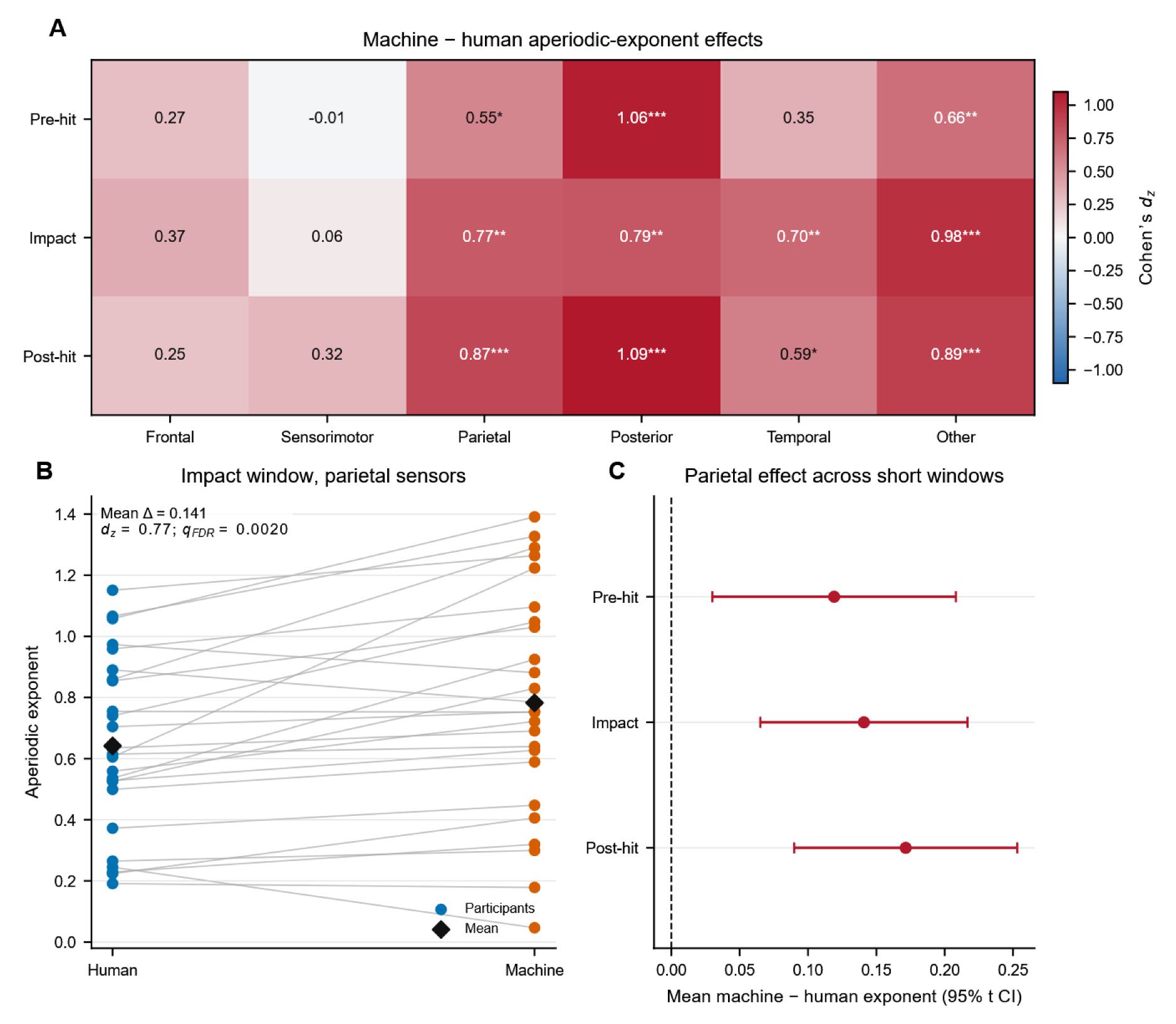
Spatiotemporal differences in the aperiodic exponent between ball-machine and human-opponent play. A, Cohen’s dz for all 18 short-window comparisons from the preferred resolution-aware max6 model. Positive values indicate a steeper aperiodic exponent during ball-machine play. Asterisks indicate Benjamini–Hochberg-adjusted paired t-test values within the fixed 18-test exponent family (*q < 0.05, **q < 0.01, ***q < 0.001). B, paired participant-level aperiodic-exponent values for the impact window over parietal sensors; connecting lines link the two conditions within a participant and diamonds show condition means. The inset reports the mean machine-minus-human difference, Cohen’s dz, and FDR-adjusted paired t-test value. C, mean parietal machine-minus-human differences across the three short windows with paired t-based 95% confidence intervals. n = 25 participants.

#### 3.2.1 Pre-hit interval

During the pre-hit interval, the clearest difference was present over the posterior sensor region. The mean aperiodic exponent was 0.469 during human-opponent play and 0.602 during ball-machine play, giving a mean ball-machine-minus-human-opponent difference of 0.133. The paired comparison produced t(24) = 5.30, p < .001, paired-t FDR q < .001, Cohen’s dz = 1.060, and Wilcoxon p < .001, FDR q < .001. The exponent was higher during ball-machine play in 21 of 25 participants, corresponding to 84.0% direction consistency.

A smaller but still significant when corrected difference was present over the parietal region. The human-opponent and ball-machine means were 0.639 and 0.758, respectively, with a mean difference of 0.119, t(24) = 2.76, p = .011, paired-t FDR q = .018, dz = 0.552, Wilcoxon p = .020, and Wilcoxon FDR q = .033. Nineteen participants, or 76.0% of the sample, showed a higher exponent during ball-machine play. The other sensor category also showed a positive difference, with human-opponent and ball-machine means of 0.649 and 0.755, a mean difference of 0.106, t(24) = 3.32, p = .003, paired-t FDR q = .006, dz = 0.664, Wilcoxon p = .002, and Wilcoxon FDR q = .004. The machine-higher direction was observed in 19 participants.

The frontal, sensorimotor, and temporal pre-hit comparisons did not survive correction. The frontal difference was 0.044, t(24) = 1.37, paired-t FDR q = .220, Wilcoxon FDR q = .218, and dz = 0.274. The sensorimotor difference was -0.003, t(24) = -0.07, paired-t FDR q = .947, Wilcoxon FDR q = .958, and dz = -0.013. The temporal difference was 0.064, t(24) = 1.73, paired-t FDR q = .133, Wilcoxon FDR q = .163, and dz = 0.346. The pre-hit effect was therefore most clearly expressed over posterior sensors, with additional corrected differences over parietal and other sensors rather than as a uniform scalp-wide increase.

#### 3.2.2 Impact interval

During the impact interval, the focal parietal robustness target showed a human-opponent mean of 0.642 and a ball-machine mean of 0.782. The mean difference was 0.141, t(24) = 3.84, p < .001, paired-t FDR q = .002, dz = 0.768, Wilcoxon p < .001, and Wilcoxon FDR q = .002. Twenty of 25 participants, or 80.0%, showed the ball-machine-greater-than-human-opponent direction.

The posterior impact comparison produced a human-opponent mean of 0.484 and a ball-machine mean of 0.597, giving a difference of 0.113, t(24) = 3.94, p < .001, paired-t FDR q = .002, dz = 0.788, Wilcoxon p < .001, and Wilcoxon FDR q < .001. The exponent was higher during ball-machine play in 21 participants. The temporal impact effect was similar in magnitude, with human-opponent and ball-machine means of 0.320 and 0.451, a mean difference of 0.131, t(24) = 3.51, p = .002, paired-t FDR q = .004, dz = 0.702, Wilcoxon p = .002, and Wilcoxon FDR q = .004. Twenty participants showed the positive direction.

The numerically largest impact difference was found in the other sensor category, where the means were 0.622 and 0.776 for human-opponent and ball-machine play, respectively. The mean difference was 0.154, t(24) = 4.88, p < .001, paired-t FDR q < .001, dz = 0.975, Wilcoxon p < .001, and Wilcoxon FDR q < .001. Twenty-two participants, or 88.0%, showed a higher exponent during ball-machine play. Because this sensor category did not correspond to a single anatomically defined distribution, it was retained in the full statistical family but was not treated as the principal neuroanatomical result.

The frontal and sensorimotor impact comparisons did not survive either FDR family. The frontal difference was 0.067, t(24) = 1.83, paired-t FDR q = .119, Wilcoxon FDR q = .100, and dz = 0.366. The sensorimotor difference was 0.014, t(24) = 0.32, paired-t FDR q = .795, Wilcoxon FDR q = .225, and dz = 0.064.

#### 3.2.3 Post-hit interval

The largest anatomically interpretable post-hit effect was present over the posterior region. The human-opponent mean was 0.451 and the ball-machine mean was 0.606, producing a mean difference of 0.155. The paired comparison gave t(24) = 5.46, p < .001, paired-t FDR q < .001, dz = 1.093, Wilcoxon p < .001, and Wilcoxon FDR q < .001. Twenty-three of the 25 participants, corresponding to 92.0% of the sample, showed a higher posterior exponent during ball-machine play.

The parietal post-hit difference was numerically slightly larger, although it had a somewhat smaller standardized effect because the participant differences were more variable. The human-opponent and ball-machine means were 0.615 and 0.786, respectively, giving a mean difference of 0.171, t(24) = 4.34, p < .001, paired-t FDR q < .001, dz = 0.867, Wilcoxon p < .001, and Wilcoxon FDR q < .001. Twenty-one participants, or 84.0%, showed the positive direction.

A corrected temporal post-hit effect was also present. The human-opponent mean was 0.265, the ball-machine mean was 0.374, and the resulting difference was 0.109, t(24) = 2.95, p = .007, paired-t FDR q = .013, dz = 0.589, Wilcoxon p = .011, and Wilcoxon FDR q = .019. Eighteen participants showed a higher exponent during machine play. The other sensor category produced a difference of 0.127, t(24) = 4.47, p < .001, paired-t FDR q < .001, dz = 0.894, Wilcoxon p < .001, and Wilcoxon FDR q < .001, with the positive direction present in 22 participants.

No corrected post-hit difference was detected over frontal or sensorimotor sensors. The frontal difference was 0.041, t(24) = 1.25, paired-t FDR q = .252, Wilcoxon FDR q = .331, and dz = 0.250. The sensorimotor difference was 0.043, t(24) = 1.59, paired-t FDR q = .161, Wilcoxon FDR q = .111, and dz = 0.318.

Taken together, the short-window results showed that the machine-higher aperiodic-exponent pattern was strongest and most consistent over posterior and parietal sensors. Posterior effects were present before, during, and after participant contact, with dz values ranging from 0.788 to 1.093 and the positive direction present in 84.0-92.0% of participants. Parietal effects were also present across all three intervals, although the pre-hit effect was smaller than the impact and post-hit effects. The full 18-comparison family, including all null comparisons and both FDR-adjusted inferential tests, is summarized in Table 2. Complete participant distributions are shown in Supplementary Figure S2, and the corresponding exponent and offset statistics are reported in Supplementary Table S3.

**Table 2.** Primary aperiodic-exponent results. All 18 short-window comparisons from the preferred resolution-aware max6 spectral model.

| Window | Sensor region | Participants | Human mean | Machine mean | Machine –<br>human mean | Machine –<br>human median | Cohen’s dz | Paired t FDR q | Wilcoxon FDR q | Machine higher, n<br>(%) |
| --- | --- | --- | --- | --- | --- | --- | --- | --- | --- | --- |
| Pre-hit | Frontal | 25 | 1.285 | 1.329 | 0.044 | 0.072 | 0.274 | 0.220 | 0.218 | 14 (56.0) |
|  | Sensorimotor | 25 | 0.895 | 0.892 | -0.003 | 0.019 | -0.013 | 0.947 | 0.958 | 15 (60.0) |
|  | Parietal | 25 | 0.639 | 0.758 | 0.119 | 0.090 | 0.552 | 0.018 | 0.033 | 19 (76.0) |
|  | Posterior | 25 | 0.469 | 0.602 | 0.133 | 0.122 | 1.060 | <0.001 | <0.001 | 21 (84.0) |
|  | Temporal | 25 | 0.332 | 0.396 | 0.064 | 0.072 | 0.346 | 0.133 | 0.163 | 15 (60.0) |
|  | Other | 25 | 0.649 | 0.755 | 0.106 | 0.110 | 0.664 | 0.006 | 0.004 | 19 (76.0) |
| Impact | Frontal | 25 | 1.230 | 1.298 | 0.067 | 0.056 | 0.366 | 0.119 | 0.100 | 17 (68.0) |
|  | Sensorimotor | 25 | 0.862 | 0.876 | 0.014 | 0.032 | 0.064 | 0.795 | 0.225 | 16 (64.0) |
|  | Parietal | 25 | 0.642 | 0.782 | 0.141 | 0.099 | 0.768 | 0.002 | 0.002 | 20 (80.0) |
|  | Posterior | 25 | 0.484 | 0.597 | 0.113 | 0.096 | 0.788 | 0.002 | <0.001 | 21 (84.0) |
|  | Temporal | 25 | 0.320 | 0.451 | 0.131 | 0.091 | 0.702 | 0.004 | 0.004 | 20 (80.0) |
|  | Other | 25 | 0.622 | 0.776 | 0.154 | 0.125 | 0.975 | <0.001 | <0.001 | 22 (88.0) |
| Post-hit | Frontal | 25 | 1.235 | 1.276 | 0.041 | 0.037 | 0.250 | 0.252 | 0.331 | 14 (56.0) |
|  | Sensorimotor | 25 | 0.859 | 0.902 | 0.042 | 0.048 | 0.318 | 0.161 | 0.111 | 15 (60.0) |
|  | Parietal | 25 | 0.615 | 0.786 | 0.171 | 0.134 | 0.867 | <0.001 | <0.001 | 21 (84.0) |
|  | Posterior | 25 | 0.451 | 0.606 | 0.155 | 0.131 | 1.093 | <0.001 | <0.001 | 23 (92.0) |
|  | Temporal | 25 | 0.265 | 0.374 | 0.109 | 0.112 | 0.589 | 0.013 | 0.019 | 18 (72.0) |
|  | Other | 25 | 0.623 | 0.750 | 0.127 | 0.133 | 0.894 | <0.001 | <0.001 | 22 (88.0) |

*Note.* Values are participant-level estimates. Machine – human values greater than zero indicate a steeper aperiodic exponent during ball-machine than human-opponent play. Benjamini–Hochberg FDR correction was applied separately to the fixed 18-test paired t-test and Wilcoxon families. Cohen’s dz is the mean paired difference divided by the SD of paired differences; q denotes an FDR-adjusted p value.

### 3.3 Residualized bandpower results

The nuisance-residualized bandpower analysis showed a broad machine-lower-than-human-opponent pattern. All 54 machine-minus-human-opponent mean differences were negative. Forty-six of the 54 comparisons survived FDR correction in both the paired-t and Wilcoxon families. Two additional comparisons survived only the paired-t correction, leaving six comparisons that did not survive either correction. All 18 beta comparisons survived both FDR procedures, as did 17 of the 18 alpha comparisons and 11 of the 18 theta comparisons. The spatial and temporal distribution of the standardized effects is illustrated in Figure 2.

**Figure 2.**
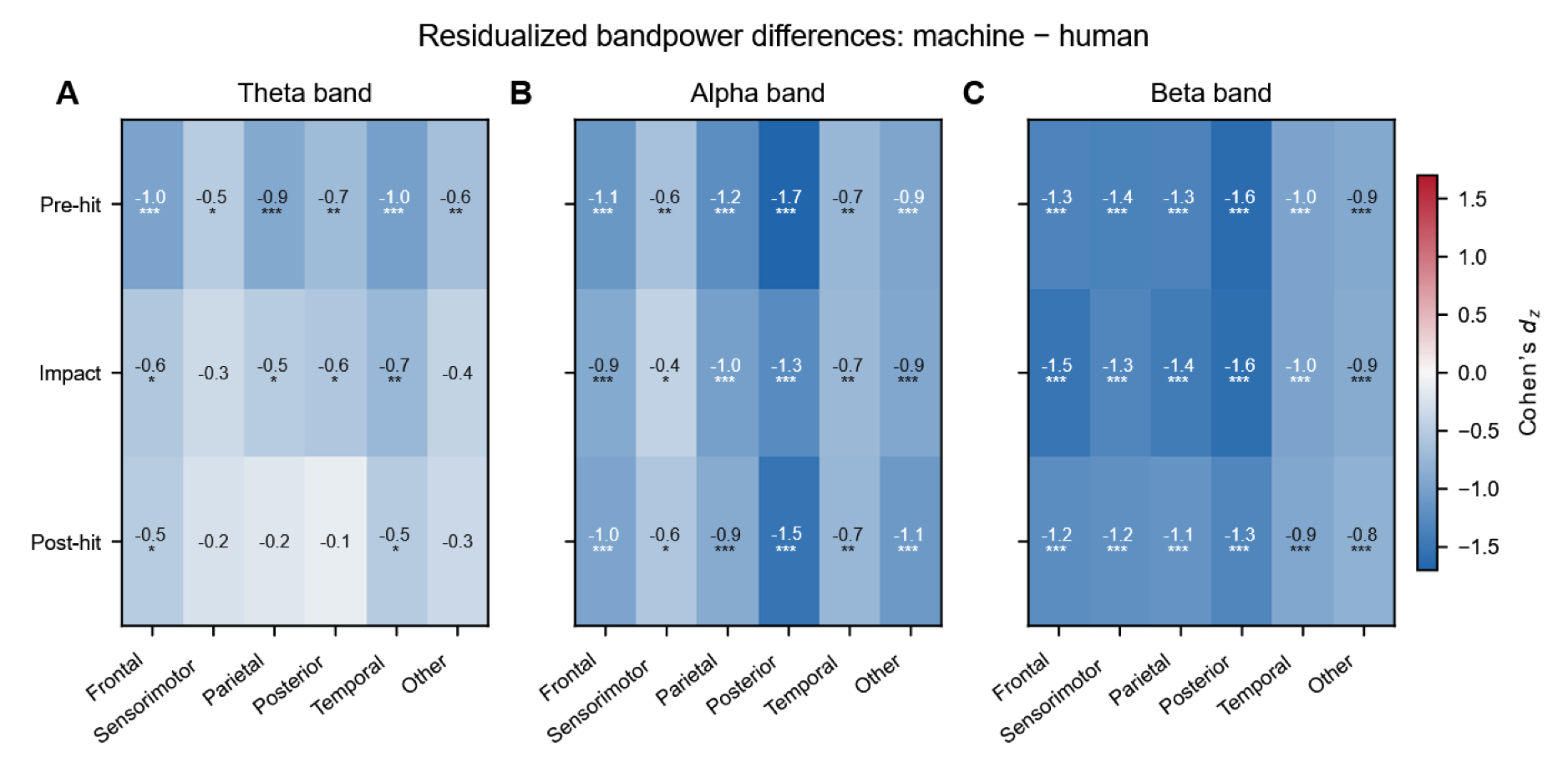
Residualized bandpower during human-opponent and ball-machine play. Cohen’s dz for machine-minus-human residualized bandpower in the theta (A), alpha (B), and beta (C) bands across three peri-hit windows and six sensor regions. Trial-level log bandpower was residualized against noise-layer RMS, neck-EMG RMS, and motion/miscellaneous-channel RMS before participant-level condition comparisons. A single symmetric color scale is used across all panels; negative values indicate lower residualized power during ball-machine play. Asterisks indicate Benjamini–Hochberg-adjusted paired t-test values across the complete 54-test residualized-bandpower family (*q < 0.05, **q < 0.01, ***q < 0.001). n = 25 participants. EMG = electromyography; RMS = root mean square.

The strongest residualized alpha effect was observed over posterior sensors during the pre-hit interval. The mean residualized value was 0.0340 during human-opponent play and -0.0308 during ball-machine play, producing a machine-minus-human-opponent difference of -0.0647. The paired comparison gave t(24) = -8.44, p < .001, paired-t FDR q < .001, dz = -1.687, Wilcoxon p < .001, and Wilcoxon FDR q < .001. Posterior alpha was also lower during ball-machine play in the impact interval, with a difference of -0.0583, t(24) = -6.29, both FDR q < .001, and dz = -1.258, and in the post-hit interval, with a difference of -0.0590, t(24) = -7.61, both FDR q < .001, and dz = -1.522.

The strongest pre-hit beta effect was also posterior. The human-opponent mean was 0.0412 and the ball-machine mean was -0.0349, giving a difference of -0.0761, t(24) = -8.09, p < .001, paired-t FDR q < .001, dz = -1.618, Wilcoxon p < .001, and Wilcoxon FDR q < .001. The posterior impact beta effect was similar, with a human-opponent mean of 0.0418, a ball-machine mean of -0.0344, and a difference of -0.0762, t(24) = -7.92, both FDR q < .001, and dz = -1.584. The post-hit posterior beta difference was -0.0687, t(24) = -6.53, both FDR q < .001, and dz = -1.307.

Large beta differences were not restricted to posterior sensors. During impact, the differences were -0.0536 over frontal sensors, t(24) = -7.52, dz = -1.504; -0.0546 over parietal sensors, t(24) = -7.14, dz = -1.428; and -0.0441 over sensorimotor sensors, t(24) = -6.56, dz = -1.311. Each of these comparisons produced paired-t and Wilcoxon FDR q < .001. The complete beta family therefore indicated a generalized reduction in residualized beta power during ball-machine play rather than an effect restricted to one sensor region.

Theta effects were more variable across time. All six pre-hit theta comparisons survived both corrections, with standardized effects ranging from dz = -0.505 over sensorimotor sensors to dz = -1.017 over temporal sensors. During impact, theta differences survived both procedures over frontal, parietal, posterior, and temporal sensors, whereas the sensorimotor comparison did not survive either correction, with a difference of -0.0114, t(24) = -1.66, paired-t FDR q = .116, Wilcoxon FDR q = .107, and dz = -0.333. The impact theta difference in the other sensor category also did not survive correction, with a difference of -0.0240, t(24) = -2.09, paired-t FDR q = .052, Wilcoxon FDR q = .070, and dz = -0.419.

Post-hit theta showed the least consistent evidence. The temporal comparison survived both procedures, with a difference of -0.027, t(24) = -2.54, paired-t FDR q = .022, Wilcoxon FDR q = .037, and dz = -0.508. The frontal comparison survived the paired-t family but not the Wilcoxon family, with a difference of -0.0249, t(24) = -2.49, paired-t FDR q = .023, Wilcoxon FDR q = .052, and dz = -0.497. Post-hit theta comparisons over sensorimotor, parietal, posterior, and other sensors did not survive either FDR procedure, with effect sizes ranging from dz = -0.111 to -0.348.

The only alpha comparison that did not survive both inferential procedures was the impact sensorimotor effect. Its mean difference was -0.0143, t(24) = -2.16, paired-t FDR q = .046, Wilcoxon FDR q = .083, and dz = -0.432. This divergence between the parametric and rank-based results suggests that the effect was less stable than the broader alpha and beta pattern and should not be emphasized as a central finding.

These analyses show that lower residualized alpha and beta power during ball-machine play remained after trial-level variance associated with noise-layer RMS, neck-EMG RMS, and motion/miscellaneous RMS had been removed. The findings are therefore described as nuisance-residualized bandpower effects rather than aperiodic-adjusted power. Table 3 reports the complete 54-comparison family, while Supplementary Table S2 provides the full descriptive and inferential statistics.

**Table 3.** Residualized bandpower results. All 54 participant-level comparisons of nuisance-residualized theta-, alpha-, and beta-band power across three peri-hit windows and six sensor regions.

| Window | Band | Sensor region | Participants | Human residualized mean | Machine residualized mean | Machine – human residualized mean | Cohen's dz | Paired t FDR q | Wilcoxon FDR q |
| --- | --- | --- | --- | --- | --- | --- | --- | --- | --- |
| Pre-hit | Theta | Frontal | 25 | 0.023 | -0.017 | -0.040 | -0.982 | <0.001 | <0.001 |
|  |  | Sensorimotor | 25 | 0.006 | -0.008 | -0.014 | -0.505 | 0.022 | 0.015 |
|  |  | Parietal | 25 | 0.012 | -0.010 | -0.021 | -0.910 | <0.001 | <0.001 |
|  |  | Posterior | 25 | 0.011 | -0.013 | -0.025 | -0.693 | 0.003 | 0.006 |
|  |  | Temporal | 25 | 0.019 | -0.017 | -0.037 | -1.017 | <0.001 | <0.001 |
|  |  | Other | 25 | 0.022 | -0.015 | -0.037 | -0.649 | 0.005 | 0.005 |
|  | Alpha | Frontal | 25 | 0.018 | -0.019 | -0.037 | -1.106 | <0.001 | <0.001 |
|  |  | Sensorimotor | 25 | 0.007 | -0.012 | -0.019 | -0.633 | 0.006 | 0.008 |
|  |  | Parietal | 25 | 0.014 | -0.019 | -0.034 | -1.212 | <0.001 | <0.001 |
|  |  | Posterior | 25 | 0.034 | -0.031 | -0.065 | -1.687 | <0.001 | <0.001 |
|  |  | Temporal | 25 | 0.014 | -0.017 | -0.031 | -0.745 | 0.002 | 0.002 |
|  |  | Other | 25 | 0.040 | -0.027 | -0.067 | -0.950 | <0.001 | <0.001 |
|  | Beta | Frontal | 25 | 0.027 | -0.024 | -0.052 | -1.339 | <0.001 | <0.001 |
|  |  | Sensorimotor | 25 | 0.022 | -0.024 | -0.046 | -1.362 | <0.001 | <0.001 |
|  |  | Parietal | 25 | 0.025 | -0.028 | -0.053 | -1.344 | <0.001 | <0.001 |
|  |  | Posterior | 25 | 0.041 | -0.035 | -0.076 | -1.618 | <0.001 | <0.001 |
|  |  | Temporal | 25 | 0.040 | -0.033 | -0.072 | -0.974 | <0.001 | <0.001 |
|  |  | Other | 25 | 0.042 | -0.030 | -0.071 | -0.908 | <0.001 | <0.001 |
| Impact | Theta | Frontal | 25 | 0.019 | -0.012 | -0.031 | -0.557 | 0.013 | 0.014 |
|  |  | Sensorimotor | 25 | 0.006 | -0.006 | -0.011 | -0.333 | 0.116 | 0.107 |
|  |  | Parietal | 25 | 0.010 | -0.008 | -0.018 | -0.500 | 0.023 | 0.019 |
|  |  | Posterior | 25 | 0.010 | -0.011 | -0.021 | -0.566 | 0.012 | 0.014 |
|  |  | Temporal | 25 | 0.018 | -0.016 | -0.034 | -0.743 | 0.002 | 0.001 |
|  |  | Other | 25 | 0.016 | -0.008 | -0.024 | -0.419 | 0.052 | 0.070 |
|  | Alpha | Frontal | 25 | 0.017 | -0.017 | -0.034 | -0.893 | <0.001 | <0.001 |
|  |  | Sensorimotor | 25 | 0.005 | -0.010 | -0.014 | -0.432 | 0.046 | 0.083 |
|  |  | Parietal | 25 | 0.014 | -0.018 | -0.032 | -1.010 | <0.001 | <0.001 |
|  |  | Posterior | 25 | 0.031 | -0.027 | -0.058 | -1.258 | <0.001 | <0.001 |
|  |  | Temporal | 25 | 0.013 | -0.016 | -0.029 | -0.724 | 0.002 | 0.003 |
|  |  | Other | 25 | 0.039 | -0.024 | -0.063 | -0.929 | <0.001 | <0.001 |
|  | Beta | Frontal | 25 | 0.029 | -0.025 | -0.054 | -1.504 | <0.001 | <0.001 |
|  |  | Sensorimotor | 25 | 0.022 | -0.022 | -0.044 | -1.311 | <0.001 | <0.001 |
|  |  | Parietal | 25 | 0.027 | -0.028 | -0.055 | -1.428 | <0.001 | <0.001 |
|  |  | Posterior | 25 | 0.042 | -0.034 | -0.076 | -1.584 | <0.001 | <0.001 |
|  |  | Temporal | 25 | 0.040 | -0.032 | -0.072 | -0.982 | <0.001 | <0.001 |
|  |  | Other | 25 | 0.041 | -0.029 | -0.070 | -0.881 | <0.001 | <0.001 |
| Post-hit | Theta | Frontal | 25 | 0.016 | -0.009 | -0.025 | -0.497 | 0.023 | 0.052 |
|  |  | Sensorimotor | 25 | 0.003 | -0.004 | -0.008 | -0.250 | 0.232 | 0.198 |
|  |  | Parietal | 25 | 0.005 | -0.002 | -0.007 | -0.192 | 0.354 | 0.466 |
|  |  | Posterior | 25 | 0.002 | -0.003 | -0.005 | -0.111 | 0.585 | 0.578 |
|  |  | Temporal | 25 | 0.015 | -0.012 | -0.027 | -0.508 | 0.022 | 0.037 |
|  |  | Other | 25 | 0.014 | -0.005 | -0.019 | -0.348 | 0.102 | 0.107 |
|  | Alpha | Frontal | 25 | 0.017 | -0.018 | -0.035 | -0.968 | <0.001 | <0.001 |
|  |  | Sensorimotor | 25 | 0.007 | -0.013 | -0.019 | -0.568 | 0.012 | 0.011 |
|  |  | Parietal | 25 | 0.014 | -0.018 | -0.031 | -0.879 | <0.001 | <0.001 |
|  |  | Posterior | 25 | 0.031 | -0.028 | -0.059 | -1.522 | <0.001 | <0.001 |
|  |  | Temporal | 25 | 0.013 | -0.018 | -0.031 | -0.746 | 0.002 | <0.001 |
|  |  | Other | 25 | 0.037 | -0.024 | -0.061 | -1.099 | <0.001 | <0.001 |
|  | Beta | Frontal | 25 | 0.026 | -0.022 | -0.048 | -1.233 | <0.001 | <0.001 |
|  |  | Sensorimotor | 25 | 0.021 | -0.022 | -0.044 | -1.198 | <0.001 | <0.001 |
|  |  | Parietal | 25 | 0.024 | -0.026 | -0.050 | -1.147 | <0.001 | <0.001 |

**Table 3.** Residualized bandpower results. All 54 participant-level comparisons of nuisance-residualized theta-, alpha-, and beta-band power across three peri-hit windows and six sensor regions. (continued)
| Window | Band | Sensor region | Participants | Human residualized mean | Machine residualized mean | Machine – human residualized mean | Cohen's dz | Paired t FDR q | Wilcoxon FDR q |
| --- | --- | --- | --- | --- | --- | --- | --- | --- | --- |
|  |  | Posterior | 25 | 0.037 | -0.031 | -0.069 | -1.307 | <0.001 | <0.001 |
|  |  | Temporal | 25 | 0.039 | -0.031 | -0.069 | -0.918 | <0.001 | <0.001 |
|  |  | Other | 25 | 0.038 | -0.026 | -0.064 | -0.815 | <0.001 | <0.001 |

*Note.* Trial-level log bandpower was residualized against noise-layer RMS, neck-EMG RMS, and motion/miscellaneous-channel RMS before within-participant condition averages were compared. Negative machine – human values indicate lower residualized power during ball-machine play. Benjamini–Hochberg correction was applied across the 54 tests separately for paired t tests and Wilcoxon signed-rank tests.

### 3.4 Spectral-model quality and peak detection

Spectral-model quality was evaluated across the initial short-window specification, resolution-aware max6, resolution-aware max3, two-second max6, and two-second aperiodic-only specifications. Because spectral parameterization treats periodic peaks as fitted components of a broader model rather than as independent observations, peak-detection rates were interpreted as descriptive diagnostics rather than as a separate inferential dataset (Donoghue et al., 2020).

Across the 900 participant x context x window x region cells in the principal resolution-aware max6 short-window model, the mean R^2^ was 0.869 and the median R^2^ was 0.950. Mean model error was 0.039 and median error was 0.033. A total of 175 of 900 cells, or 19.4%, had R^2^ < .80. Fit quality was not uniform across sensor regions. Mean temporal-region R^2^ values were 0.653 for pre-hit, 0.646 for impact, and 0.644 for post-hit spectra, whereas the corresponding mean values in the parietal region were 0.883, 0.872, and 0.894. Posterior means were 0.887, 0.898, and 0.902. The lower temporal fit quality indicates that temporal exponent results should be interpreted more cautiously than the posterior and parietal findings.

Reducing the maximum number of periodic peaks from six to three produced only a modest overall change in fit. The mean R^2^ declined from 0.869 in the max6 model to 0.863 in the max3 model, while mean error increased from 0.039 to 0.042. Across all 900 paired model cells, 675 cells, or 75.0%, had tied BIC values. The max3 model had the lower BIC in 140 cells, whereas max6 had the lower BIC in 85 cells. The mean max3-minus-max6 BIC difference was 0.352 and the median difference was exactly zero. This combination indicates that the two specifications frequently produced equivalent fitted structures, while the cells that differed did not show an overwhelming preference for one model. These quality and BIC summaries are illustrated in Supplementary Figure S3.

Periodic-peak detection varied with temporal window and model specification. In the resolution-aware max6 short-window model, theta peaks were detected in 21 of 300 pre-hit cells, 23 of 300 impact cells, and 23 of 300 post-hit cells, corresponding to rates of 7.0%, 7.7%, and 7.7%. Alpha peaks were detected in 114 pre-hit cells, 88 impact cells, and 116 post-hit cells, giving rates of 38.0%, 29.3%, and 38.7%. Beta peaks were detected in 84, 47, and 74 cells, corresponding to 28.0%, 15.7%, and 24.7%.

The better-resolved two-second max6 model produced higher peak-detection rates. Across its 300 cells, theta peaks were detected in 33 cells, or 11.0%, alpha peaks in 145 cells, or 48.3%, and beta peaks in 150 cells, or 50.0%. The increase was most apparent for beta peaks, for which the two-second detection rate of 50.0% exceeded the 15.7-28.0% range obtained from the short-window max6 analyses. This pattern is consistent with the finer 0.499-Hz frequency grid providing more information for periodic-component estimation than the 1.24-1.98-Hz short-window grids. It does not, however, establish that every peak detected in the longer window represented a stable physiological oscillation.

The aperiodic-only model contained no theta, alpha, or beta peaks in any of the 300 model cells because periodic fitting had been prohibited. This absence should not be interpreted as an empirical failure to find oscillations. It was a direct consequence of setting the maximum number of peaks to zero. The aperiodic-only specification also provided a poorer overall description of the spectra than the two-second max6 model. Mean R^2^ decreased by 0.117 and mean error increased by 0.028. Only 1.3% of cells had a higher R^2^ under the aperiodic-only specification, and only 1.0% had a lower model error. The aperiodic-only model was therefore not the better-fitting general spectral model. Its value was restricted to testing whether the focal context difference remained when periodic components could not be fitted.

Peak-detection sensitivity is shown in Supplementary Figure S4. The full PSD settings, frequency resolutions, and model parameters are reported in Supplementary Table S4, while complete quality and peak-detection summaries are provided in Supplementary Table S5.

### 3.5 Robustness of the focal parietal aperiodic-exponent effect

The impact-window parietal effect remained similar across all three short-window specifications. Under the initial short-window specification, the human-opponent mean was 0.653 and the ball-machine mean was 0.784, giving a mean difference of 0.132. The paired comparison produced t(24) = 3.67, p = .001, paired-t FDR q = .003, dz = 0.733, Wilcoxon p < .001, and Wilcoxon FDR q = .002. Twenty of 25 participants showed a higher exponent during ball-machine play. The 50,000-resample paired-bootstrap 95% confidence interval for the mean difference extended from 0.065 to 0.203.

Under the principal resolution-aware max6 specification, the human-opponent mean was 0.642 and the ball-machine mean was 0.782. The mean difference was 0.141, t(24) = 3.84, p < .001, paired-t FDR q = .002, dz = 0.768, Wilcoxon p < .001, and Wilcoxon FDR q = .002. Twenty participants showed the machine-higher direction, and the paired-bootstrap 95% confidence interval was 0.072-0.213.

The resolution-aware max3 specification produced a human-opponent mean of 0.650 and a ball-machine mean of 0.777, resulting in a difference of 0.128, t(24) = 3.68, p = .001, paired-t FDR q = .003, dz = 0.737, Wilcoxon p < .001, and Wilcoxon FDR q = .002. The positive direction was again observed in 20 participants, and the bootstrap interval was 0.063-0.197. Thus, reducing the maximum permitted number of periodic peaks from six to three changed the estimated difference by only 0.013.

Participant-level effects were also highly concordant across the short-window models. The effects from the initial short-window specification and the resolution-aware max6 model had Pearson r = .972 and Spearman *ρ* = .977. The resolution-aware max6 and max3 effects had Pearson r = .969 and Spearman *ρ* = .977, with a mean absolute participant-level effect difference of 0.014. No participant changed the direction of the parietal effect across the three short-window specifications. The short-window robustness results are illustrated in Figure 3A-B.

**Figure 3.**
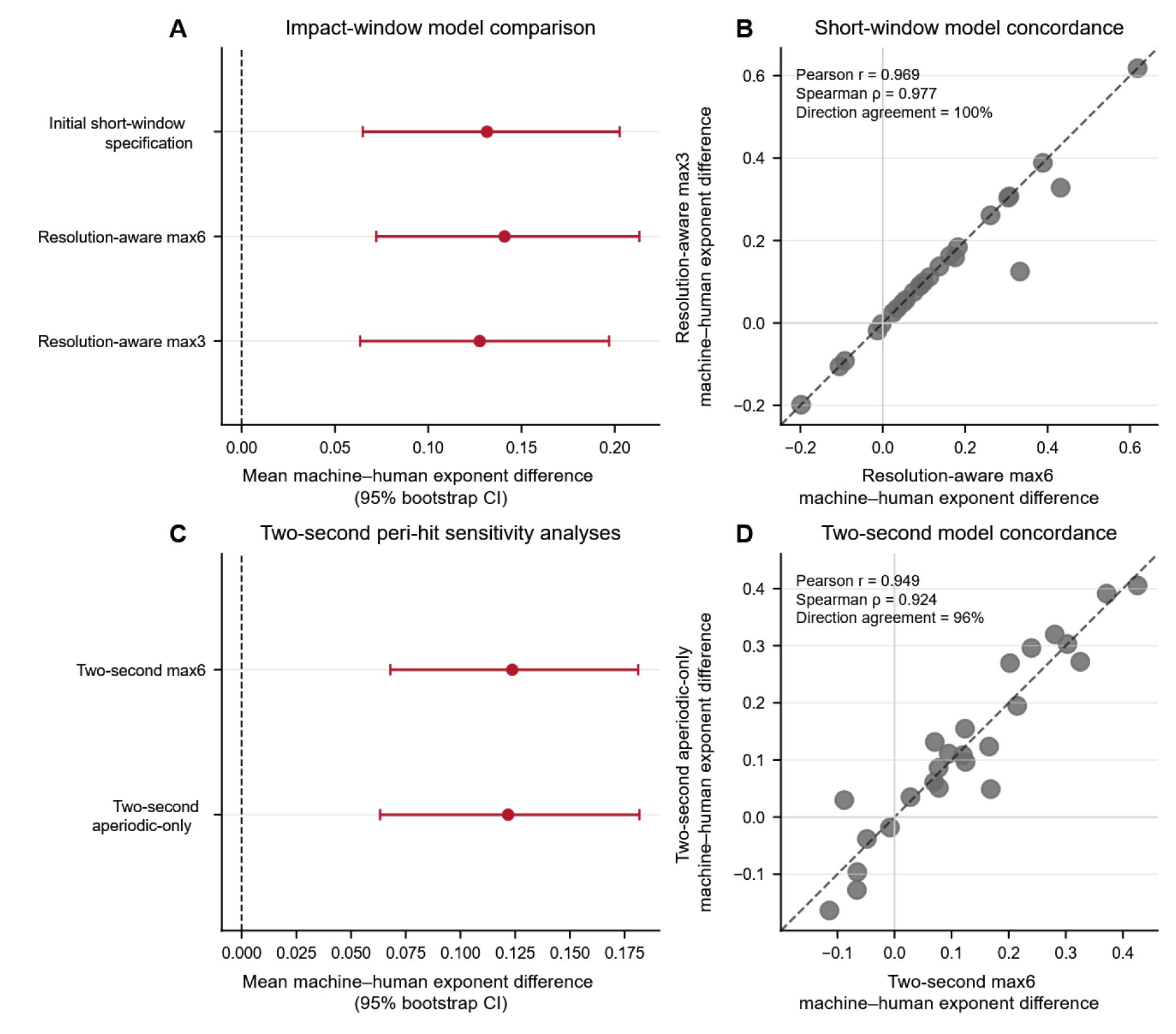
Robustness of the focal parietal aperiodic-exponent effect across spectral models. A, mean parietal machine-minus-human exponent differences for the initial short-window specification, resolution-aware max6, and resolution-aware max3 impact-window models, with 95% paired-bootstrap confidence intervals. B, participant-level concordance between the resolution-aware max6 and max3 impact-window effects; the dashed line is the identity line, and Pearson r, Spearman *ρ*, and direction agreement are reported in the panel. C, mean parietal machine-minus-human exponent differences for the two-second max6 and two-second aperiodic-only sensitivity analyses, with 95% paired-bootstrap confidence intervals. D, participant-level concordance between the two two-second estimates; the dashed line is the identity line, and Pearson r, Spearman *ρ*, and direction agreement are reported in the panel. Positive values indicate a higher aperiodic exponent during ball-machine than human-opponent play. Bootstrap intervals were based on 50,000 paired resamples. The impact-window and two-second peri-hit estimates are distinct temporal estimands. The aperiodic-only model was used as a sensitivity analysis rather than as the preferred overall spectral model. N = 25 participants.

The better-resolved two-second max6 analysis also retained a positive parietal context difference. The mean exponent was 0.659 during human-opponent play and 0.783 during ball-machine play, resulting in a difference of 0.124, t(24) = 4.20, p < .001, paired-t FDR q < .001, dz = 0.839, Wilcoxon p < .001, and Wilcoxon FDR q < .001. Nineteen of 25 participants showed a higher ball-machine exponent. The paired-bootstrap 95% confidence interval was 0.068-0.181.

The two-second aperiodic-only analysis produced a similar context effect despite its poorer overall spectral fit. The human-opponent and ball-machine means were 0.629 and 0.751, giving a difference of 0.122, t(24) = 3.93, p < .001, paired-t FDR q = .001, dz = 0.787, Wilcoxon p < .001, and Wilcoxon FDR q = .001. Twenty participants showed a higher exponent during ball-machine play, and the bootstrap interval was 0.063-0.182. Bootstrap uncertainty was estimated using paired resampling of the participant effects.

The two-second max6 and aperiodic-only participant effects were strongly related, with Pearson r = .949 and Spearman *ρ* = .924. Their group mean differences differed by only 0.002. The mean absolute participant-level difference between models was 0.037, and 24 of 25 participants retained the same effect direction, corresponding to 96.0% direction agreement. This concordance is shown in Figure 3C-D and Supplementary Figure S5. The similarity of the context effects does not mean that the aperiodic-only model was a superior spectral model. Rather, it indicates that the machine-human parietal difference was not removed when periodic peaks were excluded from the model.

Leave-one-participant-out analyses provided additional evidence that the focal finding was not dependent on a single observation. For the initial short-window specification, all 25 omitted-participant analyses retained positive differences, with means ranging from 0.111 to 0.145, Cohen’s dz values from 0.691 to 0.858, paired-t FDR q values from .001 to .007, and Wilcoxon FDR q values from .001 to .004. Under the principal max6 model, the omitted-participant mean differences ranged from 0.121 to 0.155, dz ranged from 0.726 to 0.896, paired-t FDR q ranged from .001 to .004, and Wilcoxon FDR q ranged from .001 to .004. Under max3, the mean differences ranged from 0.107 to 0.141 and all paired-t and Wilcoxon FDR q values remained below .01.

The two-second analyses showed the same pattern. After each participant was omitted from the max6 analysis, the mean difference remained positive and ranged from 0.111 to 0.134, with dz = 0.798-0.942 and both FDR q values at or below .001. For the aperiodic-only model, omitted-participant differences ranged from 0.110 to 0.134, dz = 0.745-0.916, paired-t FDR q = .001-.003, and Wilcoxon FDR q = .001-.003. All five model specifications therefore retained a positive and FDR-corrected effect in every leave-one-participant-out analysis.

The two-second fit-quality audit showed that participant-level exponent differences were associated with differences in model fit. The exponent effect correlated positively with the machine-minus-human difference in R^2^, Spearman *ρ* = .637, permutation p < .001, and negatively with the difference in model error, *ρ* = -.516, permutation p = .009. These associations indicate that fit-quality differences and exponent differences were not independent and support caution in interpreting the individual variation.

However, the estimated context effect remained positive when the modeled difference in fit quality was zero. For the R^2^-based analysis, the ordinary least-squares intercept at Δ R^2^ = 0 was 0.105, with an HC3 95% confidence interval of 0.055-0.155 and a paired-bootstrap interval of 0.059-0.158. For the model-error analysis, the corresponding intercept was 0.095, with an HC3 interval of 0.047-0.143 and a bootstrap interval of 0.053-0.146. One participant exceeded the 4/n Cook’s-distance screening threshold in each regression. These analyses were exploratory because exponent and fit quality were produced by the same model, but the positive quality-neutral intercepts suggest that the group difference was not explained entirely by the ball-machine spectra receiving a better fit. The complete audit is illustrated in Supplementary Figure S6.

Table 4 summarizes the five focal model specifications, their effect sizes, FDR-adjusted results, bootstrap intervals, and leave-one-participant-out outcomes. Supplementary Table S6 provides the complete short-window concordance, bootstrap, leave-one-participant-out, BIC, two-second, fit-quality, and aperiodic-only results. The short impact-window and two-second peri-hit estimates should not be treated as exact replications of a single temporal quantity. The impact models estimate activity within -0.2 to +0.3 s of participant contact, whereas the two-second models average across -1.0 to +1.0 s. Their agreement therefore supports robustness of the broader peri-hit context effect while retaining the distinction between temporally specific and broad-window estimates.

**Table 4.**
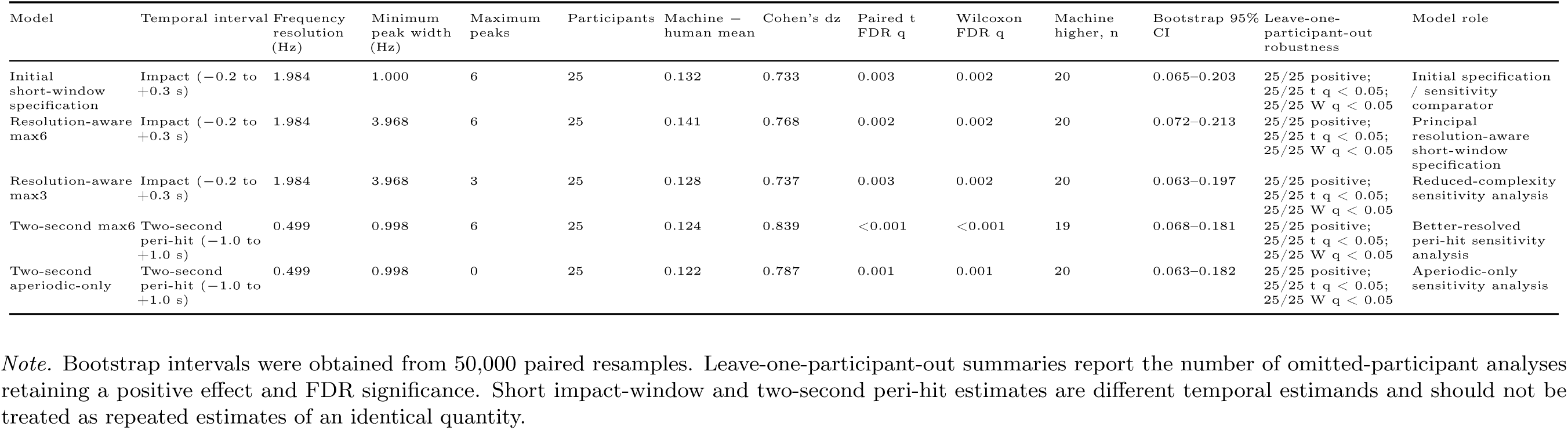
Robustness of the focal parietal effect. Robustness of the focal parietal machine-minus-human aperiodic-exponent difference across impact-window and two-second peri-hit spectral specifications.

| Model | Temporal interval | Frequency resolution (Hz) | Minimum peak width (Hz) | Maximum peaks | Participants | Machine – human mean | Cohen's dz | Paired t FDR q | Wilcoxon FDR q | Machine higher, n | Bootstrap 95% CI | Leave-one-participant-out robustness | Model role |
| --- | --- | --- | --- | --- | --- | --- | --- | --- | --- | --- | --- | --- | --- |
| Initial short-window specification | Impact (–0.2 to +0.3 s) | 1.984 | 1.000 | 6 | 25 | 0.132 | 0.733 | 0.003 | 0.002 | 20 | 0.065–0.203 | 25/25 positive; 25/25 t q < 0.05; 25/25 W q < 0.05 | Initial specification / sensitivity comparator |
| Resolution-aware max6 | Impact (–0.2 to +0.3 s) | 1.984 | 3.968 | 6 | 25 | 0.141 | 0.768 | 0.002 | 0.002 | 20 | 0.072–0.213 | 25/25 positive; 25/25 t q < 0.05; 25/25 W q < 0.05 | Principal resolution-aware short-window specification |
| Resolution-aware max3 | Impact (–0.2 to +0.3 s) | 1.984 | 3.968 | 3 | 25 | 0.128 | 0.737 | 0.003 | 0.002 | 20 | 0.063–0.197 | 25/25 positive; 25/25 t q < 0.05; 25/25 W q < 0.05 | Reduced-complexity sensitivity analysis |
| Two-second max6 | Two-second peri-hit (–1.0 to +1.0 s) | 0.499 | 0.998 | 6 | 25 | 0.124 | 0.839 | <0.001 | <0.001 | 19 | 0.068–0.181 | 25/25 positive; 25/25 t q < 0.05; 25/25 W q < 0.05 | Better-resolved peri-hit sensitivity analysis |
| Two-second aperiodic-only | Two-second peri-hit (–1.0 to +1.0 s) | 0.499 | 0.998 | 0 | 25 | 0.122 | 0.787 | 0.001 | 0.001 | 20 | 0.063–0.182 | 25/25 positive; 25/25 t q < 0.05; 25/25 W q < 0.05 | Aperiodic-only sensitivity analysis |

*Note.* Bootstrap intervals were obtained from 50,000 paired resamples. Leave-one-participant-out summaries report the number of omitted-participant analyses retaining a positive effect and FDR significance. Short impact-window and two-second peri-hit estimates are different temporal estimands and should not be treated as repeated estimates of an identical quantity.

## 4 Discussion

### 4.1 Summary of main findings

The present analysis examined whether human-opponent and ball-machine table-tennis contexts were associated with different event-locked EEG spectral patterns surrounding participant racket–ball contact. The results showed that context-related differences remained after directly recorded nuisance signals had been considered, but the clearest difference was expressed in the aperiodic structure of the power spectrum rather than being adequately characterized as a change in one canonical frequency band. Nuisance-residualized bandpower was generally lower during ball-machine than human-opponent play, particularly for posterior alpha and beta activity. This pattern indicates that the context difference was not removed when trial-level variance associated with noise-layer, neck-EMG, and motion or miscellaneous signals was residualized.

Spectral parameterization provided a different but complementary description of this pattern. In the principal resolution-aware max6 analysis, the aperiodic exponent was generally higher during ball-machine play, with the strongest and most consistent anatomically interpretable effects occurring over posterior and parietal sensors. Posterior differences were present before contact and were particularly clear after contact, while the parietal effect was evident around impact and after the hit. Participant-level direction consistency was also greatest for these posterior and parietal effects. The corresponding estimates and inferential statistics are reported in the Results and Table 2.

The focal parietal impact effect was retained under every spectral specification examined: the initial short-window model, the principal resolution-aware max6 model, the reduced-complexity max3 model, the better-resolved two-second model, and the aperiodic-only stress test. The bootstrap intervals for all specifications remained on the positive side of zero. Thus, the focal difference was not dependent on one choice of peak-width limit, peak count, window duration, or periodic-peak treatment.

The focal result was also not attributable to a single influential participant. All 25 leave-one-participant-out analyses retained a positive difference and FDR significance under each of the five model specifications. This does not constitute an independent replication, because all sensitivity analyses used the same participant sample. It does, however, show that the observed effect was not dependent on one peak-width setting, one level of model complexity, one temporal window, one treatment of periodic peaks, or one participant. The overall conclusion is therefore that human-opponent and ball-machine play were associated with different peri-hit spectral structures, with the most robust difference expressed as a steeper aperiodic exponent during ball-machine play over posterior and parietal sensors.

Taken together, these results indicate that the analysis was effective in a specific methodological sense: it detected participant-consistent differences between two naturalistic visuomotor contexts under conditions involving substantial whole-body movement and also provided a more specific description of those differences than conventional bandpower alone. This should not be taken to mean that the analysis identified a unique neural mechanism or that it has been validated for classifying contexts in new participants. Rather, its value here is that the combination of nuisance-informed analysis, spectral parameterization, resolution-aware modelling, and sensitivity testing identified a context-associated effect that remained stable across several reasonable analytical specifications and showed that the dominant difference was expressed in broadband aperiodic structure.

### 4.2 Interpretation of the aperiodic exponent

The aperiodic exponent describes how rapidly broadband spectral power decreases as frequency increases. A larger exponent corresponds to a steeper fitted spectral slope, whereas a smaller exponent corresponds to a flatter spectrum. This description should be distinguished from a direct psychological interpretation. Spectral parameterization is valuable because conventional bandpower measures can combine variation in periodic peak properties with variation in the aperiodic exponent and offset. A change measured within alpha or beta frequencies may therefore arise from a narrowband oscillation, a shift in peak frequency or bandwidth, a broader change in aperiodic structure, or some combination of these processes (Donoghue, Dominguez, et al., 2020; Donoghue, Haller, et al., 2020).

The higher ball-machine exponent should not be translated directly into greater attention, effort, engagement, workload, or arousal. Each of these psychological states can be associated with multiple neural and peripheral processes, and none was measured directly in the present analysis. Aperiodic activity itself may have several physiological origins. Recent evidence suggests that it is influenced principally by population-level postsynaptic currents but may also reflect several interacting circuit, cellular, and measurement processes (Preston et al., 2026). The exponent is consequently a meaningful description of broadband electrophysiological structure, but it is not a one-to-one index of a particular mental operation.

Computational modelling and invasive electrophysiological evidence have linked changes in power-law slope to variation in the balance of excitatory and inhibitory synaptic activity. Gao et al. (2017), for example, demonstrated that simulated and experimentally induced changes in excitation–inhibition balance could be reflected in the exponent of local field potential and electrocorticographic spectra. However, this relationship was established under conditions that differ substantially from sensor-level scalp EEG recorded during whole-body movement. Scalp measurements contain mixtures of neural generators, are affected by referencing and volume conduction, and may retain muscle and movement contamination. The present findings therefore do not provide a direct measurement of cortical excitation–inhibition balance. At most, they identify a context-associated change in a signal property that has been related to excitation–inhibition dynamics in other recording modalities and experimental settings.

The posterior and parietal distribution is consistent with a difference in the neural processes supporting visual analysis, spatial prediction, and the transformation of visual information into an interceptive response. The presence of posterior effects before contact is compatible with a contribution from anticipatory processing, while persistence during and after contact could reflect continuing visual, somatosensory, motor, or outcome-related processing. This interpretation remains intentionally broad. Sensor-region effects do not identify a specific cortical generator, and the available data cannot establish whether the underlying difference was driven by predictability, biological motion, interpersonal interaction, movement execution, visual trajectory statistics, or another contextual property.

The additional sensitivity analyses strengthen confidence that the focal effect was not an artefact of the initial periodic-peak configuration. Similar participant-level differences were obtained after the minimum peak width was made compatible with spectral resolution, after the maximum number of peaks was reduced, and after periodic peaks were removed completely. The result can therefore be described as a robust aperiodic-exponent difference without implying that its physiological or psychological origin has been determined.

### 4.3 Temporal specificity, spectral resolution, and model robustness

The short analysis windows were selected to preserve distinctions between activity before contact, around impact, and after the hit. This temporal separation is scientifically important because perception, action preparation, contact-related movement, sensory consequences, and preparation for the next exchange are unlikely to contribute equally throughout the peri-hit interval. The posterior exponent difference was present in all three windows, while the parietal effect was strongest during impact and post-hit activity. Combining the entire interval into a single estimate would have concealed these temporal variations.

The advantage of temporal specificity was accompanied by reduced frequency resolution. The pre-hit, impact, and post-hit spectra had frequency-bin spacings of approximately 1.24, 1.98, and 1.42 Hz, respectively. The 0.5-s impact interval therefore had the coarsest spectral grid. This limitation is more serious for estimating narrow periodic peaks than for describing a broadband slope across several frequencies. When only a small number of spectral points are available across a putative oscillation, its centre frequency and bandwidth cannot be estimated with the same precision as they can from a longer segment.

Time-resolved parameterization methods demonstrate that both periodic and aperiodic properties can be studied dynamically, but they also make clear that temporal localization and spectral precision must be balanced when selecting an analysis window (Wilson et al., 2022). The short windows in the present analysis should consequently not be regarded as intrinsically invalid. Rather, they provide event-stage information at the cost of limited narrow-peak resolution.

The resolution-aware model addressed this limitation by preventing periodic components from being fitted at widths that could not be represented meaningfully by the frequency grid. This required the largest adjustment for the impact interval, which had the coarsest short-window resolution. The focal parietal effect remained very similar after this change, and the bootstrap intervals from the initial and resolution-aware specifications overlapped substantially. The context difference therefore remained stable when the initial specification was replaced by the resolution-aware specification.

The two-second analysis offered a complementary test. Its finer frequency grid allowed periodic peaks to be represented with greater spectral detail than in the short-window models. The parietal machine-minus-human difference nevertheless remained positive and of comparable magnitude. Agreement between the short and broad windows indicates that the general peri-hit difference was not restricted to spectra with coarse frequency resolution.

The two sets of estimates are nevertheless not interchangeable. The short impact model quantified activity from −0.2 to +0.3 s around contact, whereas the two-second model averaged from −1.0 to +1.0 s. A context difference in the two-second model could be driven by pre-hit, impact, post-hit, or sustained contributions across the interval. It cannot determine when within those two seconds the difference occurred. The broad-window result should therefore be treated as a better-resolved sensitivity analysis of general peri-hit activity rather than as an exact replication of the impact-window estimand.

This distinction has an important consequence for interpretation. Agreement across windows strengthens the conclusion that the context effect was not produced solely by the spectral limitations of the impact interval. It does not establish that the same neural process generated both estimates. Different event stages may contribute to the short and broad models in different proportions, even when both produce a positive participant-level context difference.

### 4.4 Model diagnostics support a predominantly aperiodic interpretation

Peak detection varied substantially with the temporal interval and spectral-model configuration. Alpha and beta peaks were detected less consistently in the short-window models than in the better-resolved two-second model, particularly during the impact interval. This shows that peak detection depended partly on the spectral information available to the model. A low short-window detection rate cannot by itself establish the absence of rhythmic activity, just as a higher two-second rate does not prove that every detected peak represented a stable physiological oscillation.

Reducing the maximum number of peaks from six to three had little effect on the focal context difference. Participant-level effects were strongly concordant across the two resolution-aware models, and no participant changed effect direction. This similarity suggests that the focal exponent difference was not created by the additional peak capacity of the max6 model.

The model-selection results also did not indicate a clear overall preference for either short-window peak-count specification. Most fitted spectra yielded the same Bayesian information criterion under max6 and max3, which is compatible with many spectra containing fewer periodic components than either model was permitted to fit. This supports the use of the max3 analysis as a parsimony check rather than as a fundamentally different account of the data.

The strongest stress test was provided by the two-second aperiodic-only model. Its participant-level context effects closely matched those from the two-second max6 model and were almost always in the same direction. This similarity indicates that the focal machine–human exponent difference did not depend on the model assigning part of each spectrum to periodic peaks.

This result should not be interpreted as showing that the aperiodic-only model was the better description of the spectra. Its overall fit was poorer than the max6 model, as expected when periodic structure was present but prohibited. The aperiodic-only model was useful because it imposed an extreme constraint and showed that the group difference remained under that constraint, not because oscillatory components were unnecessary for modelling the data.

The combined diagnostic evidence supports the description of the focal context difference as predominantly aperiodic, but it does not rule out periodic activity. The large residualized alpha and beta effects show that canonical-band power differed between contexts, but because these measures retain both periodic and aperiodic contributions, they do not establish a rhythmic mechanism. Periodic and aperiodic processes may also change together rather than acting as mutually exclusive explanations. Spectral parameterization is intended to reduce conflation between these components, not to imply that only one component can carry meaningful task information.

### 4.5 Human-opponent and ball-machine contexts

Human-opponent and ball-machine play shared the basic requirements of tracking an incoming ball, preparing a stroke, making racket–ball contact, and continuing the exchange. However, the information available before the participant’s response differ. Human-opponent play could provide advance information from body position, arm motion, racket orientation, and the temporal structure of the opponent’s action. Ball-machine play lacked these interpersonal kinematic signals and instead provided mechanically generated trajectories whose regularity depended on the programmed condition.

Research on perceptual expertise in sport shows that skilled performers can use advance movement information to anticipate action outcomes. Expert racquet-sport performers, for example, are better able to use pre-contact kinematics from an opponent’s body and equipment to predict the direction of a forthcoming stroke (Abernethy & Zawi, 2007). Meta-analytic evidence also indicates that sport expertise is associated with more effective perceptual-cognitive cue use and improved response accuracy and speed, although the size of the advantage varies with the task, sport, and presentation method (Mann et al., 2007).

Human action observation may additionally engage processes involved in representing and predicting goal-directed movement. This offers one possible reason that an opponent-generated ball could produce a different neural context from a mechanically generated ball, although broad mirror-system accounts should not be treated as a direct explanation for the present sensor-level effects (Rizzolatti & Craighero, 2004). No measure in the dataset directly quantified the participant’s use of biological-motion information.

Ball-machine play may instead rely more heavily on extracting timing and trajectory regularities after release. Mechanical delivery can be regular in some respects while differing from human play in others, including shot sequence, spin, movement preparation, and interpersonal contingency. It would therefore be inappropriate to describe either context as generally more predictable or more difficult without trial-level behavioral evidence. The higher machine-condition exponent could be associated with differences in cue availability, visual sampling, movement timing, stroke selection, strategic uncertainty, or a combination of these factors.

This distinction is relevant to representative learning design. Training tasks are most likely to preserve the informational demands of competitive performance when perception and action remain coupled to the cues normally available in the target environment (Pinder et al., 2011). Ball-machine practice may be useful for repetition, timing, and trajectory-based interception, whereas practice with a human opponent preserves biological-motion and interpersonal information. The present results suggest that these practice settings are not neurally interchangeable, but they do not establish that one produces superior learning.

The robustness analyses strengthen confidence in the existence of a context-associated spectral difference. They do not isolate which contextual property caused it. Experimental manipulation of delivery regularity, visibility of opponent kinematics, spin, ball speed, stroke type, and interpersonal contingency would be needed to distinguish among these explanations.

### 4.6 Implications for training, process engagement, and transfer

From a training perspective, the important implication is not that human-opponent practice is preferable to ball-machine practice, but that the two contexts may engage partly different processes and may therefore be useful for different purposes. Behavioral performance can show whether a person performs well or improves with practice, but it does not necessarily establish which sources of information or processing strategies supported that performance. EEG provides a potentially complementary level of evidence. In the present study, two contexts that shared the broad behavioral requirement of intercepting and returning a table-tennis ball were nevertheless associated with different peri-hit spectral states. This suggests that evaluation of a training method could, in future work, consider not only whether behavioral performance changes, but also whether the training context appears to engage the perceptual and neural processes considered relevant to the target performance. This is consistent with representative learning design, in which the usefulness of practice depends on the functional information and perception–action relationships preserved by the training environment (Pinder et al., 2011). It also fits with evidence that the cognitive effects associated with sport or fitness may be process-specific rather than automatically generalizing across tasks (Guchait et al., 2026).

A related possibility concerns individual differences. The same training context need not provide the same functional challenge to every performer. Motor-learning theory has long emphasized that the usefulness of a practice condition depends partly on the interaction between task difficulty and the performer’s skill level (Guadagnoli & Lee, 2004), and evidence from simulation training similarly indicates that the relationship between training fidelity and transfer can vary according to prior skill (Doozandeh & Hedayati, 2022). The present data cannot show that the observed EEG differences predict who would benefit from human-opponent or ball-machine practice. However, they provide a framework for testing this question prospectively. A future training study could combine behavioral improvement, retention or transfer measures with participant-specific EEG responses to different practice contexts. If neural measures were shown to predict subsequent improvement in an independent sample, they could potentially help identify whether a particular individual would benefit more from training emphasizing trajectory regularity, interpersonal prediction, temporal preparation, or another process relevant to performance.

The same general principle is not restricted to sport. Many forms of training use a simplified, simulated, virtual, or otherwise altered version of the situation in which the trained skill will eventually be performed. Transfer is consequently a central issue in areas such as occupational simulation, aviation, rehabilitation, and digital learning (Barnett & Ceci, 2002; Doozandeh & Hedayati, 2022). Neuroergonomic research has similarly proposed combining brain measures with behavioral performance to understand cognitive and physical work in realistic environments, including training and neuroadaptive systems (Mehta & Parasuraman, 2013), while EEG-based cognitive-state monitoring has been investigated as a basis for adapting instructional environments to the learner (Gerjets et al., 2014). The present findings do not demonstrate such applications directly. They nevertheless illustrate the underlying methodological possibility: when a training environment differs from the activity for which training is intended, EEG may provide an additional means of asking whether the two contexts engage similar or different processing, rather than relying on behavioral outcome alone.

### 4.7 Artifact-informed interpretation

Movement-related contamination is a central difficulty in mobile EEG. Cable movement, electrode displacement, neck-muscle activity, and head or body motion can produce spectral signals that overlap with neural frequencies. In the dual-layer table-tennis system, mechanically matched noise electrodes were designed to capture non-biological signal variation associated with movement. Earlier analysis of these recordings showed that scalp channels were more strongly related to their matched noise channels than to general head or body acceleration and that incorporating noise-layer information improved the recovery of brain-related components (Studnicki et al., 2022).

The residualized-bandpower and spectral-parameterization analyses addressed the artifact problem differently. Bandpower was directly residualized against trial-level noise-layer RMS, neck-EMG RMS, and motion/miscellaneous RMS. The resulting alpha and beta differences therefore represent effects remaining after the linear variance associated with those nuisance summaries was removed. This does not mean that the residuals contained only neural activity. Unmeasured signals, nonlinear relationships, local channel contamination, and nuisance effects not well represented by global RMS measures could remain.

The aperiodic exponent was not directly residualized against these nuisance variables. Instead, its reliability was examined through regional model quality, participant-level consistency, peak-setting sensitivity, bootstrap confidence intervals, leave-one-participant-out analyses, and agreement across spectral models. Describing the entire analysis as artifact-corrected or artifact-free would imply a stronger operation than was performed. The more accurate terms are artifact-informed or artifact-audited spectral analysis.

Model quality also varied regionally. The temporal sensor group had lower mean R^2^ than the posterior and parietal groups across the short windows. Temporal effects should therefore be interpreted more cautiously, even where they survived correction. The strongest conclusions concern posterior and parietal effects, where the fits were generally more stable and where participant-level direction consistency was greater.

The two-second fit-quality audit showed that participant-level exponent differences covaried with differences in model fit, confirming that exponent and fit-quality variation were not independent. However, the modeled context effect remained positive when the fit-quality difference was set to zero. This makes a purely fit-quality explanation less likely, although it cannot exclude all influence from recording quality or movement.

### 4.8 Limitations

The study was a secondary analysis of an existing public dataset. The experimental conditions, sensors, event labels, and behavioral measures were therefore limited to those collected for the original project. Although the dataset was unusually rich for a naturalistic sport recording, the analysis could not introduce new manipulations needed to separate predictability, biological motion, social interaction, strategic variation, ball trajectory, and movement execution.

Human-opponent and ball-machine trials differed in several properties simultaneously. The observed spectral differences cannot be attributed uniquely to the presence of another person or to mechanical delivery. Human play included cooperative and competitive trials, while machine trials included stationary and moving deliveries. Collapsing these subconditions increased the number of events within each broad context but also combined potentially meaningful variation. Future analyses could model these subconditions separately, provided that sufficient participant-level data are available.

Detailed behavioral and kinematic modelling was not included. No trial-level measure of anticipation accuracy, stroke quality, return success, ball speed, spin, gaze position, joint movement, or perceived effort was related directly to the EEG outcomes. Consequently, explanations in terms of prediction, visual information, motor preparation, engagement, or difficulty remain hypotheses rather than demonstrated mechanisms.

The implications for training should therefore be regarded as prospective rather than demonstrated outcomes of the present study. Participants were not assigned to alternative training programs, and the analysis did not measure learning rate, retention, transfer to another task, or participant-specific benefit from one practice context. The present results consequently show that the two contexts were associated with different EEG spectral states, but not that these differences caused different learning outcomes or that they can presently be used to prescribe training for an individual. Establishing that practical value would require longitudinal studies that combine neural measurements with behavioral and kinematic outcomes and include an independent assessment of transfer. Of particular interest would be whether an individual’s neural response to different practice contexts predicts subsequent improvement or retention beyond information already provided by behavioral performance.

The analysis was conducted at the sensor-region level. Labels such as posterior, parietal, and sensorimotor describe scalp distributions and do not identify the cortical generators responsible for the observed effects. Volume conduction, reference choice, spatial mixing, and non-neural sources all contribute to sensor-level EEG. The public dataset includes anatomical MRI and digitized electrode locations that could support future source-resolved analyses, although source localization during mobile behavior would introduce additional modelling assumptions.

Residual artifacts cannot be excluded. Direct nuisance residualization reduced linear variance associated with noise-layer, EMG, and movement signals, but no nuisance model can account for contamination that was not measured or was represented inadequately. The aperiodic outcomes were audited through fit quality and model sensitivity rather than directly residualized. They should therefore not be interpreted as purely neural measurements.

Short-window frequency resolution limited periodic-peak interpretation. The 1.24–1.98-Hz frequency grids could represent broadband slope but provided restricted information about narrow peaks. Resolution-aware peak-width limits reduced the risk of fitting components narrower than the grid could support, but they could not create spectral information that was absent from the short segments. Periodic peak frequency and bandwidth estimates from these windows should consequently be treated as secondary diagnostics.

The short impact-window and two-second peri-hit effects were different temporal estimands. Agreement between them supports a general peri-hit context effect, but the numerical estimates should not be treated as repeated measurements of an identical quantity. The broader interval cannot establish whether its effect arose before, during, or after contact.

The complete model hierarchy was not prospectively preregistered. The initial short-window specification was defined first, and the resolution-aware and broader-window analyses were added after the formal spectral-resolution audit identified the need to examine dependence on peak-width limits, model complexity, window duration, and periodic-peak modelling. These additional analyses should therefore be interpreted as sensitivity analyses within the same dataset rather than as independent confirmatory tests. Their consistency provides evidence of analytic robustness, but it does not provide the same separation between prediction and postdiction as prospective preregistration (Nosek et al., 2018).

Adjacent short windows shared one 4-ms boundary sample because MNE cropping included both endpoints. This overlap was negligible relative to the durations of the windows and is unlikely to account for the consistent regional findings. Nevertheless, the windows were not completely disjoint, and this detail should be considered when interpreting extremely precise distinctions at their boundaries.

The aperiodic-only model was an intentionally restricted stress test. It fitted the overall spectra less effectively than the two-second max6 model, as would be expected when genuine periodic structure was present but prohibited. Its similar condition effect supports independence from a particular peak-fitting solution, but the aperiodic-only model should not be presented as the preferred generative account of the spectra.

Finally, the other sensor category contained channels that could not be assigned clearly to one of the predefined anatomical sensor groups. Although this category produced several statistically corrected effects, it did not correspond to a coherent anatomical distribution. Its results were retained to preserve the complete statistical family but should not be emphasized as evidence for a specific neuroanatomical process.

## 5 Conclusion

Human-opponent and ball-machine table-tennis play were associated with different peri-hit EEG spectral structures. Nuisance-residualized alpha and beta power was generally lower during ball-machine play, while the preferred spectral-parameterization analysis showed a higher ball-machine aperiodic exponent, particularly over posterior and parietal sensors. The posterior post-hit effect was the strongest standardized anatomically interpretable contrast and showed high participant-level direction consistency.

Aligning the lower peak-width limit with short-window frequency resolution did not remove the focal parietal effect. The difference remained under the resolution-aware max6 model, the reduced-complexity max3 model, the better-resolved two-second max6 model, and the aperiodic-only sensitivity analysis. Each model produced a positive bootstrap interval, and every leave-one-participant-out analysis retained a positive and FDR-significant result.

Taken together, the primary analysis and multiple sensitivity analyses indicate that the two playing contexts were associated with a robust difference in broadband aperiodic spectral structure. The analyses do not establish that ball-machine play caused a particular cognitive state, nor do they identify a unique physiological mechanism. The findings instead demonstrate that event-locked mobile EEG can distinguish naturalistic visuomotor contexts when movement-related nuisance signals, spectral resolution, model complexity, fit quality, and participant influence are examined explicitly.

More broadly, the findings show how mobile EEG may be useful not only for establishing whether naturalistic task contexts differ, but also for characterizing the form of that difference when a practice environment is not identical to the activity for which performance is ultimately being trained. Combined with behavioral measures, this type of analysis could allow future studies to ask whether a training context engages the processes that are intended to be trained and whether that pattern differs across individuals. The principle may be relevant not only to sport but also to simulation, rehabilitation, education, and other settings in which practice provides an approximation of later performance. Such applications will require longitudinal and out-of-sample validation, but the present study provides a methodological basis for investigating them.

