## Supplementary Material for "Resolution-aware spectral parameterization reveals robust context-dependent aperiodic EEG dynamics during real-world table tennis"

### Supplementary Figures

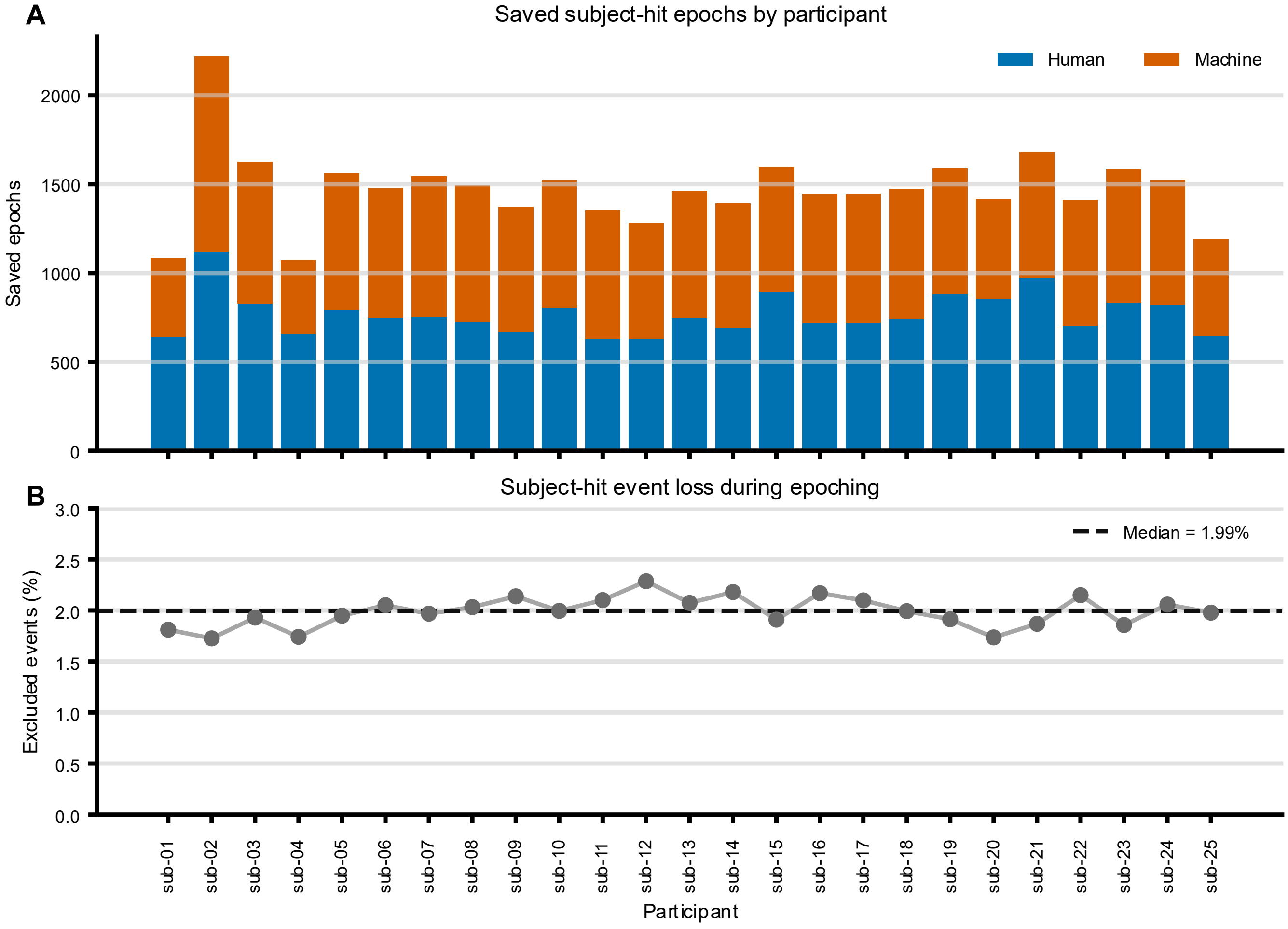


Supplementary Figure S1. Participant-level epoch retention. A, saved human-opponent and ball-machine participant-hit epochs for each participant. B, percentage of candidate participant-hit events excluded during epoching; a zero-based event-loss scale is used instead of a truncated 0–100% retention scale. The dashed horizontal line indicates the group meadian event loss.

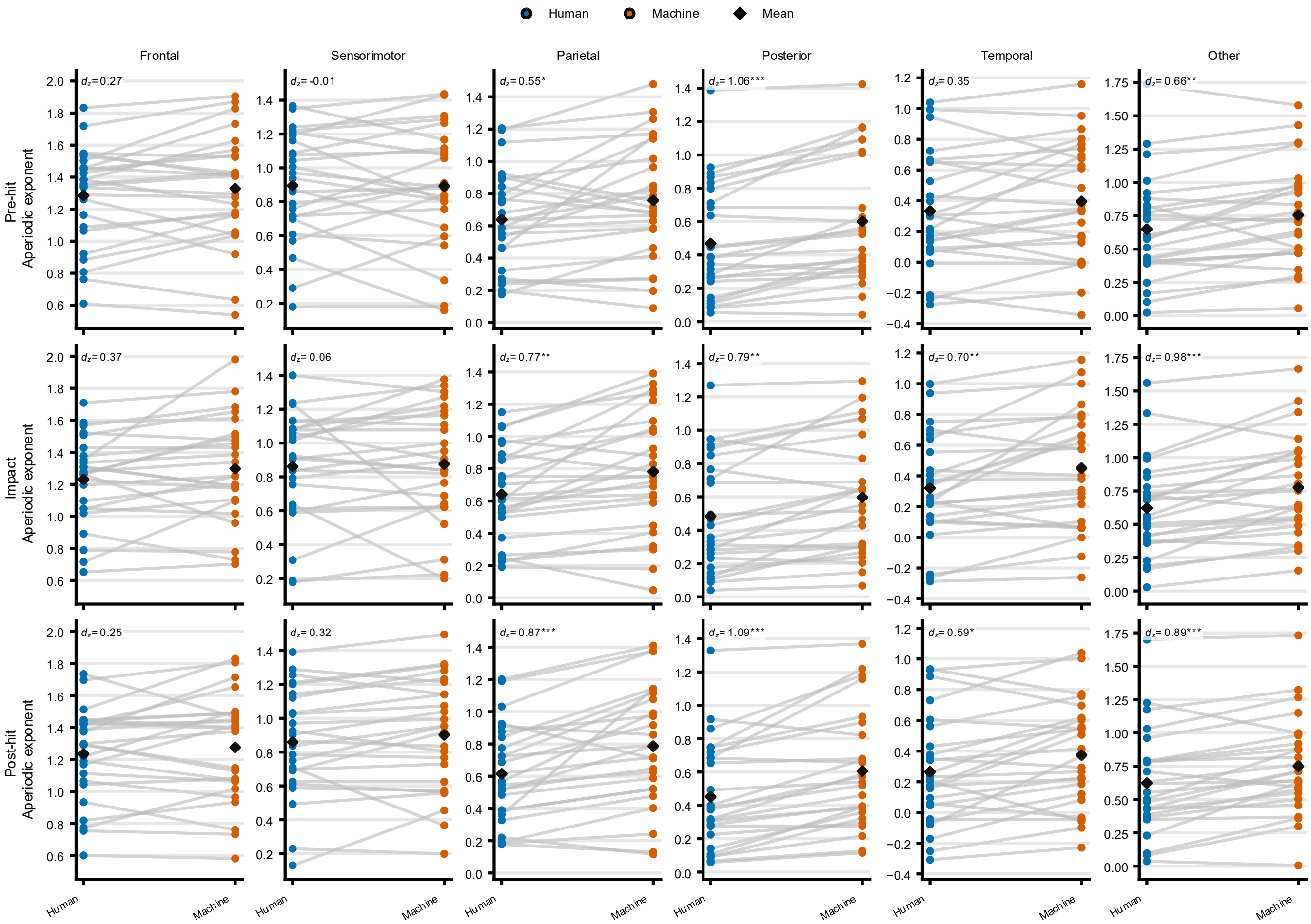


Supplementary Figure S2. Complete short-window participant distributions. Paired participant-level aperiodic-exponent values for all 18 combinations of three peri-hit windows and six sensor regions from the preferred resolution-aware max6 model. Rows correspond to pre-hit, impact, and post-hit windows, and columns correspond to sensor regions. Connecting lines link human-opponent and ball-machine values within participants; diamonds show condition means. Each sensor-region column uses a common y-axis range across the three windows. Panel annotations report Cohen's dz, with asterisks indicating Benjamini–Hochberg-adjusted paired t-test values within the fixed 18-test exponent family (*q $<$ 0.05, **q $<$ 0.01, ***q $<$ 0.001). n = 25 participants.

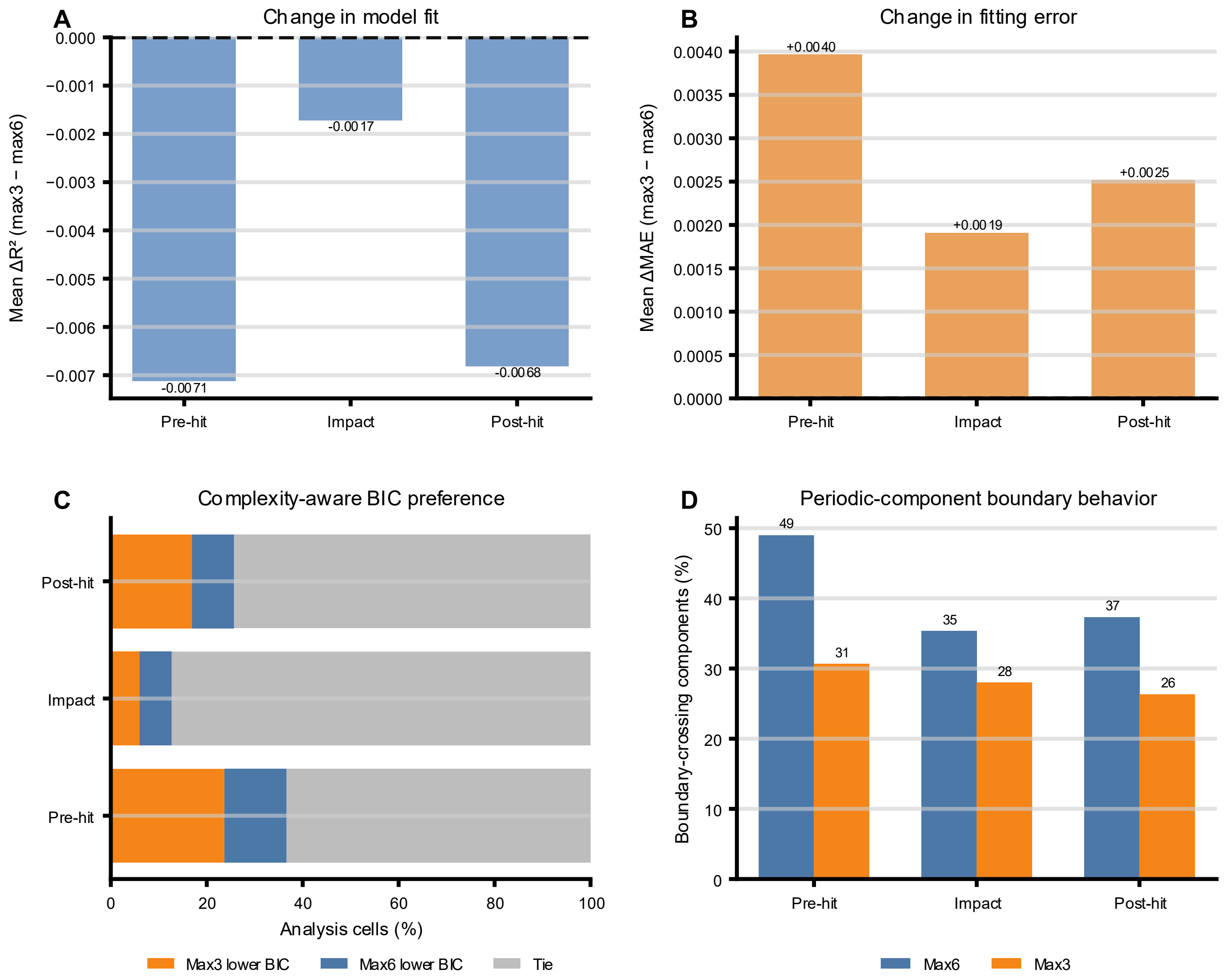


Supplementary Figure S3. Spectral-model quality and max6–max3 model selection. A, mean change in $R^2$ when the maximum number of periodic peaks was reduced from six to three (max3 − max6). B, corresponding mean change in mean absolute error. C, percentages of model cells favoring max3 or max6 according to the Bayesian information criterion; lower BIC indicates the preferred model. D, percentages of periodic components crossing the predefined frequency-band boundaries in max6 and max3 fits. Summaries are shown by short analysis window. These are descriptive model diagnostics based on paired model cells and are not independent-cell inferential tests. BIC = Bayesian information criterion; MAE = mean absolute error.

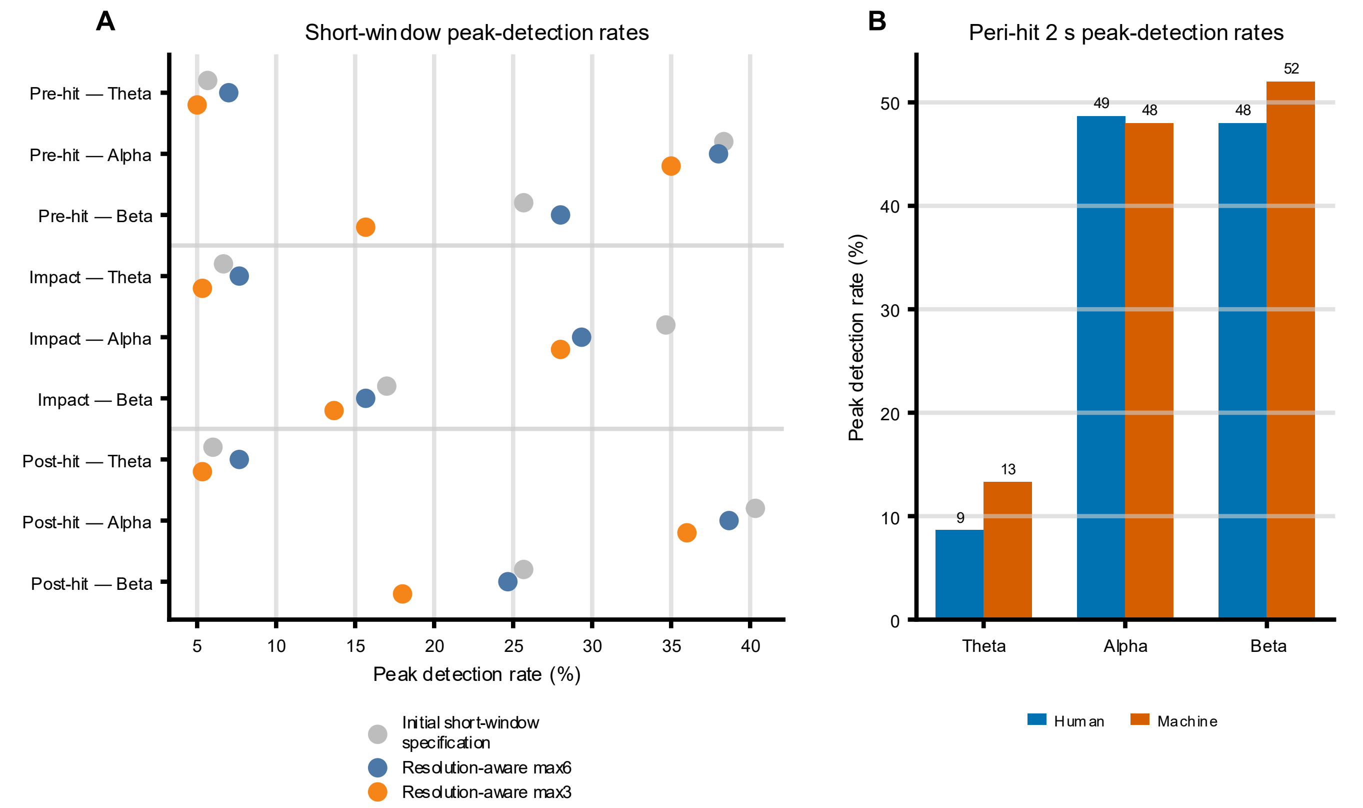


Supplementary Figure S4. Peak-detection sensitivity. A, theta-, alpha-, and beta-peak detection rates across the initial short-window specification, resolution-aware max6, and resolution-aware max3 short-window models. B, human-opponent and ball-machine peak-detection rates for the two-second peri-hit max6 model. Rates denote the percentage of fitted model cells in which a periodic peak was detected in the corresponding band. The aperiodic-only model is not plotted as an observed zero because periodic components were disabled by design.

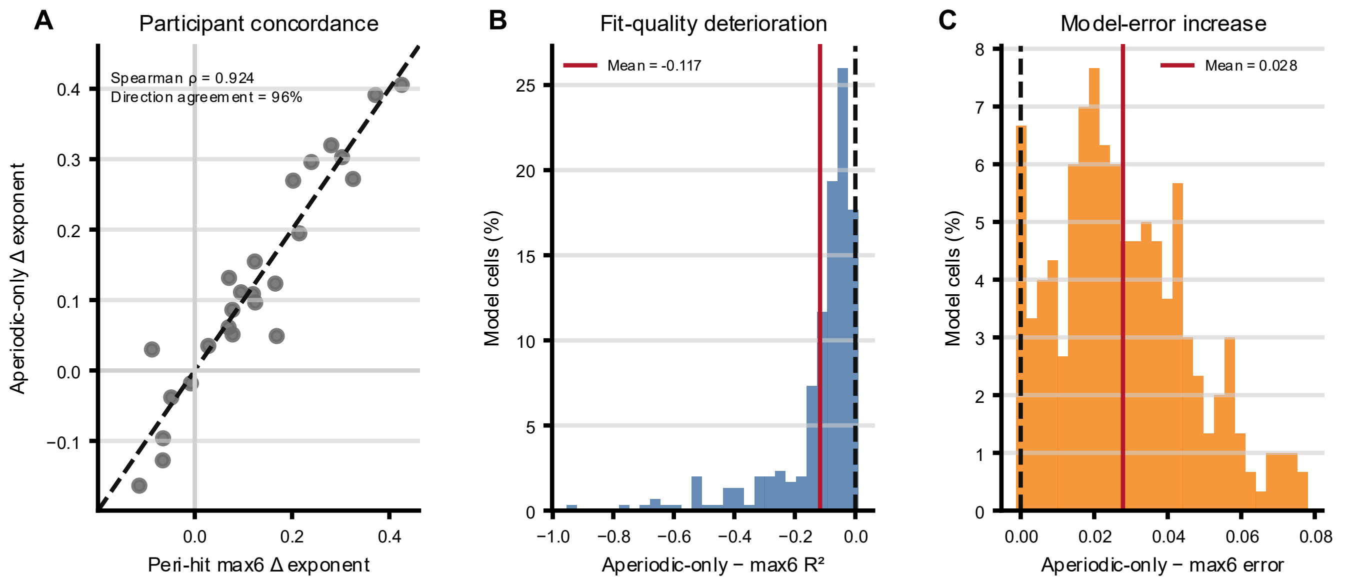


Supplementary Figure S5. Aperiodic-only stress test. A, participant-level concordance between parietal machine-minus-human exponent effects from the two-second peri-hit max6 and aperiodic-only models. The dashed line is the identity line; Spearman ρ and directional agreement are shown in the panel. B, distribution of cell-level changes in $R^2$

after periodic peaks were disabled (aperiodic-only − max6). C, corresponding distribution of changes in model error. Vertical dashed lines indicate zero and solid lines indicate the mean change. Histograms show percentages of paired model cells. Cell-level summaries are descriptive and are not treated as independent inferential observations.

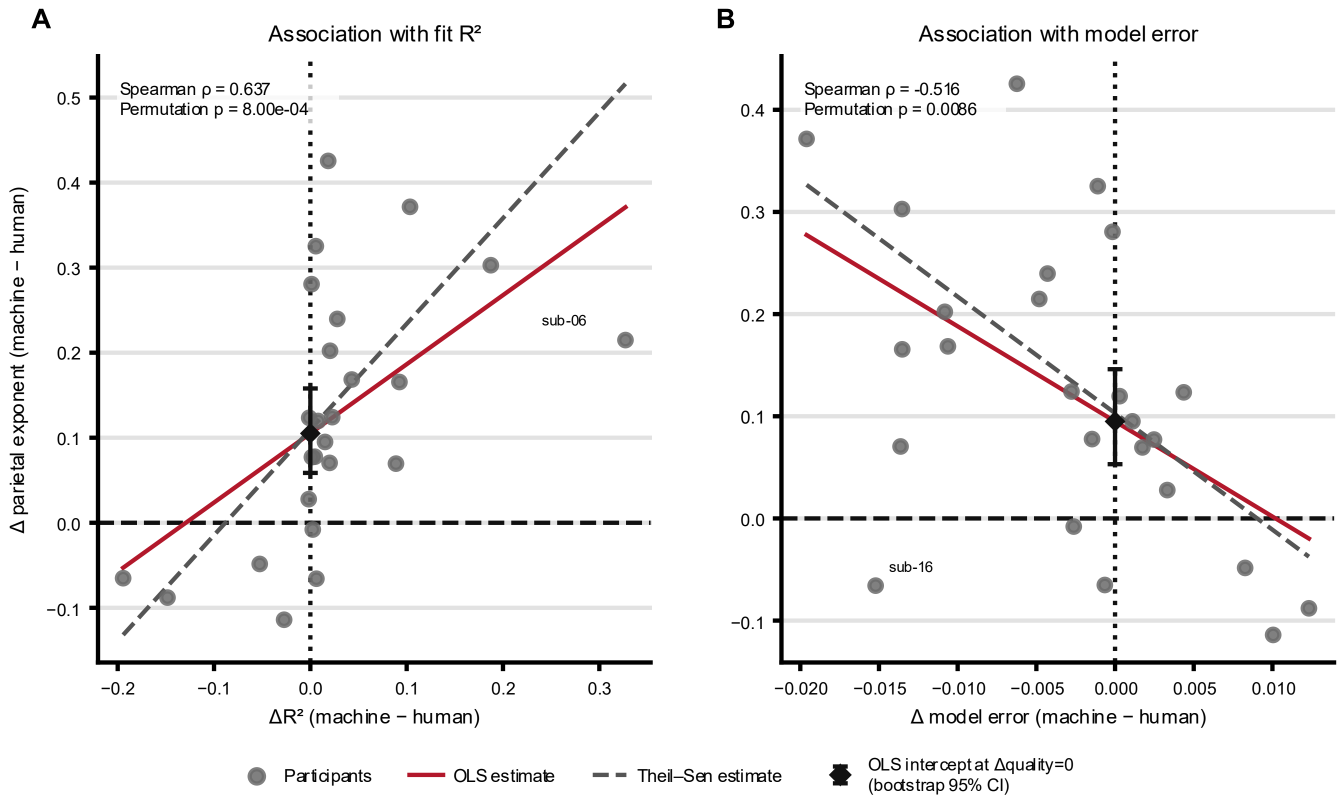


Supplementary Figure S6. Two-second fit-quality influence audit. Association between the participant-level parietal machine-minus-human aperiodic-exponent difference and the corresponding machine-minus-human difference in $R^2$ (A) or spectral-model error (B) for the two-second peri-hit max6 model. Points represent participants. Solid lines show ordinary least-squares estimates and dashed lines show Theil–Sen estimates. The diamond at a fit-quality difference of zero is the OLS intercept with a paired-bootstrap 95% confidence interval. Spearman $\rho$ and permutation p values are reported in each panel. Participant labels identify observations with Cook's distance greater than $4/n$. These analyses are exploratory because the exponent and fit-quality metrics are outputs of the same spectral model.

### Supplementary tables

Supplementary Table S1. Per-participant epoch counts and retention. Candidate participant-hit events, saved epochs, condition-specific epoch counts, and retention proportions for each participant.

| Subject | Candidate subject-hit events | Saved epochs | Human-opponent epochs | Ball-machine epochs | Retention (%) | Human proportion of saved epochs (%) | Ball-machine proportion of saved epochs (%) |
|---|---|---|---|---|---|---|---|
| Participant 01 | 1104 | 1084 | 641 | 443 | 98.2 | 59.133 | 40.867 |
| Participant 02 | 2259 | 2220 | 1119 | 1101 | 98.3 | 50.405 | 49.595 |
| Participant 03 | 1658 | 1626 | 829 | 797 | 98.1 | 50.984 | 49.016 |
| Participant 04 | 1091 | 1072 | 658 | 414 | 98.3 | 61.381 | 38.619 |
| Participant 05 | 1591 | 1560 | 790 | 770 | 98.1 | 50.641 | 49.359 |
| Participant 06 | 1511 | 1480 | 750 | 730 | 97.9 | 50.676 | 49.324 |
| Participant 07 | 1575 | 1544 | 752 | 792 | 98.0 | 48.705 | 51.295 |
| Participant 08 | 1525 | 1494 | 723 | 771 | 98.0 | 48.394 | 51.606 |

| Subject | Candidate subject-hit events | Saved epochs | Human-opponent epochs | Ball-machine epochs | Retention (%) | Human proportion of saved epochs (%) | Ball-machine proportion of saved epochs (%) |
|---|---|---|---|---|---|---|---|
| Participant 09 | 1403 | 1373 | 669 | 704 | 97.9 | 48.725 | 51.275 |
| Participant 10 | 1553 | 1522 | 804 | 718 | 98.0 | 52.825 | 47.175 |
| Participant 11 | 1380 | 1351 | 628 | 723 | 97.9 | 46.484 | 53.516 |
| Participant 12 | 1312 | 1282 | 632 | 650 | 97.7 | 49.298 | 50.702 |
| Participant 13 | 1495 | 1464 | 747 | 717 | 97.9 | 51.025 | 48.975 |
| Participant 14 | 1422 | 1391 | 690 | 701 | 97.8 | 49.605 | 50.395 |
| Participant 15 | 1623 | 1592 | 893 | 699 | 98.1 | 56.093 | 43.907 |
| Participant 16 | 1475 | 1443 | 717 | 726 | 97.8 | 49.688 | 50.312 |
| Participant 17 | 1477 | 1446 | 720 | 726 | 97.9 | 49.793 | 50.207 |
| Participant 18 | 1505 | 1475 | 740 | 735 | 98.0 | 50.169 | 49.831 |
| Participant 19 | 1620 | 1589 | 880 | 709 | 98.1 | 55.381 | 44.619 |
| Participant 20 | 1440 | 1415 | 854 | 561 | 98.3 | 60.353 | 39.647 |
| Participant 21 | 1712 | 1680 | 969 | 711 | 98.1 | 57.679 | 42.321 |
| Participant 22 | 1442 | 1411 | 703 | 708 | 97.8 | 49.823 | 50.177 |
| Participant 23 | 1615 | 1585 | 834 | 751 | 98.1 | 52.618 | 47.382 |
| Participant 24 | 1555 | 1523 | 823 | 700 | 97.9 | 54.038 | 45.962 |
| Participant 25 | 1213 | 1189 | 647 | 542 | 98.0 | 54.415 | 45.585 |

*Note.* Retention (%) = 100 × saved epochs / candidate participant-hit events. The human-opponent and ball-machine proportions are calculated relative to saved epochs within each participant.

Supplementary Table S2. Full residualized-bandpower statistics. Complete residualized-bandpower statistics across all windows, canonical bands, and sensor regions.

| Window | Band | Sensor region | Participants | Human residualized mean | Machine residualized mean | Machine − human residualized mean | Difference SD | Paired t | Paired t p | Paired t FDR q | Wilcoxon p | Wilcoxon FDR q | Cohen's dz |
|---|---|---|---|---|---|---|---|---|---|---|---|---|---|
| Pre-hit | Alpha | Posterior | 25 | 0.0340 | -0.0308 | -0.0647 | 0.0384 | -8.4371 | <0.001 | <0.001 | <0.001 | <0.001 | -1.6874 |
| Impact | Beta | Posterior | 25 | 0.0418 | -0.0344 | -0.0762 | 0.0481 | -7.9217 | <0.001 | <0.001 | <0.001 | <0.001 | -1.5843 |
| Pre-hit | Beta | Posterior | 25 | 0.0412 | -0.0349 | -0.0761 | 0.0471 | -8.0877 | <0.001 | <0.001 | <0.001 | <0.001 | -1.6175 |
| Impact | Beta | Frontal | 25 | 0.0285 | -0.0250 | -0.0536 | 0.0356 | -7.5199 | <0.001 | <0.001 | <0.001 | <0.001 | -1.5040 |
| Post-hit | Alpha | Posterior | 25 | 0.0309 | -0.0282 | -0.0590 | 0.0388 | -7.6111 | <0.001 | <0.001 | <0.001 | <0.001 | -1.5222 |
| Impact | Beta | Parietal | 25 | 0.0269 | -0.0277 | -0.0546 | 0.0383 | -7.1391 | <0.001 | <0.001 | <0.001 | <0.001 | -1.4278 |
| Pre-hit | Beta | Sensorimotor | 25 | 0.0220 | -0.0241 | -0.0462 | 0.0339 | -6.8079 | <0.001 | <0.001 | <0.001 | <0.001 | -1.3616 |
| Pre-hit | Beta | Frontal | 25 | 0.0274 | -0.0245 | -0.0519 | 0.0387 | -6.6964 | <0.001 | <0.001 | <0.001 | <0.001 | -1.3393 |
| Pre-hit | Beta | Parietal | 25 | 0.0250 | -0.0276 | -0.0526 | 0.0392 | -6.7182 | <0.001 | <0.001 | <0.001 | <0.001 | -1.3436 |
| Post-hit | Beta | Posterior | 25 | 0.0372 | -0.0315 | -0.0687 | 0.0526 | -6.5349 | <0.001 | <0.001 | <0.001 | <0.001 | -1.3070 |
| Impact | Beta | Sensorimotor | 25 | 0.0217 | -0.0224 | -0.0441 | 0.0337 | -6.5557 | <0.001 | <0.001 | <0.001 | <0.001 | -1.3111 |
| Impact | Alpha | Posterior | 25 | 0.0311 | -0.0272 | -0.0583 | 0.0464 | -6.2905 | <0.001 | <0.001 | <0.001 | <0.001 | -1.2581 |
| Post-hit | Beta | Frontal | 25 | 0.0261 | -0.0224 | -0.0485 | 0.0393 | -6.1649 | <0.001 | <0.001 | <0.001 | <0.001 | -1.2330 |
| Pre-hit | Alpha | Parietal | 25 | 0.0145 | -0.0192 | -0.0336 | 0.0277 | -6.0615 | <0.001 | <0.001 | <0.001 | <0.001 | -1.2123 |
| Post-hit | Beta | Sensorimotor | 25 | 0.0214 | -0.0223 | -0.0437 | 0.0365 | -5.9894 | <0.001 | <0.001 | <0.001 | <0.001 | -1.1979 |
| Post-hit | Beta | Parietal | 25 | 0.0241 | -0.0262 | -0.0503 | 0.0439 | -5.7326 | <0.001 | <0.001 | <0.001 | <0.001 | -1.1465 |
| Pre-hit | Alpha | Frontal | 25 | 0.0181 | -0.0192 | -0.0374 | 0.0338 | -5.5282 | <0.001 | <0.001 | <0.001 | <0.001 | -1.1056 |
| Post-hit | Alpha | Other | 25 | 0.0372 | -0.0241 | -0.0613 | 0.0558 | -5.4937 | <0.001 | <0.001 | <0.001 | <0.001 | -1.0987 |
| Pre-hit | Theta | Temporal | 25 | 0.0193 | -0.0173 | -0.0365 | 0.0359 | -5.0856 | <0.001 | <0.001 | <0.001 | <0.001 | -1.0171 |
| Impact | Alpha | Parietal | 25 | 0.0138 | -0.0178 | -0.0316 | 0.0313 | -5.0509 | <0.001 | <0.001 | <0.001 | <0.001 | -1.0102 |
| Impact | Beta | Temporal | 25 | 0.0402 | -0.0319 | -0.0721 | 0.0734 | -4.9122 | <0.001 | <0.001 | <0.001 | <0.001 | -0.9824 |
| Pre-hit | Theta | Frontal | 25 | 0.0226 | -0.0172 | -0.0398 | 0.0405 | -4.9100 | <0.001 | <0.001 | <0.001 | <0.001 | -0.9820 |
| Pre-hit | Beta | Temporal | 25 | 0.0396 | -0.0326 | -0.0722 | 0.0741 | -4.8711 | <0.001 | <0.001 | <0.001 | <0.001 | -0.9742 |
| Post-hit | Alpha | Frontal | 25 | 0.0172 | -0.0181 | -0.0353 | 0.0364 | -4.8405 | <0.001 | <0.001 | <0.001 | <0.001 | -0.9681 |
| Pre-hit | Alpha | Other | 25 | 0.0400 | -0.0268 | -0.0668 | 0.0704 | -4.7476 | <0.001 | <0.001 | <0.001 | <0.001 | -0.9495 |
| Impact | Alpha | Other | 25 | 0.0390 | -0.0241 | -0.0632 | 0.0680 | -4.6450 | <0.001 | <0.001 | <0.001 | <0.001 | -0.9290 |
| Post-hit | Beta | Temporal | 25 | 0.0385 | -0.0309 | -0.0694 | 0.0756 | -4.5914 | <0.001 | <0.001 | <0.001 | <0.001 | -0.9183 |
| Pre-hit | Beta | Other | 25 | 0.0418 | -0.0297 | -0.0715 | 0.0787 | -4.5412 | <0.001 | <0.001 | <0.001 | <0.001 | -0.9082 |
| Pre-hit | Theta | Parietal | 25 | 0.0116 | -0.0096 | -0.0212 | 0.0233 | -4.5479 | <0.001 | <0.001 | <0.001 | <0.001 | -0.9096 |
| Impact | Alpha | Frontal | 25 | 0.0170 | -0.0174 | -0.0344 | 0.0385 | -4.4654 | <0.001 | <0.001 | <0.001 | <0.001 | -0.8931 |
| Impact | Beta | Other | 25 | 0.0411 | -0.0285 | -0.0697 | 0.0791 | -4.4068 | <0.001 | <0.001 | <0.001 | <0.001 | -0.8814 |
| Post-hit | Alpha | Parietal | 25 | 0.0135 | -0.0176 | -0.0311 | 0.0354 | -4.3935 | <0.001 | <0.001 | <0.001 | <0.001 | -0.8787 |
| Post-hit | Beta | Other | 25 | 0.0376 | -0.0261 | -0.0637 | 0.0781 | -4.0759 | <0.001 | <0.001 | <0.001 | <0.001 | -0.8152 |
| Pre-hit | Alpha | Temporal | 25 | 0.0139 | -0.0175 | -0.0314 | 0.0422 | -3.7232 | 0.001 | 0.002 | 0.001 | 0.002 | -0.7446 |
| Post-hit | Alpha | Temporal | 25 | 0.0132 | -0.0177 | -0.0309 | 0.0415 | -3.7276 | 0.001 | 0.002 | <0.001 | <0.001 | -0.7455 |
| Impact | Theta | Temporal | 25 | 0.0181 | -0.0155 | -0.0336 | 0.0452 | -3.7141 | 0.001 | 0.002 | <0.001 | 0.001 | -0.7428 |
| Impact | Alpha | Temporal | 25 | 0.0128 | -0.0159 | -0.0287 | 0.0396 | -3.6181 | 0.001 | 0.002 | 0.002 | 0.003 | -0.7236 |
| Pre-hit | Theta | Posterior | 25 | 0.0115 | -0.0133 | -0.0248 | 0.0358 | -3.4637 | 0.002 | 0.003 | 0.004 | 0.006 | -0.6927 |
| Pre-hit | Theta | Other | 25 | 0.0217 | -0.0152 | -0.0369 | 0.0569 | -3.2426 | 0.003 | 0.005 | 0.004 | 0.005 | -0.6485 |
| Pre-hit | Alpha | Sensorimotor | 25 | 0.0070 | -0.0124 | -0.0193 | 0.0305 | -3.1655 | 0.004 | 0.006 | 0.006 | 0.008 | -0.6331 |
| Impact | Theta | Posterior | 25 | 0.0100 | -0.0111 | -0.0211 | 0.0373 | -2.8310 | 0.009 | 0.012 | 0.011 | 0.014 | -0.5662 |
| Post-hit | Alpha | Sensorimotor | 25 | 0.0067 | -0.0126 | -0.0192 | 0.0339 | -2.8408 | 0.009 | 0.012 | 0.008 | 0.011 | -0.5682 |
| Impact | Theta | Frontal | 25 | 0.0191 | -0.0120 | -0.0311 | 0.0558 | -2.7860 | 0.010 | 0.013 | 0.011 | 0.014 | -0.5572 |
| Post-hit | Theta | Temporal | 25 | 0.0146 | -0.0119 | -0.0265 | 0.0522 | -2.5417 | 0.018 | 0.022 | 0.032 | 0.037 | -0.5083 |
| Pre-hit | Theta | Sensorimotor | 25 | 0.0062 | -0.0077 | -0.0139 | 0.0275 | -2.5233 | 0.019 | 0.022 | 0.012 | 0.015 | -0.5047 |
| Impact | Theta | Parietal | 25 | 0.0101 | -0.0076 | -0.0176 | 0.0352 | -2.5022 | 0.020 | 0.023 | 0.016 | 0.019 | -0.5004 |
| Post-hit | Theta | Frontal | 25 | 0.0158 | -0.0091 | -0.0249 | 0.0502 | -2.4867 | 0.020 | 0.023 | 0.045 | 0.052 | -0.4973 |
| Impact | Alpha | Sensorimotor | 25 | 0.0046 | -0.0097 | -0.0143 | 0.0331 | -2.1588 | 0.041 | 0.046 | 0.075 | 0.083 | -0.4318 |

| Window | Band | Sensor region | Participants | Human residualized mean | Machine residualized mean | Machine − human residualized mean | Difference SD | Paired t | Paired t p | Paired t FDR q | Wilcoxon p | Wilcoxon FDR q | Cohen's dz |
|---|---|---|---|---|---|---|---|---|---|---|---|---|---|
| Impact | Theta | Other | 25 | 0.0162 | -0.0079 | -0.0240 | 0.0574 | -2.0936 | 0.047 | 0.052 | 0.063 | 0.070 | -0.4187 |
| Post-hit | Theta | Other | 25 | 0.0140 | -0.0049 | -0.0189 | 0.0542 | -1.7414 | 0.094 | 0.102 | 0.101 | 0.107 | -0.3483 |
| Impact | Theta | Sensorimotor | 25 | 0.0056 | -0.0058 | -0.0114 | 0.0343 | -1.6641 | 0.109 | 0.116 | 0.101 | 0.107 | -0.3328 |
| Post-hit | Theta | Sensorimotor | 25 | 0.0035 | -0.0042 | -0.0077 | 0.0307 | -1.2495 | 0.224 | 0.232 | 0.191 | 0.198 | -0.2499 |
| Post-hit | Theta | Parietal | 25 | 0.0051 | -0.0021 | -0.0072 | 0.0376 | -0.9590 | 0.347 | 0.354 | 0.458 | 0.466 | -0.1918 |
| Post-hit | Theta | Posterior | 25 | 0.0024 | -0.0026 | -0.0050 | 0.0450 | -0.5533 | 0.585 | 0.585 | 0.578 | 0.578 | -0.1107 |

*Note.* Human and machine means are participant-level means of nuisance-residualized log power. Machine − human values below zero indicate lower residualized power during ball-machine play. FDR correction was applied across the complete 54-test family separately for paired t tests and Wilcoxon signed-rank tests.

Supplementary Table S3. Full short-window aperiodic-exponent and offset statistics. Complete participant-level statistics for the preferred resolution-aware max6 model. The table is divided into estimate and inferential panels for each aperiodic feature.

Aperiodic exponent: effect estimates

| Window | Sensor region | Participants | Human mean | Human median | Machine mean | Machine median | Machine − human mean | Machine − human median | Difference SD | Cohen's dz |
|---|---|---|---|---|---|---|---|---|---|---|
| Pre-hit | Frontal | 25 | 1.2850 | 1.3571 | 1.3286 | 1.4069 | 0.0436 | 0.0718 | 0.1592 | 0.2738 |
| Pre-hit | Sensorimotor | 25 | 0.8950 | 0.9378 | 0.8923 | 0.8870 | -0.0026 | 0.0195 | 0.1981 | -0.0134 |
| Pre-hit | Parietal | 25 | 0.6386 | 0.6346 | 0.7577 | 0.7154 | 0.1191 | 0.0897 | 0.2158 | 0.5520 |
| Pre-hit | Posterior | 25 | 0.4689 | 0.3836 | 0.6018 | 0.5244 | 0.1329 | 0.1223 | 0.1253 | 1.0604 |
| Pre-hit | Temporal | 25 | 0.3322 | 0.2970 | 0.3959 | 0.3443 | 0.0637 | 0.0720 | 0.1839 | 0.3462 |
| Pre-hit | Other | 25 | 0.6491 | 0.5779 | 0.7552 | 0.7201 | 0.1061 | 0.1097 | 0.1598 | 0.6638 |
| Impact | Frontal | 25 | 1.2304 | 1.2702 | 1.2977 | 1.2523 | 0.0673 | 0.0555 | 0.1837 | 0.3664 |
| Impact | Sensorimotor | 25 | 0.8616 | 0.9092 | 0.8755 | 0.8921 | 0.0140 | 0.0322 | 0.2179 | 0.0642 |
| Impact | Parietal | 25 | 0.6416 | 0.6148 | 0.7825 | 0.7533 | 0.1409 | 0.0989 | 0.1836 | 0.7677 |
| Impact | Posterior | 25 | 0.4835 | 0.3573 | 0.5968 | 0.5507 | 0.1132 | 0.0962 | 0.1436 | 0.7884 |
| Impact | Temporal | 25 | 0.3197 | 0.2615 | 0.4505 | 0.4035 | 0.1308 | 0.0913 | 0.1864 | 0.7021 |
| Impact | Other | 25 | 0.6220 | 0.5624 | 0.7765 | 0.7526 | 0.1544 | 0.1251 | 0.1583 | 0.9755 |
| Post-hit | Frontal | 25 | 1.2349 | 1.2937 | 1.2760 | 1.3950 | 0.0410 | 0.0368 | 0.1643 | 0.2499 |
| Post-hit | Sensorimotor | 25 | 0.8590 | 0.8661 | 0.9015 | 0.9523 | 0.0425 | 0.0482 | 0.1337 | 0.3179 |
| Post-hit | Parietal | 25 | 0.6148 | 0.5676 | 0.7863 | 0.7904 | 0.1715 | 0.1338 | 0.1977 | 0.8674 |
| Post-hit | Posterior | 25 | 0.4512 | 0.3746 | 0.6058 | 0.5396 | 0.1547 | 0.1311 | 0.1416 | 1.0926 |
| Post-hit | Temporal | 25 | 0.2655 | 0.2060 | 0.3741 | 0.3660 | 0.1086 | 0.1115 | 0.1843 | 0.5894 |
| Post-hit | Other | 25 | 0.6226 | 0.5532 | 0.7498 | 0.7104 | 0.1272 | 0.1334 | 0.1422 | 0.8944 |

Aperiodic exponent: inferential statistics and direction consistency

| Window | Sensor region | Paired t | Paired t p | Paired t FDR q | Wilcoxon p | Wilcoxon FDR q | Machine lower, n (%) | Machine higher, n (%) | Equal, n |
|---|---|---|---|---|---|---|---|---|---|
| Pre-hit | Frontal | 1.3692 | 0.184 | 0.220 | 0.182 | 0.218 | 11 (44.0) | 14 (56.0) | 0 |
| Pre-hit | Sensorimotor | -0.0668 | 0.947 | 0.947 | 0.958 | 0.958 | 10 (40.0) | 15 (60.0) | 0 |
| Pre-hit | Parietal | 2.7600 | 0.011 | 0.018 | 0.020 | 0.033 | 6 (24.0) | 19 (76.0) | 0 |
| Pre-hit | Posterior | 5.3018 | <0.001 | <0.001 | <0.001 | <0.001 | 4 (16.0) | 21 (84.0) | 0 |
| Pre-hit | Temporal | 1.7311 | 0.096 | 0.133 | 0.127 | 0.163 | 10 (40.0) | 15 (60.0) | 0 |
| Pre-hit | Other | 3.3191 | 0.003 | 0.006 | 0.002 | 0.004 | 6 (24.0) | 19 (76.0) | 0 |
| Impact | Frontal | 1.8322 | 0.079 | 0.119 | 0.067 | 0.100 | 8 (32.0) | 17 (68.0) | 0 |

| Window | Sensor region | Paired t | Paired t p | Paired t FDR q | Wilcoxon p | Wilcoxon FDR q | Machine lower, n (%) | Machine higher, n (%) | Equal, n |
|---|---|---|---|---|---|---|---|---|---|
| Impact | Sensorimotor | 0.3208 | 0.751 | 0.795 | 0.200 | 0.225 | 9 (36.0) | 16 (64.0) | 0 |
| Impact | Parietal | 3.8384 | <0.001 | 0.002 | <0.001 | 0.002 | 5 (20.0) | 20 (80.0) | 0 |
| Impact | Posterior | 3.9421 | <0.001 | 0.002 | <0.001 | <0.001 | 4 (16.0) | 21 (84.0) | 0 |
| Impact | Temporal | 3.5103 | 0.002 | 0.004 | 0.002 | 0.004 | 5 (20.0) | 20 (80.0) | 0 |
| Impact | Other | 4.8774 | <0.001 | <0.001 | <0.001 | <0.001 | 3 (12.0) | 22 (88.0) | 0 |
| Post-hit | Frontal | 1.2494 | 0.224 | 0.252 | 0.312 | 0.331 | 11 (44.0) | 14 (56.0) | 0 |
| Post-hit | Sensorimotor | 1.5894 | 0.125 | 0.161 | 0.080 | 0.111 | 10 (40.0) | 15 (60.0) | 0 |
| Post-hit | Parietal | 4.3369 | <0.001 | <0.001 | <0.001 | <0.001 | 4 (16.0) | 21 (84.0) | 0 |
| Post-hit | Posterior | 5.4628 | <0.001 | <0.001 | <0.001 | <0.001 | 2 (8.0) | 23 (92.0) | 0 |
| Post-hit | Temporal | 2.9470 | 0.007 | 0.013 | 0.011 | 0.019 | 7 (28.0) | 18 (72.0) | 0 |
| Post-hit | Other | 4.4721 | <0.001 | <0.001 | <0.001 | <0.001 | 3 (12.0) | 22 (88.0) | 0 |

Aperiodic offset: effect estimates

| Window | Sensor region | Participants | Human mean | Human median | Machine mean | Machine median | Machine − human mean | Machine − human median | Difference SD | Cohen's dz |
|---|---|---|---|---|---|---|---|---|---|---|
| Pre-hit | Frontal | 25 | -10.4030 | -10.3082 | -10.4295 | -10.4321 | -0.0266 | 0.0038 | 0.1831 | -0.1451 |
| Pre-hit | Sensorimotor | 25 | -11.2693 | -11.2572 | -11.3599 | -11.2849 | -0.0906 | -0.0133 | 0.2558 | -0.3542 |
| Pre-hit | Parietal | 25 | -11.2723 | -11.2514 | -11.2195 | -11.2257 | 0.0528 | 0.0415 | 0.2206 | 0.2393 |
| Pre-hit | Posterior | 25 | -11.0698 | -11.0967 | -11.0288 | -11.0861 | 0.0410 | 0.0505 | 0.1155 | 0.3555 |
| Pre-hit | Temporal | 25 | -11.3114 | -11.3453 | -11.3414 | -11.4480 | -0.0300 | -0.0159 | 0.1959 | -0.1533 |
| Pre-hit | Other | 25 | -10.7380 | -10.7600 | -10.7273 | -10.7476 | 0.0107 | 0.0066 | 0.1873 | 0.0571 |
| Impact | Frontal | 25 | -10.4523 | -10.5005 | -10.4432 | -10.5011 | 0.0091 | -0.0141 | 0.2402 | 0.0380 |
| Impact | Sensorimotor | 25 | -11.2939 | -11.2984 | -11.3615 | -11.2843 | -0.0676 | 0.0087 | 0.2798 | -0.2418 |
| Impact | Parietal | 25 | -11.2656 | -11.2750 | -11.1788 | -11.1342 | 0.0868 | 0.0362 | 0.1851 | 0.4690 |
| Impact | Posterior | 25 | -11.0542 | -11.0592 | -11.0295 | -11.0495 | 0.0247 | 0.0097 | 0.1318 | 0.1876 |
| Impact | Temporal | 25 | -11.3205 | -11.3728 | -11.2567 | -11.2837 | 0.0638 | 0.0277 | 0.2401 | 0.2656 |
| Impact | Other | 25 | -10.7682 | -10.7595 | -10.6873 | -10.7066 | 0.0809 | 0.0634 | 0.2000 | 0.4045 |
| Post-hit | Frontal | 25 | -10.4507 | -10.4143 | -10.4910 | -10.5522 | -0.0403 | -0.0204 | 0.2086 | -0.1930 |
| Post-hit | Sensorimotor | 25 | -11.3103 | -11.3026 | -11.3394 | -11.2706 | -0.0291 | 0.0188 | 0.1676 | -0.1738 |
| Post-hit | Parietal | 25 | -11.3092 | -11.3246 | -11.1917 | -11.1349 | 0.1175 | 0.0920 | 0.2083 | 0.5643 |
| Post-hit | Posterior | 25 | -11.0994 | -11.0922 | -11.0277 | -11.0919 | 0.0717 | 0.0700 | 0.1505 | 0.4766 |
| Post-hit | Temporal | 25 | -11.3971 | -11.4836 | -11.3723 | -11.4836 | 0.0248 | 0.0535 | 0.1952 | 0.1270 |
| Post-hit | Other | 25 | -10.7822 | -10.7766 | -10.7462 | -10.7480 | 0.0360 | 0.0325 | 0.1681 | 0.2141 |

Aperiodic offset: inferential statistics and direction consistency

| Window | Sensor region | Paired t | Paired t p | Paired t FDR q | Wilcoxon p | Wilcoxon FDR q | Machine lower, n (%) | Machine higher, n (%) | Equal, n |
|---|---|---|---|---|---|---|---|---|---|
| Pre-hit | Frontal | -0.7256 | 0.475 | 0.570 | 0.560 | 0.720 | 12 (48.0) | 13 (52.0) | 0 |
| Pre-hit | Sensorimotor | -1.7712 | 0.089 | 0.268 | 0.252 | 0.554 | 15 (60.0) | 10 (40.0) | 0 |
| Pre-hit | Parietal | 1.1964 | 0.243 | 0.486 | 0.339 | 0.554 | 10 (40.0) | 15 (60.0) | 0 |
| Pre-hit | Posterior | 1.7775 | 0.088 | 0.268 | 0.120 | 0.360 | 11 (44.0) | 14 (56.0) | 0 |
| Pre-hit | Temporal | -0.7664 | 0.451 | 0.570 | 0.615 | 0.738 | 14 (56.0) | 11 (44.0) | 0 |
| Pre-hit | Other | 0.2855 | 0.778 | 0.823 | 0.853 | 0.903 | 12 (48.0) | 13 (52.0) | 0 |
| Impact | Frontal | 0.1899 | 0.851 | 0.851 | 0.958 | 0.958 | 13 (52.0) | 12 (48.0) | 0 |
| Impact | Sensorimotor | -1.2089 | 0.238 | 0.486 | 0.833 | 0.903 | 11 (44.0) | 14 (56.0) | 0 |
| Impact | Parietal | 2.3452 | 0.028 | 0.166 | 0.024 | 0.106 | 8 (32.0) | 17 (68.0) | 0 |
| Impact | Posterior | 0.9382 | 0.357 | 0.536 | 0.300 | 0.554 | 11 (44.0) | 14 (56.0) | 0 |
| Impact | Temporal | 1.3278 | 0.197 | 0.486 | 0.275 | 0.554 | 9 (36.0) | 16 (64.0) | 0 |
| Impact | Other | 2.0223 | 0.054 | 0.245 | 0.022 | 0.106 | 7 (28.0) | 18 (72.0) | 0 |
| Post-hit | Frontal | -0.9649 | 0.344 | 0.536 | 0.426 | 0.639 | 14 (56.0) | 11 (44.0) | 0 |
| Post-hit | Sensorimotor | -0.8689 | 0.393 | 0.545 | 0.560 | 0.720 | 11 (44.0) | 14 (56.0) | 0 |

| Window | Sensor region | Paired t | Paired t p | Paired t FDR q | Wilcoxon p | Wilcoxon FDR q | Machine lower, n (%) | Machine higher, n (%) | Equal, n |
|---|---|---|---|---|---|---|---|---|---|
| Post-hit | Parietal | 2.8213 | 0.009 | 0.166 | 0.004 | 0.075 | 4 (16.0) | 21 (84.0) | 0 |
| Post-hit | Posterior | 2.3828 | 0.025 | 0.166 | 0.014 | 0.106 | 7 (28.0) | 18 (72.0) | 0 |
| Post-hit | Temporal | 0.6351 | 0.531 | 0.598 | 0.325 | 0.554 | 10 (40.0) | 15 (60.0) | 0 |
| Post-hit | Other | 1.0703 | 0.295 | 0.531 | 0.080 | 0.289 | 7 (28.0) | 18 (72.0) | 0 |

*Note.* Positive machine − human values indicate larger values during ball-machine play. Benjamini–Hochberg correction was applied separately within each fixed 18-test feature family (aperiodic exponent and aperiodic offset) and separately for the two paired tests.

Supplementary Table S4. Spectral resolution, power-spectral-density settings, and spectral-model settings. Timing, Welch power-spectral-density settings, frequency resolution, and specparam settings for every model.

#### Model roles and temporal definitions

| Analysis | Model role | Window | Window start (s) | Window end (s) | Nominal duration (s) | Samples |
|---|---|---|---|---|---|---|
| Initial short-window specification | Initial specification / sensitivity comparator | Pre-hit | -1.0000 | -0.2000 | 0.8000 | 201 |
| Initial short-window specification | Initial specification / sensitivity comparator | Impact | -0.2000 | 0.3000 | 0.5000 | 126 |
| Initial short-window specification | Initial specification / sensitivity comparator | Post-hit | 0.3000 | 1.0000 | 0.7000 | 176 |
| Resolution-aware max6 | Principal resolution-aware short-window specification | Pre-hit | -1.0000 | -0.2000 | 0.8000 | 201 |
| Resolution-aware max6 | Principal resolution-aware short-window specification | Impact | -0.2000 | 0.3000 | 0.5000 | 126 |
| Resolution-aware max6 | Principal resolution-aware short-window specification | Post-hit | 0.3000 | 1.0000 | 0.7000 | 176 |
| Resolution-aware max3 | Reduced-complexity sensitivity analysis | Pre-hit | -1.0000 | -0.2000 | 0.8000 | 201 |
| Resolution-aware max3 | Reduced-complexity sensitivity analysis | Impact | -0.2000 | 0.3000 | 0.5000 | 126 |
| Resolution-aware max3 | Reduced-complexity sensitivity analysis | Post-hit | 0.3000 | 1.0000 | 0.7000 | 176 |
| Two-second max6 | Better-resolved peri-hit sensitivity analysis | Two-second peri-hit | -1.0000 | 1.0000 | 2.0000 | 501 |
| Two-second aperiodic-only | Aperiodic-only sensitivity analysis | Two-second peri-hit | -1.0000 | 1.0000 | 2.0000 | 501 |

#### Power-spectral-density and spectral-parameterization settings

| Analysis | Window | Welch window function | FFT length | Welch segment length | Welch overlap | Frequency resolution (Hz) | Fit minimum (Hz) | Fit maximum (Hz) | Minimum peak width (Hz) | Maximum peak width (Hz) | Maximum peaks | Peak threshold (SD) | Minimum peak height | Aperiodic mode | Spectral-model API |
|---|---|---|---|---|---|---|---|---|---|---|---|---|---|---|---|
| Initial short-window specification | Pre-hit | Hamming | 201 | 201 | 100 | 1.2438 | 3.0000 | 40.0000 | 1.000 | 8.000 | 6 | 2.0000 | 0.050 | fixed | specparam |
| Initial short-window specification | Impact | Hamming | 126 | 126 | 63 | 1.9841 | 3.0000 | 40.0000 | 1.000 | 8.000 | 6 | 2.0000 | 0.050 | fixed | specparam |
| Initial short-window specification | Post-hit | Hamming | 176 | 176 | 88 | 1.4205 | 3.0000 | 40.0000 | 1.000 | 8.000 | 6 | 2.0000 | 0.050 | fixed | specparam |
| Resolution-aware max6 | Pre-hit | Hamming | 201 | 201 | 100 | 1.2438 | 3.0000 | 40.0000 | 2.488 | 8.000 | 6 | 2.0000 | 0.050 | fixed | specparam |
| Resolution-aware max6 | Impact | Hamming | 126 | 126 | 63 | 1.9841 | 3.0000 | 40.0000 | 3.968 | 8.000 | 6 | 2.0000 | 0.050 | fixed | specparam |
| Resolution-aware max6 | Post-hit | Hamming | 176 | 176 | 88 | 1.4205 | 3.0000 | 40.0000 | 2.841 | 8.000 | 6 | 2.0000 | 0.050 | fixed | specparam |
| Resolution-aware max3 | Pre-hit | Hamming | 201 | 201 | 100 | 1.2438 | 3.0000 | 40.0000 | 2.488 | 8.000 | 3 | 2.0000 | 0.050 | fixed | specparam |
| Resolution-aware max3 | Impact | Hamming | 126 | 126 | 63 | 1.9841 | 3.0000 | 40.0000 | 3.968 | 8.000 | 3 | 2.0000 | 0.050 | fixed | specparam |
| Resolution-aware max3 | Post-hit | Hamming | 176 | 176 | 88 | 1.4205 | 3.0000 | 40.0000 | 2.841 | 8.000 | 3 | 2.0000 | 0.050 | fixed | specparam |
| Two-second max6 | Two-second peri-hit | Hamming | 501 | 501 | 0 | 0.4990 | 3.0000 | 40.0000 | 0.998 | 8.000 | 6 | 2.0000 | 0.050 | fixed | specparam |
| Two-second aperiodic-only | Two-second peri-hit | Hamming | 501 | 501 | 0 | 0.4990 | 3.0000 | 40.0000 | 0.998 | 8.000 | 0 | 2.0000 | 0.050 | fixed | specparam |

*Note.* The short-window models use window-specific FFT lengths and frequency resolutions. Minimum peak widths in the resolution-aware models were constrained by the available frequency grid. The aperiodic-only model sets maximum peaks to zero by design. PSD = power spectral density; FFT = fast Fourier transform.

Supplementary Table S5. Fit-quality and peak-detection summaries across spectral models. Descriptive model-fit and periodic-peak-detection summaries across short-window and two-second spectral specifications. Each section is presented using only the fields relevant to that output.

### Section 1. Short-window fit quality

| Grouping | Model cells | Mean $R^2$ | Median $R^2$ | $R^2$ Q1 | Minimum $R^2$ | Mean error | Median error | Error Q3 | Maximum error | $R^2 < 0.8$, n | Fraction $R^2 < 0.8$ | Context | Analysis | Window | Sensor region |
|---|---|---|---|---|---|---|---|---|---|---|---|---|---|---|---|
| Overall | 900 | 0.8708 | 0.9527 | 0.8693 | 0.0000 | 0.0386 | 0.0326 | 0.0487 | 0.1335 | 171 | 0.1900 | — | Initial short-window specification | — | — |
| Window | 300 | 0.8729 | 0.9569 | 0.8647 | 0.0000 | 0.0395 | 0.0335 | 0.0489 | 0.1330 | 58 | 0.1933 | — | Initial short-window specification | Impact | — |
| Window | 300 | 0.8707 | 0.9507 | 0.8712 | 0.0124 | 0.0375 | 0.0311 | 0.0467 | 0.1240 | 55 | 0.1833 | — | Initial short-window specification | Post-hit | — |
| Window | 300 | 0.8689 | 0.9502 | 0.8807 | 0.0004 | 0.0390 | 0.0336 | 0.0504 | 0.1335 | 58 | 0.1933 | — | Initial short-window specification | Pre-hit | — |
| Window × context | 150 | 0.8630 | 0.9452 | 0.8523 | 0.0027 | 0.0404 | 0.0336 | 0.0494 | 0.1274 | 31 | 0.2067 | Human opponent | Initial short-window specification | Impact | — |
| Window × context | 150 | 0.8624 | 0.9431 | 0.8505 | 0.0124 | 0.0376 | 0.0313 | 0.0447 | 0.1240 | 29 | 0.1933 | Human opponent | Initial short-window specification | Post-hit | — |
| Window × context | 150 | 0.8566 | 0.9364 | 0.8466 | 0.0004 | 0.0403 | 0.0351 | 0.0509 | 0.1229 | 30 | 0.2000 | Human opponent | Initial short-window specification | Pre-hit | — |
| Window × context | 150 | 0.8827 | 0.9635 | 0.8861 | 0.0000 | 0.0386 | 0.0327 | 0.0477 | 0.1330 | 27 | 0.1800 | Ball machine | Initial short-window specification | Impact | — |
| Window × context | 150 | 0.8790 | 0.9557 | 0.8766 | 0.0404 | 0.0374 | 0.0310 | 0.0477 | 0.1172 | 26 | 0.1733 | Ball machine | Initial short-window specification | Post-hit | — |
| Window × context | 150 | 0.8811 | 0.9601 | 0.9010 | 0.0008 | 0.0376 | 0.0329 | 0.0490 | 0.1335 | 28 | 0.1867 | Ball machine | Initial short-window specification | Pre-hit | — |
| Window × region | 50 | 0.9205 | 0.9609 | 0.9140 | 0.5759 | 0.0634 | 0.0634 | 0.0901 | 0.1330 | 6 | 0.1200 | — | Initial short-window specification | Impact | Frontal |
| Window × region | 50 | 0.9460 | 0.9606 | 0.9348 | 0.7174 | 0.0325 | 0.0295 | 0.0427 | 0.0665 | 1 | 0.0200 | — | Initial short-window specification | Impact | Other |
| Window × region | 50 | 0.8748 | 0.9713 | 0.8792 | 0.1569 | 0.0337 | 0.0304 | 0.0434 | 0.0801 | 10 | 0.2000 | — | Initial short-window specification | Impact | Parietal |
| Window × region | 50 | 0.9027 | 0.9510 | 0.8599 | 0.5129 | 0.0258 | 0.0239 | 0.0306 | 0.0597 | 9 | 0.1800 | — | Initial short-window specification | Impact | Posterior |
| Window × region | 50 | 0.9266 | 0.9756 | 0.9361 | 0.4789 | 0.0359 | 0.0315 | 0.0470 | 0.0939 | 7 | 0.1400 | — | Initial short-window specification | Impact | Sensorimotor |
| Window × region | 50 | 0.6666 | 0.7795 | 0.4327 | 0.0000 | 0.0456 | 0.0421 | 0.0577 | 0.1213 | 25 | 0.5000 | — | Initial short-window specification | Impact | Temporal |
| Window × region | 50 | 0.9145 | 0.9597 | 0.9048 | 0.5096 | 0.0602 | 0.0586 | 0.0850 | 0.1240 | 5 | 0.1000 | — | Initial short-window specification | Post-hit | Frontal |
| Window × region | 50 | 0.9466 | 0.9613 | 0.9383 | 0.7557 | 0.0289 | 0.0254 | 0.0367 | 0.0588 | 2 | 0.0400 | — | Initial short-window specification | Post-hit | Other |
| Window × region | 50 | 0.8883 | 0.9587 | 0.9051 | 0.0886 | 0.0311 | 0.0275 | 0.0402 | 0.0807 | 9 | 0.1800 | — | Initial short-window specification | Post-hit | Parietal |
| Window × region | 50 | 0.9064 | 0.9423 | 0.8611 | 0.5275 | 0.0264 | 0.0251 | 0.0325 | 0.0491 | 6 | 0.1200 | — | Initial short-window specification | Post-hit | Posterior |
| Window × region | 50 | 0.9182 | 0.9681 | 0.9108 | 0.4500 | 0.0348 | 0.0301 | 0.0451 | 0.0876 | 6 | 0.1200 | — | Initial short-window specification | Post-hit | Sensorimotor |
| Window × region | 50 | 0.6502 | 0.7110 | 0.4257 | 0.0124 | 0.0436 | 0.0371 | 0.0500 | 0.1172 | 27 | 0.5400 | — | Initial short-window specification | Post-hit | Temporal |
| Window × region | 50 | 0.9232 | 0.9628 | 0.9365 | 0.4128 | 0.0591 | 0.0557 | 0.0759 | 0.1335 | 5 | 0.1000 | — | Initial short-window specification | Pre-hit | Frontal |
| Window × region | 50 | 0.9396 | 0.9546 | 0.9306 | 0.6795 | 0.0321 | 0.0287 | 0.0401 | 0.0723 | 2 | 0.0400 | — | Initial short-window specification | Pre-hit | Other |
| Window × region | 50 | 0.8825 | 0.9636 | 0.8892 | 0.1830 | 0.0341 | 0.0302 | 0.0430 | 0.0898 | 8 | 0.1600 | — | Initial short-window specification | Pre-hit | Parietal |
| Window × region | 50 | 0.8885 | 0.9373 | 0.8067 | 0.4162 | 0.0284 | 0.0260 | 0.0357 | 0.0580 | 12 | 0.2400 | — | Initial short-window specification | Pre-hit | Posterior |
| Window × region | 50 | 0.9314 | 0.9730 | 0.9285 | 0.3508 | 0.0337 | 0.0276 | 0.0475 | 0.0634 | 4 | 0.0800 | — | Initial short-window specification | Pre-hit | Sensorimotor |
| Window × region | 50 | 0.6479 | 0.7504 | 0.3770 | 0.0004 | 0.0463 | 0.0429 | 0.0587 | 0.0858 | 27 | 0.5400 | — | Initial short-window specification | Pre-hit | Temporal |
| Overall | 900 | 0.8686 | 0.9504 | 0.8601 | 0.0000 | 0.0391 | 0.0332 | 0.0491 | 0.1335 | 175 | 0.1944 | — | Resolution-aware max6 | — | — |
| Window | 300 | 0.8661 | 0.9535 | 0.8523 | 0.0000 | 0.0407 | 0.0345 | 0.0514 | 0.1330 | 63 | 0.2100 | — | Resolution-aware max6 | Impact | — |
| Window | 300 | 0.8689 | 0.9466 | 0.8575 | 0.0124 | 0.0384 | 0.0319 | 0.0460 | 0.1240 | 54 | 0.1800 | — | Resolution-aware max6 | Post-hit | — |
| Window | 300 | 0.8707 | 0.9511 | 0.8832 | 0.0004 | 0.0383 | 0.0322 | 0.0494 | 0.1335 | 58 | 0.1933 | — | Resolution-aware max6 | Pre-hit | — |
| Window × context | 150 | 0.8560 | 0.9420 | 0.8280 | 0.0027 | 0.0419 | 0.0344 | 0.0547 | 0.1274 | 33 | 0.2200 | Human opponent | Resolution-aware max6 | Impact | — |
| Window × context | 150 | 0.8607 | 0.9367 | 0.8460 | 0.0124 | 0.0389 | 0.0329 | 0.0449 | 0.1240 | 29 | 0.1933 | Human opponent | Resolution-aware max6 | Post-hit | — |
| Window × context | 150 | 0.8601 | 0.9481 | 0.8442 | 0.0004 | 0.0397 | 0.0335 | 0.0506 | 0.1229 | 30 | 0.2000 | Human opponent | Resolution-aware max6 | Pre-hit | — |
| Window × context | 150 | 0.8763 | 0.9613 | 0.8842 | 0.0000 | 0.0394 | 0.0345 | 0.0494 | 0.1330 | 30 | 0.2000 | Ball machine | Resolution-aware max6 | Impact | — |
| Window × context | 150 | 0.8771 | 0.9526 | 0.8744 | 0.0257 | 0.0379 | 0.0317 | 0.0478 | 0.1172 | 25 | 0.1667 | Ball machine | Resolution-aware max6 | Post-hit | — |
| Window × context | 150 | 0.8813 | 0.9605 | 0.8942 | 0.0008 | 0.0369 | 0.0307 | 0.0483 | 0.1335 | 28 | 0.1867 | Ball machine | Resolution-aware max6 | Pre-hit | — |
| Window × region | 50 | 0.9181 | 0.9556 | 0.9251 | 0.5759 | 0.0625 | 0.0607 | 0.0834 | 0.1330 | 7 | 0.1400 | — | Resolution-aware max6 | Impact | Frontal |
| Window × region | 50 | 0.9416 | 0.9593 | 0.9336 | 0.7174 | 0.0332 | 0.0292 | 0.0427 | 0.0728 | 3 | 0.0600 | — | Resolution-aware max6 | Impact | Other |
| Window × region | 50 | 0.8721 | 0.9715 | 0.8792 | 0.1569 | 0.0341 | 0.0305 | 0.0440 | 0.0801 | 10 | 0.2000 | — | Resolution-aware max6 | Impact | Parietal |
| Window × region | 50 | 0.8977 | 0.9489 | 0.8438 | 0.5127 | 0.0277 | 0.0262 | 0.0339 | 0.0604 | 10 | 0.2000 | — | Resolution-aware max6 | Impact | Posterior |
| Window × region | 50 | 0.9210 | 0.9704 | 0.9391 | 0.4016 | 0.0389 | 0.0346 | 0.0491 | 0.0939 | 6 | 0.1200 | — | Resolution-aware max6 | Impact | Sensorimotor |
| Window × region | 50 | 0.6461 | 0.7217 | 0.3815 | 0.0000 | 0.0476 | 0.0450 | 0.0577 | 0.1213 | 27 | 0.5400 | — | Resolution-aware max6 | Impact | Temporal |
| Window × region | 50 | 0.9116 | 0.9511 | 0.8855 | 0.5096 | 0.0610 | 0.0589 | 0.0850 | 0.1240 | 5 | 0.1000 | — | Resolution-aware max6 | Post-hit | Frontal |
| Window × region | 50 | 0.9454 | 0.9608 | 0.9383 | 0.7558 | 0.0298 | 0.0256 | 0.0384 | 0.0589 | 2 | 0.0400 | — | Resolution-aware max6 | Post-hit | Other |
| Window × region | 50 | 0.8936 | 0.9585 | 0.8997 | 0.0886 | 0.0309 | 0.0280 | 0.0390 | 0.0807 | 8 | 0.1600 | — | Resolution-aware max6 | Post-hit | Parietal |
| Window × region | 50 | 0.9023 | 0.9378 | 0.8575 | 0.5276 | 0.0277 | 0.0261 | 0.0351 | 0.0491 | 7 | 0.1400 | — | Resolution-aware max6 | Post-hit | Posterior |
| Window × region | 50 | 0.9167 | 0.9655 | 0.8930 | 0.4357 | 0.0363 | 0.0325 | 0.0451 | 0.0876 | 5 | 0.1000 | — | Resolution-aware max6 | Post-hit | Sensorimotor |
| Window × region | 50 | 0.6438 | 0.7293 | 0.4257 | 0.0124 | 0.0447 | 0.0402 | 0.0493 | 0.1172 | 27 | 0.5400 | — | Resolution-aware max6 | Post-hit | Temporal |
| Window × region | 50 | 0.9270 | 0.9642 | 0.9395 | 0.4128 | 0.0556 | 0.0520 | 0.0724 | 0.1335 | 5 | 0.1000 | — | Resolution-aware max6 | Pre-hit | Frontal |
| Window × region | 50 | 0.9396 | 0.9572 | 0.9306 | 0.6777 | 0.0325 | 0.0298 | 0.0389 | 0.0723 | 2 | 0.0400 | — | Resolution-aware max6 | Pre-hit | Other |
| Window × region | 50 | 0.8834 | 0.9626 | 0.8921 | 0.1830 | 0.0336 | 0.0287 | 0.0428 | 0.0898 | 8 | 0.1600 | — | Resolution-aware max6 | Pre-hit | Parietal |
| Window × region | 50 | 0.8871 | 0.9372 | 0.8067 | 0.4168 | 0.0290 | 0.0270 | 0.0357 | 0.0581 | 12 | 0.2400 | — | Resolution-aware max6 | Pre-hit | Posterior |
| Window × region | 50 | 0.9348 | 0.9832 | 0.9445 | 0.3508 | 0.0317 | 0.0245 | 0.0449 | 0.0629 | 4 | 0.0800 | — | Resolution-aware max6 | Pre-hit | Sensorimotor |
| Window × region | 50 | 0.6525 | 0.7505 | 0.4235 | 0.0004 | 0.0475 | 0.0442 | 0.0587 | 0.0859 | 27 | 0.5400 | — | Resolution-aware max6 | Pre-hit | Temporal |
| Overall | 900 | 0.8634 | 0.9441 | 0.8527 | 0.0000 | 0.0419 | 0.0368 | 0.0523 | 0.1335 | 180 | 0.2000 | — | Resolution-aware max3 | — | — |
| Window | 300 | 0.8644 | 0.9516 | 0.8523 | 0.0000 | 0.0426 | 0.0368 | 0.0543 | 0.1330 | 63 | 0.2100 | — | Resolution-aware max3 | Impact | — |
| Window | 300 | 0.8621 | 0.9411 | 0.8485 | 0.0124 | 0.0409 | 0.0350 | 0.0498 | 0.1240 | 59 | 0.1967 | — | Resolution-aware max3 | Post-hit | — |
| Window | 300 | 0.8636 | 0.9400 | 0.8660 | 0.0004 | 0.0423 | 0.0381 | 0.0528 | 0.1335 | 58 | 0.1933 | — | Resolution-aware max3 | Pre-hit | — |
| Window × context | 150 | 0.8542 | 0.9407 | 0.8280 | 0.0027 | 0.0436 | 0.0385 | 0.0563 | 0.1274 | 33 | 0.2200 | Human opponent | Resolution-aware max3 | Impact | — |
| Window × context | 150 | 0.8526 | 0.9335 | 0.8316 | 0.0124 | 0.0411 | 0.0347 | 0.0512 | 0.1240 | 32 | 0.2133 | Human opponent | Resolution-aware max3 | Post-hit | — |
| Window × context | 150 | 0.8528 | 0.9306 | 0.8429 | 0.0004 | 0.0431 | 0.0405 | 0.0534 | 0.1229 | 30 | 0.2000 | Human opponent | Resolution-aware max3 | Pre-hit | — |
| Window × context | 150 | 0.8746 | 0.9571 | 0.8842 | 0.0000 | 0.0415 | 0.0361 | 0.0519 | 0.1330 | 30 | 0.2000 | Ball machine | Resolution-aware max3 | Impact | — |
| Window × context | 150 | 0.8715 | 0.9464 | 0.8726 | 0.0257 | 0.0407 | 0.0357 | 0.0494 | 0.1172 | 27 | 0.1800 | Ball machine | Resolution-aware max3 | Post-hit | — |
| Window × context | 150 | 0.8744 | 0.9464 | 0.8875 | 0.0008 | 0.0414 | 0.0360 | 0.0523 | 0.1335 | 28 | 0.1867 | Ball machine | Resolution-aware max3 | Pre-hit | — |
| Window × region | 50 | 0.9151 | 0.9556 | 0.9251 | 0.5759 | 0.0682 | 0.0629 | 0.0834 | 0.1330 | 7 | 0.1400 | — | Resolution-aware max3 | Impact | Frontal |
| Window × region | 50 | 0.9412 | 0.9593 | 0.9336 | 0.7174 | 0.0339 | 0.0313 | 0.0429 | 0.0728 | 3 | 0.0600 | — | Resolution-aware max3 | Impact | Other |
| Window × region | 50 | 0.8702 | 0.9708 | 0.8792 | 0.1569 | 0.0354 | 0.0332 | 0.0446 | 0.0801 | 10 | 0.2000 | — | Resolution-aware max3 | Impact | Parietal |
| Window × region | 50 | 0.8955 | 0.9388 | 0.8438 | 0.5127 | 0.0295 | 0.0277 | 0.0357 | 0.0682 | 10 | 0.2000 | — | Resolution-aware max3 | Impact | Posterior |
| Window × region | 50 | 0.9195 | 0.9680 | 0.9361 | 0.4016 | 0.0401 | 0.0407 | 0.0516 | 0.0939 | 6 | 0.1200 | — | Resolution-aware max3 | Impact | Sensorimotor |
| Window × region | 50 | 0.6450 | 0.7217 | 0.3815 | 0.0000 | 0.0483 | 0.0450 | 0.0577 | 0.1213 | 27 | 0.5400 | — | Resolution-aware max3 | Impact | Temporal |
| Window × region | 50 | 0.9066 | 0.9488 | 0.8855 | 0.5096 | 0.0665 | 0.0663 | 0.0876 | 0.1240 | 5 | 0.1000 | — | Resolution-aware max3 | Post-hit | Frontal |
| Window × region | 50 | 0.9419 | 0.9585 | 0.9376 | 0.7558 | 0.0316 | 0.0293 | 0.0397 | 0.0589 | 2 | 0.0400 | — | Resolution-aware max3 | Post-hit | Other |
| Window × region | 50 | 0.8908 | 0.9503 | 0.8963 | 0.0886 | 0.0325 | 0.0295 | 0.0402 | 0.0807 | 9 | 0.1800 | — | Resolution-aware max3 | Post-hit | Parietal |
| Window × region | 50 | 0.8962 | 0.9333 | 0.8575 | 0.5276 | 0.0298 | 0.0266 | 0.0380 | 0.0679 | 8 | 0.1600 | — | Resolution-aware max3 | Post-hit | Posterior |

| Grouping | Model cells | Mean $R^2$ | Median $R^2$ | $R^2$ Q1 | Minimum $R^2$ | Mean error | Median error | Error Q3 | Maximum error | $R^2 < 0.8$, n | Fraction $R^2 < 0.8$ | Context | Analysis | Window | Sensor region |
|---|---|---|---|---|---|---|---|---|---|---|---|---|---|---|---|
| Window × region | 50 | 0.9104 | 0.9648 | 0.8907 | 0.4357 | 0.0385 | 0.0333 | 0.0544 | 0.0876 | 6 | 0.1200 | — | Resolution-aware max3 | Post-hit | Sensorimotor |
| Window × region | 50 | 0.6266 | 0.7003 | 0.4215 | 0.0124 | 0.0466 | 0.0438 | 0.0576 | 0.1172 | 29 | 0.5800 | — | Resolution-aware max3 | Post-hit | Temporal |
| Window × region | 50 | 0.9170 | 0.9524 | 0.9268 | 0.4128 | 0.0644 | 0.0638 | 0.0785 | 0.1335 | 5 | 0.1000 | — | Resolution-aware max3 | Pre-hit | Frontal |
| Window × region | 50 | 0.9324 | 0.9464 | 0.9261 | 0.6777 | 0.0360 | 0.0325 | 0.0484 | 0.0723 | 2 | 0.0400 | — | Resolution-aware max3 | Pre-hit | Other |
| Window × region | 50 | 0.8767 | 0.9492 | 0.8892 | 0.1830 | 0.0368 | 0.0340 | 0.0480 | 0.0898 | 8 | 0.1600 | — | Resolution-aware max3 | Pre-hit | Parietal |
| Window × region | 50 | 0.8843 | 0.9254 | 0.8067 | 0.4168 | 0.0304 | 0.0283 | 0.0369 | 0.0581 | 12 | 0.2400 | — | Resolution-aware max3 | Pre-hit | Posterior |
| Window × region | 50 | 0.9260 | 0.9726 | 0.9284 | 0.3508 | 0.0364 | 0.0351 | 0.0482 | 0.0637 | 4 | 0.0800 | — | Resolution-aware max3 | Pre-hit | Sensorimotor |
| Window × region | 50 | 0.6452 | 0.7505 | 0.4235 | 0.0004 | 0.0496 | 0.0478 | 0.0587 | 0.0915 | 27 | 0.5400 | — | Resolution-aware max3 | Pre-hit | Temporal |

### Section 2. Short-window peak detection

| Analysis | Window | Model cells | Peaks present | Detection rate | Band |
|---|---|---|---|---|---|
| Initial short-window specification | Pre-hit | 300 | 17 | 0.0567 | Theta |
| Initial short-window specification | Pre-hit | 300 | 115 | 0.3833 | Alpha |
| Initial short-window specification | Pre-hit | 300 | 77 | 0.2567 | Beta |
| Initial short-window specification | Impact | 300 | 20 | 0.0667 | Theta |
| Initial short-window specification | Impact | 300 | 104 | 0.3467 | Alpha |
| Initial short-window specification | Impact | 300 | 51 | 0.1700 | Beta |
| Initial short-window specification | Post-hit | 300 | 18 | 0.0600 | Theta |
| Initial short-window specification | Post-hit | 300 | 121 | 0.4033 | Alpha |
| Initial short-window specification | Post-hit | 300 | 77 | 0.2567 | Beta |
| Resolution-aware max6 | Pre-hit | 300 | 21 | 0.0700 | Theta |
| Resolution-aware max6 | Pre-hit | 300 | 114 | 0.3800 | Alpha |
| Resolution-aware max6 | Pre-hit | 300 | 84 | 0.2800 | Beta |
| Resolution-aware max6 | Impact | 300 | 23 | 0.0767 | Theta |
| Resolution-aware max6 | Impact | 300 | 88 | 0.2933 | Alpha |
| Resolution-aware max6 | Impact | 300 | 47 | 0.1567 | Beta |
| Resolution-aware max6 | Post-hit | 300 | 23 | 0.0767 | Theta |
| Resolution-aware max6 | Post-hit | 300 | 116 | 0.3867 | Alpha |
| Resolution-aware max6 | Post-hit | 300 | 74 | 0.2467 | Beta |
| Resolution-aware max3 | Pre-hit | 300 | 15 | 0.0500 | Theta |
| Resolution-aware max3 | Pre-hit | 300 | 105 | 0.3500 | Alpha |
| Resolution-aware max3 | Pre-hit | 300 | 47 | 0.1567 | Beta |
| Resolution-aware max3 | Impact | 300 | 16 | 0.0533 | Theta |
| Resolution-aware max3 | Impact | 300 | 84 | 0.2800 | Alpha |
| Resolution-aware max3 | Impact | 300 | 41 | 0.1367 | Beta |
| Resolution-aware max3 | Post-hit | 300 | 16 | 0.0533 | Theta |
| Resolution-aware max3 | Post-hit | 300 | 108 | 0.3600 | Alpha |
| Resolution-aware max3 | Post-hit | 300 | 54 | 0.1800 | Beta |

### Section 3. Two-second max6 peak detection

| Context | Analysis | Window | Model cells | Peaks present | Detection rate | Band |
|---|---|---|---|---|---|---|
| All contexts | Two-second max6 | Two-second peri-hit | 300 | 33 | 0.1100 | Theta |
| Human opponent | Two-second max6 | Two-second peri-hit | 150 | 13 | 0.0867 | Theta |
| Ball machine | Two-second max6 | Two-second peri-hit | 150 | 20 | 0.1333 | Theta |
| All contexts | Two-second max6 | Two-second peri-hit | 300 | 145 | 0.4833 | Alpha |
| Human opponent | Two-second max6 | Two-second peri-hit | 150 | 73 | 0.4867 | Alpha |
| Ball machine | Two-second max6 | Two-second peri-hit | 150 | 72 | 0.4800 | Alpha |
| All contexts | Two-second max6 | Two-second peri-hit | 300 | 150 | 0.5000 | Beta |
| Human opponent | Two-second max6 | Two-second peri-hit | 150 | 72 | 0.4800 | Beta |
| Ball machine | Two-second max6 | Two-second peri-hit | 150 | 78 | 0.5200 | Beta |

### Section 4. Aperiodic-only peak validation

| Analysis | Window | Model cells | Peaks present | Band | All peak parameters missing |
|---|---|---|---|---|---|
| Two-second aperiodic-only | Two-second peri-hit | 300 | 0 | Theta | Yes |
| Two-second aperiodic-only | Two-second peri-hit | 300 | 0 | Alpha | Yes |
| Two-second aperiodic-only | Two-second peri-hit | 300 | 0 | Beta | Yes |

### Section 5. Aperiodic-only fit-quality change

| Grouping | Model cells | Context | Sensor region | Analysis | Mean $\Delta R^2$ (aperiodic-only − max6) | Median $\Delta R^2$ (aperiodic-only − max6) | Fraction aperiodic-only higher $R^2$ | Mean $\Delta$error (aperiodic-only − max6) | Median $\Delta$error (aperiodic-only − max6) | Fraction aperiodic-only lower error |
|---|---|---|---|---|---|---|---|---|---|---|
| Overall | 300 | — | — | Two-second aperiodic-only minus two-second max6 | -0.1168 | -0.0667 | 0.0133 | 0.0278 | 0.0257 | 0.0100 |
| Region | 50 | — | Frontal | Two-second aperiodic-only minus two-second max6 | -0.0579 | -0.0531 | 0.0000 | 0.0378 | 0.0400 | 0.0000 |
| Region | 50 | — | Other | Two-second aperiodic-only minus two-second max6 | -0.1489 | -0.0830 | 0.0000 | 0.0298 | 0.0264 | 0.0000 |
| Region | 50 | — | Parietal | Two-second aperiodic-only minus two-second max6 | -0.0946 | -0.0602 | 0.0400 | 0.0246 | 0.0198 | 0.0200 |
| Region | 50 | — | Posterior | Two-second aperiodic-only minus two-second max6 | -0.1697 | -0.0952 | 0.0200 | 0.0255 | 0.0214 | 0.0200 |
| Region | 50 | — | Sensorimotor | Two-second aperiodic-only minus two-second max6 | -0.0680 | -0.0599 | 0.0000 | 0.0280 | 0.0260 | 0.0200 |
| Region | 50 | — | Temporal | Two-second aperiodic-only minus two-second max6 | -0.1615 | -0.1122 | 0.0200 | 0.0212 | 0.0162 | 0.0000 |
| Region × context | 25 | Human opponent | Frontal | Two-second aperiodic-only minus two-second max6 | -0.0600 | -0.0626 | 0.0000 | 0.0369 | 0.0405 | 0.0000 |
| Region × context | 25 | Human opponent | Other | Two-second aperiodic-only minus two-second max6 | -0.1821 | -0.1127 | 0.0000 | 0.0314 | 0.0318 | 0.0000 |
| Region × context | 25 | Human opponent | Parietal | Two-second aperiodic-only minus two-second max6 | -0.1019 | -0.0618 | 0.0800 | 0.0225 | 0.0169 | 0.0400 |
| Region × context | 25 | Human opponent | Posterior | Two-second aperiodic-only minus two-second max6 | -0.2035 | -0.1232 | 0.0400 | 0.0253 | 0.0214 | 0.0400 |
| Region × context | 25 | Human opponent | Sensorimotor | Two-second aperiodic-only minus two-second max6 | -0.0805 | -0.0789 | 0.0000 | 0.0272 | 0.0258 | 0.0000 |
| Region × context | 25 | Human opponent | Temporal | Two-second aperiodic-only minus two-second max6 | -0.1891 | -0.1199 | 0.0400 | 0.0207 | 0.0150 | 0.0000 |
| Region × context | 25 | Ball machine | Frontal | Two-second aperiodic-only minus two-second max6 | -0.0558 | -0.0502 | 0.0000 | 0.0387 | 0.0396 | 0.0000 |
| Region × context | 25 | Ball machine | Other | Two-second aperiodic-only minus two-second max6 | -0.1157 | -0.0665 | 0.0000 | 0.0283 | 0.0257 | 0.0000 |
| Region × context | 25 | Ball machine | Parietal | Two-second aperiodic-only minus two-second max6 | -0.0874 | -0.0523 | 0.0000 | 0.0267 | 0.0239 | 0.0000 |
| Region × context | 25 | Ball machine | Posterior | Two-second aperiodic-only minus two-second max6 | -0.1358 | -0.0937 | 0.0000 | 0.0256 | 0.0214 | 0.0000 |
| Region × context | 25 | Ball machine | Sensorimotor | Two-second aperiodic-only minus two-second max6 | -0.0555 | -0.0580 | 0.0000 | 0.0288 | 0.0262 | 0.0400 |
| Region × context | 25 | Ball machine | Temporal | Two-second aperiodic-only minus two-second max6 | -0.1340 | -0.0985 | 0.0000 | 0.0218 | 0.0175 | 0.0000 |

*Note.* $R^2$ and error summarize spectral-model fit quality. Peak-detection rates are proportions of model cells in which a peak was present. Fit-quality and peak-detection summaries are descriptive diagnostics; model cells are not treated as independent inferential observations. In the aperiodic-only validation section, missing periodic-peak parameters are expected because periodic components were disabled by design.

Supplementary Table S6. Full robustness results: bootstrap, leave-one-participant-out, concordance, and BIC summaries. Complete robustness outputs in sectioned form. Each section is presented using only its populated fields so that unrelated metrics are not forced into a single wide schema.

### Section 1. Short-window focal-model comparison

| Model | Window | Sensor region | Feature | Participants | Human mean | Machine mean | Human median | Machine median | Machine − human mean | Machine − human median | Difference SD | Cohen's dz | Paired t | Paired t p | Wilcoxon p | Machine lower, n | Machine higher, n | Equal, n | Machine lower (%) | Machine higher (%) | Paired t FDR q | Wilcoxon FDR q |
|---|---|---|---|---|---|---|---|---|---|---|---|---|---|---|---|---|---|---|---|---|---|---|
| Initial short-window specification | Impact | Parietal | Aperiodic exponent | 25 | 0.6526 | 0.7841 | 0.6266 | 0.7533 | 0.1315 | 0.0979 | 0.1794 | 0.7331 | 3.6655 | 0.001 | <0.001 | 5 | 20 | 0 | 20.0000 | 80.0000 | 0.003 | 0.002 |
| Resolution-aware max6 | Impact | Parietal | Aperiodic exponent | 25 | 0.6416 | 0.7825 | 0.6148 | 0.7533 | 0.1409 | 0.0989 | 0.1836 | 0.7677 | 3.8384 | <0.001 | <0.001 | 5 | 20 | 0 | 20.0000 | 80.0000 | 0.002 | 0.002 |
| Resolution-aware max3 | Impact | Parietal | Aperiodic exponent | 25 | 0.6498 | 0.7774 | 0.6148 | 0.7533 | 0.1276 | 0.0989 | 0.1731 | 0.7369 | 3.6847 | 0.001 | <0.001 | 5 | 20 | 0 | 20.0000 | 80.0000 | 0.003 | 0.002 |

### Section 2. Short-window bootstrap

| Model | Participants | Bootstrap resamples | Random seed | Observed mean difference | Mean difference 95% CI, lower | Mean difference 95% CI, upper | Observed Cohen's dz | Cohen's dz 95% CI, lower | Cohen's dz 95% CI, upper | Bootstrap fraction mean > 0 |
|---|---|---|---|---|---|---|---|---|---|---|
| Initial short-window specification | 25 | 50000 | 20262594 | 0.1315 | 0.0650 | 0.2026 | 0.7331 | 0.4100 | 1.1842 | 1.0000 |
| Resolution-aware max6 | 25 | 50000 | 20262928 | 0.1409 | 0.0722 | 0.2132 | 0.7677 | 0.4367 | 1.2211 | 1.0000 |
| Resolution-aware max3 | 25 | 50000 | 20262925 | 0.1276 | 0.0635 | 0.1969 | 0.7369 | 0.4085 | 1.2074 | 1.0000 |

### Section 3. Short-window leave-one-participant-out

| Model | Excluded participant | Window | Sensor region | Feature | Participants | Human mean | Machine mean | Human median | Machine median | Machine − human mean | Machine − human median | Difference SD | Cohen's dz | Paired t | Paired t p | Wilcoxon p | Machine lower, n | Machine higher, n | Equal, n | Machine lower (%) | Machine higher (%) | Paired t FDR q | Wilcoxon FDR q |
|---|---|---|---|---|---|---|---|---|---|---|---|---|---|---|---|---|---|---|---|---|---|---|---|
| Initial short-window specification | Participant 01 | Impact | Parietal | Aperiodic exponent | 24 | 0.6579 | 0.7822 | 0.6307 | 0.7470 | 0.1243 | 0.0945 | 0.1795 | 0.6925 | 3.3925 | 0.003 | 0.001 | 5 | 19 | 0 | 20.8333 | 79.1667 | 0.006 | 0.004 |
| Initial short-window specification | Participant 02 | Impact | Parietal | Aperiodic exponent | 24 | 0.6555 | 0.7783 | 0.6307 | 0.7470 | 0.1228 | 0.0945 | 0.1778 | 0.6908 | 3.3843 | 0.003 | 0.001 | 5 | 19 | 0 | 20.8333 | 79.1667 | 0.007 | 0.004 |
| Initial short-window specification | Participant 03 | Impact | Parietal | Aperiodic exponent | 24 | 0.6643 | 0.7982 | 0.6307 | 0.7692 | 0.1339 | 0.1051 | 0.1829 | 0.7322 | 3.5869 | 0.002 | <0.001 | 5 | 19 | 0 | 20.8333 | 79.1667 | 0.004 | 0.002 |
| Initial short-window specification | Participant 04 | Impact | Parietal | Aperiodic exponent | 24 | 0.6442 | 0.7739 | 0.6090 | 0.7470 | 0.1297 | 0.0945 | 0.1830 | 0.7087 | 3.4718 | 0.002 | 0.001 | 5 | 19 | 0 | 20.8333 | 79.1667 | 0.005 | 0.003 |
| Initial short-window specification | Participant 05 | Impact | Parietal | Aperiodic exponent | 24 | 0.6439 | 0.7628 | 0.6090 | 0.7470 | 0.1188 | 0.0945 | 0.1714 | 0.6932 | 3.3962 | 0.002 | 0.001 | 5 | 19 | 0 | 20.8333 | 79.1667 | 0.006 | 0.004 |
| Initial short-window specification | Participant 06 | Impact | Parietal | Aperiodic exponent | 24 | 0.6706 | 0.7999 | 0.6307 | 0.7692 | 0.1293 | 0.0945 | 0.1829 | 0.7071 | 3.4640 | 0.002 | 0.001 | 5 | 19 | 0 | 20.8333 | 79.1667 | 0.005 | 0.003 |
| Initial short-window specification | Participant 07 | Impact | Parietal | Aperiodic exponent | 24 | 0.6426 | 0.7841 | 0.6090 | 0.7470 | 0.1415 | 0.1051 | 0.1760 | 0.8042 | 3.9396 | <0.001 | <0.001 | 4 | 20 | 0 | 16.6667 | 83.3333 | 0.002 | <0.001 |
| Initial short-window specification | Participant 08 | Impact | Parietal | Aperiodic exponent | 24 | 0.6711 | 0.8094 | 0.6307 | 0.7692 | 0.1383 | 0.1051 | 0.1800 | 0.7681 | 3.7630 | 0.001 | <0.001 | 4 | 20 | 0 | 16.6667 | 83.3333 | 0.003 | 0.002 |
| Initial short-window specification | Participant 09 | Impact | Parietal | Aperiodic exponent | 24 | 0.6703 | 0.8035 | 0.6307 | 0.7692 | 0.1332 | 0.1051 | 0.1830 | 0.7278 | 3.5653 | 0.002 | 0.001 | 5 | 19 | 0 | 20.8333 | 79.1667 | 0.004 | 0.003 |
| Initial short-window specification | Participant 10 | Impact | Parietal | Aperiodic exponent | 24 | 0.6487 | 0.7860 | 0.6090 | 0.7692 | 0.1373 | 0.1051 | 0.1809 | 0.7590 | 3.7182 | 0.001 | <0.001 | 4 | 20 | 0 | 16.6667 | 83.3333 | 0.003 | 0.002 |
| Initial short-window specification | Participant 11 | Impact | Parietal | Aperiodic exponent | 24 | 0.6552 | 0.7658 | 0.6307 | 0.7470 | 0.1107 | 0.0945 | 0.1491 | 0.7421 | 3.6355 | 0.001 | 0.001 | 5 | 19 | 0 | 20.8333 | 79.1667 | 0.004 | 0.004 |
| Initial short-window specification | Participant 12 | Impact | Parietal | Aperiodic exponent | 24 | 0.6688 | 0.8039 | 0.6307 | 0.7692 | 0.1352 | 0.1051 | 0.1823 | 0.7414 | 3.6321 | 0.001 | <0.001 | 5 | 19 | 0 | 20.8333 | 79.1667 | 0.004 | 0.002 |
| Initial short-window specification | Participant 13 | Impact | Parietal | Aperiodic exponent | 24 | 0.6490 | 0.7732 | 0.6090 | 0.7470 | 0.1242 | 0.0945 | 0.1794 | 0.6923 | 3.3915 | 0.003 | 0.001 | 5 | 19 | 0 | 20.8333 | 79.1667 | 0.006 | 0.004 |
| Initial short-window specification | Participant 14 | Impact | Parietal | Aperiodic exponent | 24 | 0.6537 | 0.7897 | 0.6131 | 0.7692 | 0.1360 | 0.1051 | 0.1818 | 0.7481 | 3.6648 | 0.001 | <0.001 | 5 | 19 | 0 | 20.8333 | 79.1667 | 0.003 | 0.002 |
| Initial short-window specification | Participant 15 | Impact | Parietal | Aperiodic exponent | 24 | 0.6534 | 0.7880 | 0.6090 | 0.7692 | 0.1347 | 0.1051 | 0.1825 | 0.7377 | 3.6141 | 0.001 | <0.001 | 5 | 19 | 0 | 20.8333 | 79.1667 | 0.004 | 0.002 |
| Initial short-window specification | Participant 16 | Impact | Parietal | Aperiodic exponent | 24 | 0.6391 | 0.7799 | 0.6090 | 0.7470 | 0.1408 | 0.1051 | 0.1771 | 0.7950 | 3.8949 | <0.001 | <0.001 | 4 | 20 | 0 | 16.6667 | 83.3333 | 0.002 | <0.001 |
| Initial short-window specification | Participant 17 | Impact | Parietal | Aperiodic exponent | 24 | 0.6565 | 0.7861 | 0.6307 | 0.7692 | 0.1295 | 0.0945 | 0.1830 | 0.7079 | 3.4681 | 0.002 | 0.001 | 5 | 19 | 0 | 20.8333 | 79.1667 | 0.005 | 0.003 |
| Initial short-window specification | Participant 18 | Impact | Parietal | Aperiodic exponent | 24 | 0.6578 | 0.7907 | 0.6307 | 0.7692 | 0.1329 | 0.1018 | 0.1831 | 0.7259 | 3.5560 | 0.002 | 0.001 | 5 | 19 | 0 | 20.8333 | 79.1667 | 0.004 | 0.003 |
| Initial short-window specification | Participant 19 | Impact | Parietal | Aperiodic exponent | 24 | 0.6589 | 0.7921 | 0.6307 | 0.7692 | 0.1332 | 0.1051 | 0.1830 | 0.7276 | 3.5647 | 0.002 | 0.001 | 5 | 19 | 0 | 20.8333 | 79.1667 | 0.004 | 0.003 |
| Initial short-window specification | Participant 20 | Impact | Parietal | Aperiodic exponent | 24 | 0.6398 | 0.7710 | 0.6090 | 0.7470 | 0.1312 | 0.0945 | 0.1832 | 0.7160 | 3.5076 | 0.002 | 0.001 | 5 | 19 | 0 | 20.8333 | 79.1667 | 0.005 | 0.003 |
| Initial short-window specification | Participant 21 | Impact | Parietal | Aperiodic exponent | 24 | 0.6354 | 0.7615 | 0.6090 | 0.7470 | 0.1261 | 0.0945 | 0.1812 | 0.6961 | 3.4103 | 0.002 | 0.001 | 5 | 19 | 0 | 20.8333 | 79.1667 | 0.006 | 0.004 |
| Initial short-window specification | Participant 22 | Impact | Parietal | Aperiodic exponent | 24 | 0.6696 | 0.8149 | 0.6307 | 0.7692 | 0.1453 | 0.1051 | 0.1693 | 0.8581 | 4.2036 | <0.001 | <0.001 | 4 | 20 | 0 | 16.6667 | 83.3333 | <0.001 | <0.001 |
| Initial short-window specification | Participant 23 | Impact | Parietal | Aperiodic exponent | 24 | 0.6271 | 0.7589 | 0.6090 | 0.7470 | 0.1318 | 0.0945 | 0.1832 | 0.7192 | 3.5234 | 0.002 | 0.001 | 5 | 19 | 0 | 20.8333 | 79.1667 | 0.004 | 0.003 |
| Initial short-window specification | Participant 24 | Impact | Parietal | Aperiodic exponent | 24 | 0.6505 | 0.7854 | 0.6090 | 0.7629 | 0.1350 | 0.1051 | 0.1824 | 0.7400 | 3.6254 | 0.001 | <0.001 | 5 | 19 | 0 | 20.8333 | 79.1667 | 0.004 | 0.002 |
| Initial short-window specification | Participant 25 | Impact | Parietal | Aperiodic exponent | 24 | 0.6319 | 0.7642 | 0.6090 | 0.7470 | 0.1323 | 0.0945 | 0.1832 | 0.7222 | 3.5380 | 0.002 | 0.001 | 5 | 19 | 0 | 20.8333 | 79.1667 | 0.005 | 0.003 |
| Resolution-aware max6 | Participant 01 | Impact | Parietal | Aperiodic exponent | 24 | 0.6464 | 0.7805 | 0.6248 | 0.7527 | 0.1341 | 0.0948 | 0.1843 | 0.7278 | 3.5656 | 0.002 | 0.001 | 5 | 19 | 0 | 20.8333 | 79.1667 | 0.004 | 0.004 |
| Resolution-aware max6 | Participant 02 | Impact | Parietal | Aperiodic exponent | 24 | 0.6460 | 0.7766 | 0.6248 | 0.7527 | 0.1306 | 0.0948 | 0.1800 | 0.7257 | 3.5553 | 0.002 | 0.001 | 5 | 19 | 0 | 20.8333 | 79.1667 | 0.004 | 0.004 |
| Resolution-aware max6 | Participant 03 | Impact | Parietal | Aperiodic exponent | 24 | 0.6528 | 0.7964 | 0.6248 | 0.7692 | 0.1436 | 0.1061 | 0.1870 | 0.7682 | 3.7636 | 0.001 | <0.001 | 5 | 19 | 0 | 20.8333 | 79.1667 | 0.003 | 0.002 |
| Resolution-aware max6 | Participant 04 | Impact | Parietal | Aperiodic exponent | 24 | 0.6327 | 0.7722 | 0.6101 | 0.7527 | 0.1395 | 0.0948 | 0.1874 | 0.7444 | 3.6468 | 0.001 | 0.001 | 5 | 19 | 0 | 20.8333 | 79.1667 | 0.003 | 0.003 |
| Resolution-aware max6 | Participant 05 | Impact | Parietal | Aperiodic exponent | 24 | 0.6325 | 0.7613 | 0.6101 | 0.7527 | 0.1288 | 0.0948 | 0.1770 | 0.7276 | 3.5647 | 0.002 | 0.001 | 5 | 19 | 0 | 20.8333 | 79.1667 | 0.004 | 0.004 |
| Resolution-aware max6 | Participant 06 | Impact | Parietal | Aperiodic exponent | 24 | 0.6590 | 0.7982 | 0.6248 | 0.7692 | 0.1392 | 0.0948 | 0.1873 | 0.7432 | 3.6407 | 0.001 | 0.001 | 5 | 19 | 0 | 20.8333 | 79.1667 | 0.004 | 0.003 |
| Resolution-aware max6 | Participant 07 | Impact | Parietal | Aperiodic exponent | 24 | 0.6312 | 0.7824 | 0.6101 | 0.7527 | 0.1512 | 0.1061 | 0.1801 | 0.8395 | 4.1128 | <0.001 | <0.001 | 4 | 20 | 0 | 16.6667 | 83.3333 | 0.001 | <0.001 |
| Resolution-aware max6 | Participant 08 | Impact | Parietal | Aperiodic exponent | 24 | 0.6603 | 0.8076 | 0.6248 | 0.7692 | 0.1473 | 0.1061 | 0.1846 | 0.7978 | 3.9085 | <0.001 | <0.001 | 4 | 20 | 0 | 16.6667 | 83.3333 | 0.002 | 0.002 |
| Resolution-aware max6 | Participant 09 | Impact | Parietal | Aperiodic exponent | 24 | 0.6588 | 0.8018 | 0.6248 | 0.7692 | 0.1430 | 0.1061 | 0.1872 | 0.7639 | 3.7424 | 0.001 | <0.001 | 5 | 19 | 0 | 20.8333 | 79.1667 | 0.003 | 0.002 |
| Resolution-aware max6 | Participant 10 | Impact | Parietal | Aperiodic exponent | 24 | 0.6369 | 0.7838 | 0.6101 | 0.7692 | 0.1469 | 0.1061 | 0.1850 | 0.7940 | 3.8900 | <0.001 | <0.001 | 4 | 20 | 0 | 16.6667 | 83.3333 | 0.002 | 0.002 |
| Resolution-aware max6 | Participant 11 | Impact | Parietal | Aperiodic exponent | 24 | 0.6431 | 0.7641 | 0.6248 | 0.7527 | 0.1210 | 0.0948 | 0.1576 | 0.7678 | 3.7615 | 0.001 | 0.001 | 5 | 19 | 0 | 20.8333 | 79.1667 | 0.003 | 0.004 |
| Resolution-aware max6 | Participant 12 | Impact | Parietal | Aperiodic exponent | 24 | 0.6573 | 0.8026 | 0.6248 | 0.7692 | 0.1453 | 0.1061 | 0.1861 | 0.7807 | 3.8248 | <0.001 | <0.001 | 5 | 19 | 0 | 20.8333 | 79.1667 | 0.002 | 0.002 |
| Resolution-aware max6 | Participant 13 | Impact | Parietal | Aperiodic exponent | 24 | 0.6375 | 0.7714 | 0.6101 | 0.7527 | 0.1340 | 0.0948 | 0.1841 | 0.7276 | 3.5647 | 0.002 | 0.001 | 5 | 19 | 0 | 20.8333 | 79.1667 | 0.004 | 0.004 |
| Resolution-aware max6 | Participant 14 | Impact | Parietal | Aperiodic exponent | 24 | 0.6427 | 0.7884 | 0.6201 | 0.7692 | 0.1457 | 0.1061 | 0.1859 | 0.7841 | 3.8413 | <0.001 | <0.001 | 5 | 19 | 0 | 20.8333 | 79.1667 | 0.003 | 0.002 |
| Resolution-aware max6 | Participant 15 | Impact | Parietal | Aperiodic exponent | 24 | 0.6419 | 0.7863 | 0.6101 | 0.7692 | 0.1445 | 0.1061 | 0.1866 | 0.7740 | 3.7919 | <0.001 | <0.001 | 5 | 19 | 0 | 20.8333 | 79.1667 | 0.002 | 0.002 |
| Resolution-aware max6 | Participant 16 | Impact | Parietal | Aperiodic exponent | 24 | 0.6277 | 0.7784 | 0.6101 | 0.7527 | 0.1506 | 0.1061 | 0.1808 | 0.8328 | 4.0800 | <0.001 | <0.001 | 4 | 20 | 0 | 16.6667 | 83.3333 | 0.001 | <0.001 |
| Resolution-aware max6 | Participant 17 | Impact | Parietal | Aperiodic exponent | 24 | 0.6450 | 0.7850 | 0.6248 | 0.7692 | 0.1400 | 0.0948 | 0.1874 | 0.7469 | 3.6588 | 0.001 | 0.001 | 5 | 19 | 0 | 20.8333 | 79.1667 | 0.003 | 0.003 |
| Resolution-aware max6 | Participant 18 | Impact | Parietal | Aperiodic exponent | 24 | 0.6463 | 0.7890 | 0.6248 | 0.7692 | 0.1427 | 0.1019 | 0.1873 | 0.7617 | 3.7315 | 0.001 | 0.001 | 5 | 19 | 0 | 20.8333 | 79.1667 | 0.003 | 0.003 |
| Resolution-aware max6 | Participant 19 | Impact | Parietal | Aperiodic exponent | 24 | 0.6475 | 0.7905 | 0.6248 | 0.7692 | 0.1430 | 0.1061 | 0.1872 | 0.7641 | 3.7435 | 0.001 | <0.001 | 5 | 19 | 0 | 20.8333 | 79.1667 | 0.003 | 0.002 |
| Resolution-aware max6 | Participant 20 | Impact | Parietal | Aperiodic exponent | 24 | 0.6283 | 0.7694 | 0.6101 | 0.7527 | 0.1411 | 0.0948 | 0.1875 | 0.7523 | 3.6857 | 0.001 | 0.001 | 5 | 19 | 0 | 20.8333 | 79.1667 | 0.003 | 0.003 |
| Resolution-aware max6 | Participant 21 | Impact | Parietal | Aperiodic exponent | 24 | 0.6239 | 0.7598 | 0.6101 | 0.7527 | 0.1359 | 0.0948 | 0.1858 | 0.7316 | 3.5842 | 0.002 | 0.001 | 5 | 19 | 0 | 20.8333 | 79.1667 | 0.004 | 0.004 |
| Resolution-aware max6 | Participant 22 | Impact | Parietal | Aperiodic exponent | 24 | 0.6581 | 0.8131 | 0.6248 | 0.7692 | 0.1550 | 0.1061 | 0.1731 | 0.8959 | 4.3890 | <0.001 | <0.001 | 4 | 20 | 0 | 16.6667 | 83.3333 | <0.001 | <0.001 |
| Resolution-aware max6 | Participant 23 | Impact | Parietal | Aperiodic exponent | 24 | 0.6242 | 0.7571 | 0.6101 | 0.7527 | 0.1329 | 0.0948 | 0.1830 | 0.7263 | 3.5581 | 0.002 | 0.001 | 5 | 19 | 0 | 20.8333 | 79.1667 | 0.004 | 0.003 |
| Resolution-aware max6 | Participant 24 | Impact | Parietal | Aperiodic exponent | 24 | 0.6389 | 0.7837 | 0.6101 | 0.7686 | 0.1448 | 0.1061 | 0.1865 | 0.7763 | 3.8031 | <0.001 | <0.001 | 5 | 19 | 0 | 20.8333 | 79.1667 | 0.003 | 0.002 |
| Resolution-aware max6 | Participant 25 | Impact | Parietal | Aperiodic exponent | 24 | 0.6204 | 0.7624 | 0.6101 | 0.7527 | 0.1421 | 0.0948 | 0.1874 | 0.7580 | 3.7137 | 0.001 | 0.001 | 5 | 19 | 0 | 20.8333 | 79.1667 | 0.003 | 0.003 |
| Resolution-aware max3 | Participant 01 | Impact | Parietal | Aperiodic exponent | 24 | 0.6550 | 0.7752 | 0.6248 | 0.7527 | 0.1202 | 0.0948 | 0.1728 | 0.6957 | 3.4082 | 0.002 | 0.002 | 5 | 19 | 0 | 20.8333 | 79.1667 | 0.006 | 0.004 |
| Resolution-aware max3 | Participant 02 | Impact | Parietal | Aperiodic exponent | 24 | 0.6545 | 0.7712 | 0.6248 | 0.7527 | 0.1167 | 0.0948 | 0.1679 | 0.6951 | 3.4052 | 0.002 | 0.002 | 5 | 19 | 0 | 20.8333 | 79.1667 | 0.005 | 0.004 |
| Resolution-aware max3 | Participant 03 | Impact | Parietal | Aperiodic exponent | 24 | 0.6613 | 0.7911 | 0.6248 | 0.7692 | 0.1298 | 0.1053 | 0.1765 | 0.7352 | 3.6018 | 0.002 | 0.001 | 5 | 19 | 0 | 20.8333 | 79.1667 | 0.004 | 0.003 |
| Resolution-aware max3 | Participant 04 | Impact | Parietal | Aperiodic exponent | 24 | 0.6412 | 0.7675 | 0.6101 | 0.7527 | 0.1263 | 0.0948 | 0.1767 | 0.7145 | 3.5003 | 0.002 | 0.001 | 5 | 19 | 0 | 20.8333 | 79.1667 | 0.004 | 0.003 |
| Resolution-aware max3 | Participant 05 | Impact | Parietal | Aperiodic exponent | 24 | 0.6411 | 0.7603 | 0.6101 | 0.7527 | 0.1192 | 0.0948 | 0.1716 | 0.6947 | 3.4033 | 0.002 | 0.002 | 5 | 19 | 0 | 20.8333 | 79.1667 | 0.006 | 0.004 |
| Resolution-aware max3 | Participant 06 | Impact | Parietal | Aperiodic exponent | 24 | 0.6676 | 0.7928 | 0.6248 | 0.7692 | 0.1252 | 0.0948 | 0.1765 | 0.7098 | 3.4772 | 0.002 | 0.001 | 5 | 19 | 0 | 20.8333 | 79.1667 | 0.005 | 0.004 |
| Resolution-aware max3 | Participant 07 | Impact | Parietal | Aperiodic exponent | 24 | 0.6398 | 0.7770 | 0.6101 | 0.7527 | 0.1373 | 0.1053 | 0.1698 | 0.8085 | 3.9610 | <0.001 | <0.001 | 4 | 20 | 0 | 16.6667 | 83.3333 | 0.001 | <0.001 |

| Model | Excluded participant | Window | Sensor region | Feature | Participants | Human mean | Machine mean | Human median | Machine median | Machine − human mean | Machine − human median | Difference SD | Cohen's dz | Paired t | Paired t p | Wilcoxon p | Machine lower, n | Machine higher, n | Equal, n | Machine lower (%) | Machine higher (%) | Paired t FDR q | Wilcoxon FDR q |
|---|---|---|---|---|---|---|---|---|---|---|---|---|---|---|---|---|---|---|---|---|---|---|---|
| Resolution-aware max3 | Participant 08 | Impact | Parietal | Aperiodic exponent | 24 | 0.6689 | 0.8025 | 0.6248 | 0.7692 | 0.1337 | 0.1053 | 0.1741 | 0.7677 | 3.7610 | 0.001 | <0.001 | 4 | 20 | 0 | 16.6667 | 83.3333 | 0.002 | 0.002 |
| Resolution-aware max3 | Participant 09 | Impact | Parietal | Aperiodic exponent | 24 | 0.6673 | 0.7964 | 0.6248 | 0.7692 | 0.1291 | 0.1053 | 0.1767 | 0.7308 | 3.5803 | 0.002 | 0.001 | 5 | 19 | 0 | 20.8333 | 79.1667 | 0.004 | 0.003 |
| Resolution-aware max3 | Participant 10 | Impact | Parietal | Aperiodic exponent | 24 | 0.6454 | 0.7784 | 0.6101 | 0.7692 | 0.1330 | 0.1053 | 0.1747 | 0.7615 | 3.7307 | 0.001 | <0.001 | 4 | 20 | 0 | 16.6667 | 83.3333 | 0.002 | 0.002 |
| Resolution-aware max3 | Participant 11 | Impact | Parietal | Aperiodic exponent | 24 | 0.6516 | 0.7588 | 0.6248 | 0.7527 | 0.1072 | 0.0948 | 0.1428 | 0.7505 | 3.6768 | 0.001 | 0.002 | 5 | 19 | 0 | 20.8333 | 79.1667 | 0.003 | 0.004 |
| Resolution-aware max3 | Participant 12 | Impact | Parietal | Aperiodic exponent | 24 | 0.6658 | 0.7973 | 0.6248 | 0.7692 | 0.1314 | 0.1053 | 0.1758 | 0.7479 | 3.6639 | 0.001 | 0.001 | 5 | 19 | 0 | 20.8333 | 79.1667 | 0.003 | 0.003 |
| Resolution-aware max3 | Participant 13 | Impact | Parietal | Aperiodic exponent | 24 | 0.6460 | 0.7661 | 0.6101 | 0.7527 | 0.1201 | 0.0948 | 0.1727 | 0.6955 | 3.4074 | 0.002 | 0.002 | 5 | 19 | 0 | 20.8333 | 79.1667 | 0.006 | 0.004 |
| Resolution-aware max3 | Participant 14 | Impact | Parietal | Aperiodic exponent | 24 | 0.6512 | 0.7831 | 0.6201 | 0.7692 | 0.1319 | 0.1053 | 0.1755 | 0.7514 | 3.6809 | 0.001 | 0.001 | 5 | 19 | 0 | 20.8333 | 79.1667 | 0.003 | 0.003 |
| Resolution-aware max3 | Participant 15 | Impact | Parietal | Aperiodic exponent | 24 | 0.6504 | 0.7810 | 0.6101 | 0.7692 | 0.1306 | 0.1053 | 0.1762 | 0.7411 | 3.6306 | 0.001 | 0.001 | 5 | 19 | 0 | 20.8333 | 79.1667 | 0.003 | 0.002 |
| Resolution-aware max3 | Participant 16 | Impact | Parietal | Aperiodic exponent | 24 | 0.6363 | 0.7730 | 0.6101 | 0.7527 | 0.1367 | 0.1053 | 0.1706 | 0.8016 | 3.9269 | <0.001 | <0.001 | 4 | 20 | 0 | 16.6667 | 83.3333 | 0.002 | <0.001 |
| Resolution-aware max3 | Participant 17 | Impact | Parietal | Aperiodic exponent | 24 | 0.6536 | 0.7797 | 0.6248 | 0.7692 | 0.1261 | 0.0948 | 0.1767 | 0.7137 | 3.4966 | 0.002 | 0.001 | 5 | 19 | 0 | 20.8333 | 79.1667 | 0.004 | 0.004 |
| Resolution-aware max3 | Participant 18 | Impact | Parietal | Aperiodic exponent | 24 | 0.6548 | 0.7836 | 0.6248 | 0.7692 | 0.1288 | 0.1012 | 0.1768 | 0.7286 | 3.5694 | 0.002 | 0.001 | 5 | 19 | 0 | 20.8333 | 79.1667 | 0.004 | 0.003 |
| Resolution-aware max3 | Participant 19 | Impact | Parietal | Aperiodic exponent | 24 | 0.6560 | 0.7852 | 0.6248 | 0.7692 | 0.1292 | 0.1053 | 0.1767 | 0.7311 | 3.5815 | 0.002 | 0.001 | 5 | 19 | 0 | 20.8333 | 79.1667 | 0.004 | 0.003 |
| Resolution-aware max3 | Participant 20 | Impact | Parietal | Aperiodic exponent | 24 | 0.6369 | 0.7641 | 0.6101 | 0.7527 | 0.1272 | 0.0948 | 0.1768 | 0.7192 | 3.5233 | 0.002 | 0.001 | 5 | 19 | 0 | 20.8333 | 79.1667 | 0.004 | 0.004 |
| Resolution-aware max3 | Participant 21 | Impact | Parietal | Aperiodic exponent | 24 | 0.6324 | 0.7545 | 0.6101 | 0.7527 | 0.1220 | 0.0948 | 0.1746 | 0.6990 | 3.4245 | 0.002 | 0.002 | 5 | 19 | 0 | 20.8333 | 79.1667 | 0.006 | 0.004 |
| Resolution-aware max3 | Participant 22 | Impact | Parietal | Aperiodic exponent | 24 | 0.6666 | 0.8078 | 0.6248 | 0.7692 | 0.1412 | 0.1053 | 0.1627 | 0.8676 | 4.2506 | <0.001 | <0.001 | 4 | 20 | 0 | 16.6667 | 83.3333 | 0.001 | <0.001 |
| Resolution-aware max3 | Participant 23 | Impact | Parietal | Aperiodic exponent | 24 | 0.6241 | 0.7518 | 0.6101 | 0.7527 | 0.1277 | 0.0948 | 0.1769 | 0.7221 | 3.5374 | 0.002 | 0.001 | 5 | 19 | 0 | 20.8333 | 79.1667 | 0.004 | 0.003 |
| Resolution-aware max3 | Participant 24 | Impact | Parietal | Aperiodic exponent | 24 | 0.6475 | 0.7784 | 0.6101 | 0.7686 | 0.1309 | 0.1053 | 0.1761 | 0.7434 | 3.6419 | 0.001 | 0.001 | 5 | 19 | 0 | 20.8333 | 79.1667 | 0.004 | 0.003 |
| Resolution-aware max3 | Participant 25 | Impact | Parietal | Aperiodic exponent | 24 | 0.6289 | 0.7572 | 0.6101 | 0.7527 | 0.1283 | 0.0948 | 0.1768 | 0.7253 | 3.5533 | 0.002 | 0.001 | 5 | 19 | 0 | 20.8333 | 79.1667 | 0.004 | 0.004 |

### Section 4. Short-window model concordance

| Participants | Model A | Model B | Pearson r | Spearman ρ | Mean difference between model effects | Median difference between model effects | Mean absolute effect difference | Maximum absolute effect difference | Participants changing direction, n | Direction agreement fraction |
|---|---|---|---|---|---|---|---|---|---|---|
| 25 | Initial short-window specification | Resolution-aware max6 | 0.9722 | 0.9769 | 0.0094 | 0.0000 | 0.0134 | 0.2082 | 0 | 1.0000 |
| 25 | Initial short-window specification | Resolution-aware max3 | 0.9909 | 0.9985 | -0.0039 | 0.0000 | 0.0096 | 0.1080 | 0 | 1.0000 |
| 25 | Resolution-aware max6 | Resolution-aware max3 | 0.9690 | 0.9769 | -0.0133 | 0.0000 | 0.0135 | 0.2082 | 0 | 1.0000 |

### Section 5. Max6-versus-max3 BIC and residual summary

| Window | Sensor region | Grouping | Model cells | Mean peaks, max6 | Mean peaks, max3 | Mean Δ peaks (max3 − max6) | Mean ΔBIC (max3 − max6) | Median ΔBIC (max3 − max6) | ΔBIC Q1 | ΔBIC Q3 | Max3 lower BIC, n | Max6 lower BIC, n | BIC ties, n | Fraction max3 lower BIC | Fraction max6 lower BIC | Mean $\Delta R^2$ (max3 − max6) | Mean ΔMAE (max3 − max6) | Mean ΔNRMSE (max3 − max6) | Mean absolute exponent change | Max6 boundary crossings, n | Max3 boundary crossings, n | Context |
|---|---|---|---|---|---|---|---|---|---|---|---|---|---|---|---|---|---|---|---|---|---|---|
| — | — | Overall | 900 | 1.4489 | 1.0911 | -0.3578 | 0.3517 | 0.0000 | 0.0000 | 0.0000 | 140 | 85 | 675 | 0.1556 | 0.0944 | -0.0052 | 0.0028 | 0.0030 | 0.0152 | 365 | 255 | — |
| Pre-hit | — | Window | 300 | 1.7667 | 1.1867 | -0.5800 | 0.1698 | 0.0000 | 0.0000 | 0.0000 | 71 | 39 | 190 | 0.2367 | 0.1300 | -0.0071 | 0.0040 | 0.0043 | 0.0197 | 147 | 92 | — |
| Impact | — | Window | 300 | 1.1167 | 0.9567 | -0.1600 | 0.6606 | 0.0000 | 0.0000 | 0.0000 | 18 | 20 | 262 | 0.0600 | 0.0667 | -0.0017 | 0.0019 | 0.0018 | 0.0114 | 106 | 84 | — |
| Post-hit | — | Window | 300 | 1.4633 | 1.1300 | -0.3333 | 0.2248 | 0.0000 | 0.0000 | 0.0000 | 51 | 26 | 223 | 0.1700 | 0.0867 | -0.0068 | 0.0025 | 0.0030 | 0.0145 | 112 | 79 | — |
| Pre-hit | — | Window × context | 150 | 1.6600 | 1.1267 | -0.5333 | -0.5576 | 0.0000 | 0.0000 | 0.0000 | 37 | 15 | 98 | 0.2467 | 0.1000 | -0.0073 | 0.0035 | 0.0038 | 0.0176 | 67 | 41 | Human opponent |
| Pre-hit | — | Window × context | 150 | 1.8733 | 1.2467 | -0.6267 | 0.8973 | 0.0000 | 0.0000 | 0.0000 | 34 | 24 | 92 | 0.2267 | 0.1600 | -0.0070 | 0.0044 | 0.0049 | 0.0219 | 80 | 51 | Ball machine |
| Impact | — | Window × context | 150 | 1.1067 | 0.9533 | -0.1533 | 0.6687 | 0.0000 | 0.0000 | 0.0000 | 9 | 8 | 133 | 0.0600 | 0.0533 | -0.0018 | 0.0017 | 0.0018 | 0.0096 | 49 | 38 | Human opponent |
| Impact | — | Window × context | 150 | 1.1267 | 0.9600 | -0.1667 | 0.6525 | 0.0000 | 0.0000 | 0.0000 | 9 | 12 | 129 | 0.0600 | 0.0800 | -0.0017 | 0.0021 | 0.0018 | 0.0133 | 57 | 46 | Ball machine |
| Post-hit | — | Window × context | 150 | 1.3667 | 1.0733 | -0.2933 | 0.3103 | 0.0000 | 0.0000 | 0.0000 | 22 | 12 | 116 | 0.1467 | 0.0800 | -0.0080 | 0.0022 | 0.0030 | 0.0133 | 52 | 38 | Human opponent |
| Post-hit | — | Window × context | 150 | 1.5600 | 1.1867 | -0.3733 | 0.1393 | 0.0000 | 0.0000 | 0.0000 | 29 | 14 | 107 | 0.1933 | 0.0933 | -0.0056 | 0.0029 | 0.0031 | 0.0157 | 60 | 41 | Ball machine |
| Pre-hit | Frontal | Window × region | 50 | 2.1000 | 0.9800 | -1.1200 | 1.1020 | -0.6113 | -6.3850 | 0.0000 | 25 | 10 | 15 | 0.5000 | 0.2000 | -0.0100 | 0.0088 | 0.0073 | 0.0461 | 52 | 31 | — |
| Pre-hit | Sensorimotor | Window × region | 50 | 2.1400 | 1.4400 | -0.7000 | 1.1259 | 0.0000 | -2.5825 | 0.0000 | 13 | 9 | 28 | 0.2600 | 0.1800 | -0.0088 | 0.0047 | 0.0055 | 0.0233 | 30 | 18 | — |
| Pre-hit | Parietal | Window × region | 50 | 1.5600 | 1.1200 | -0.4400 | 1.1732 | 0.0000 | 0.0000 | 0.0000 | 6 | 8 | 36 | 0.1200 | 0.1600 | -0.0067 | 0.0032 | 0.0040 | 0.0175 | 18 | 11 | — |
| Pre-hit | Posterior | Window × region | 50 | 1.5000 | 1.2400 | -0.2600 | 0.0050 | 0.0000 | 0.0000 | 0.0000 | 4 | 5 | 41 | 0.0800 | 0.1000 | -0.0028 | 0.0014 | 0.0020 | 0.0059 | 16 | 12 | — |
| Pre-hit | Temporal | Window × region | 50 | 1.4400 | 1.0200 | -0.4200 | -1.7565 | 0.0000 | 0.0000 | 0.0000 | 11 | 3 | 36 | 0.2200 | 0.0600 | -0.0073 | 0.0021 | 0.0028 | 0.0078 | 14 | 10 | — |
| Pre-hit | Other | Window × region | 50 | 1.8600 | 1.3200 | -0.5400 | -0.6306 | 0.0000 | 0.0000 | 0.0000 | 12 | 4 | 34 | 0.2400 | 0.0800 | -0.0071 | 0.0035 | 0.0044 | 0.0179 | 17 | 10 | — |
| Impact | Frontal | Window × region | 50 | 1.0800 | 0.7800 | -0.3000 | 3.3600 | 0.0000 | 0.0000 | 0.0000 | 3 | 8 | 39 | 0.0600 | 0.1600 | -0.0030 | 0.0056 | 0.0043 | 0.0361 | 38 | 29 | — |
| Impact | Sensorimotor | Window × region | 50 | 1.0200 | 0.9600 | -0.0600 | 0.8383 | 0.0000 | 0.0000 | 0.0000 | 1 | 2 | 47 | 0.0200 | 0.0400 | -0.0015 | 0.0011 | 0.0015 | 0.0086 | 13 | 11 | — |
| Impact | Parietal | Window × region | 50 | 1.2400 | 1.0400 | -0.2000 | -0.3158 | 0.0000 | 0.0000 | 0.0000 | 6 | 2 | 42 | 0.1200 | 0.0400 | -0.0019 | 0.0013 | 0.0013 | 0.0068 | 14 | 11 | — |
| Impact | Posterior | Window × region | 50 | 1.2400 | 1.0600 | -0.1800 | 0.5693 | 0.0000 | 0.0000 | 0.0000 | 2 | 5 | 43 | 0.0400 | 0.1000 | -0.0023 | 0.0019 | 0.0021 | 0.0102 | 18 | 13 | — |
| Impact | Temporal | Window × region | 50 | 0.8200 | 0.7200 | -0.1000 | -0.0810 | 0.0000 | 0.0000 | 0.0000 | 3 | 2 | 45 | 0.0600 | 0.0400 | -0.0011 | 0.0007 | 0.0009 | 0.0028 | 10 | 8 | — |
| Impact | Other | Window × region | 50 | 1.3000 | 1.1800 | -0.1200 | -0.4076 | 0.0000 | 0.0000 | 0.0000 | 3 | 1 | 46 | 0.0600 | 0.0200 | -0.0005 | 0.0007 | 0.0006 | 0.0040 | 13 | 12 | — |
| Post-hit | Frontal | Window × region | 50 | 1.3800 | 0.7800 | -0.6000 | 1.4317 | 0.0000 | -1.6923 | 0.0000 | 14 | 6 | 30 | 0.2800 | 0.1200 | -0.0050 | 0.0055 | 0.0044 | 0.0369 | 36 | 25 | — |
| Post-hit | Sensorimotor | Window × region | 50 | 1.6000 | 1.2200 | -0.3800 | -0.5503 | 0.0000 | 0.0000 | 0.0000 | 10 | 5 | 35 | 0.2000 | 0.1000 | -0.0063 | 0.0022 | 0.0029 | 0.0140 | 21 | 15 | — |
| Post-hit | Parietal | Window × region | 50 | 1.5600 | 1.2200 | -0.3400 | -0.2207 | 0.0000 | 0.0000 | 0.0000 | 10 | 4 | 36 | 0.2000 | 0.0800 | -0.0028 | 0.0016 | 0.0020 | 0.0097 | 16 | 12 | — |
| Post-hit | Posterior | Window × region | 50 | 1.4000 | 1.1600 | -0.2400 | 0.6313 | 0.0000 | 0.0000 | 0.0000 | 5 | 5 | 40 | 0.1000 | 0.1000 | -0.0061 | 0.0021 | 0.0032 | 0.0091 | 14 | 11 | — |
| Post-hit | Temporal | Window × region | 50 | 1.2000 | 1.0000 | -0.2000 | -0.3104 | 0.0000 | 0.0000 | 0.0000 | 5 | 2 | 43 | 0.1000 | 0.0400 | -0.0172 | 0.0019 | 0.0032 | 0.0088 | 11 | 7 | — |
| Post-hit | Other | Window × region | 50 | 1.6400 | 1.4000 | -0.2400 | 0.3673 | 0.0000 | 0.0000 | 0.0000 | 7 | 4 | 39 | 0.1400 | 0.0800 | -0.0035 | 0.0018 | 0.0025 | 0.0086 | 14 | 11 | — |

### Section 6. Focal-cell BIC summary

| Participants | Mean ΔBIC (max3 − max6) | Median ΔBIC (max3 − max6) | Max3 lower BIC, n | Max6 lower BIC, n | BIC ties, n | Mean $\Delta R^2$ (max3 − max6) | Mean ΔMAE (max3 − max6) | Mean absolute exponent change | Context |
|---|---|---|---|---|---|---|---|---|---|
| 25 | -0.3027 | 0.0000 | 3 | 1 | 21 | -0.0021 | 0.0013 | 0.0085 | Human opponent |
| 25 | -0.3288 | 0.0000 | 3 | 1 | 21 | -0.0016 | 0.0014 | 0.0051 | Ball machine |

### Section 7. BIC condition-balance tests

| Window | Sensor region | Participants | Paired t | Paired t p | Wilcoxon p | Human mean ΔBIC | Machine mean ΔBIC | Machine − human mean ΔBIC | Human: max3 preferred, n | Machine: max3 preferred, n | Human only: max3 preferred, n | Machine only: max3 preferred, n | Discordant preference pairs, n | Exact preference-balance p | Paired t FDR q | Wilcoxon FDR q | Exact preference-balance FDR q |
|---|---|---|---|---|---|---|---|---|---|---|---|---|---|---|---|---|---|
| Pre-hit | Frontal | 25 | -0.0674 | 0.947 | 0.876 | 1.1980 | 1.0060 | -0.1920 | 13.0000 | 12.0000 | 5.0000 | 4.0000 | 9.0000 | 1.000 | 0.983 | 0.985 | 1.000 |
| Pre-hit | Sensorimotor | 25 | 2.1946 | 0.038 | 0.031 | -3.0112 | 5.2630 | 8.2742 | 7.0000 | 6.0000 | 4.0000 | 3.0000 | 7.0000 | 1.000 | 0.686 | 0.556 | 1.000 |
| Pre-hit | Parietal | 25 | -0.3533 | 0.727 | 0.477 | 1.6084 | 0.7380 | -0.8704 | 3.0000 | 3.0000 | 2.0000 | 2.0000 | 4.0000 | 1.000 | 0.983 | 0.974 | 1.000 |
| Pre-hit | Posterior | 25 | 0.6551 | 0.519 | 0.263 | -0.3179 | 0.3278 | 0.6456 | 2.0000 | 2.0000 | 1.0000 | 1.0000 | 2.0000 | 1.000 | 0.849 | 0.854 | 1.000 |
| Pre-hit | Temporal | 25 | 1.4671 | 0.155 | 0.155 | -2.9313 | -0.5816 | 2.3497 | 7.0000 | 4.0000 | 5.0000 | 2.0000 | 7.0000 | 0.453 | 0.849 | 0.854 | 1.000 |
| Pre-hit | Other | 25 | -0.6828 | 0.501 | 0.650 | 0.1084 | -1.3697 | -1.4781 | 5.0000 | 7.0000 | 4.0000 | 6.0000 | 10.0000 | 0.754 | 0.849 | 0.974 | 1.000 |
| Impact | Frontal | 25 | -0.2214 | 0.827 | 0.866 | 3.5074 | 3.2126 | -0.2948 | 2.0000 | 1.0000 | 2.0000 | 1.0000 | 3.0000 | 1.000 | 0.983 | 0.985 | 1.000 |
| Impact | Sensorimotor | 25 | -0.7351 | 0.469 | 0.593 | 1.4179 | 0.2586 | -1.1594 | 0.0000 | 1.0000 | 0.0000 | 1.0000 | 1.0000 | 1.000 | 0.849 | 0.974 | 1.000 |
| Impact | Parietal | 25 | -0.0216 | 0.983 | 0.866 | -0.3027 | -0.3288 | -0.0262 | 3.0000 | 3.0000 | 3.0000 | 3.0000 | 6.0000 | 1.000 | 0.983 | 0.985 | 1.000 |
| Impact | Posterior | 25 | 1.1340 | 0.268 | 0.249 | -0.2344 | 1.3731 | 1.6075 | 1.0000 | 1.0000 | 1.0000 | 1.0000 | 2.0000 | 1.000 | 0.849 | 0.854 | 1.000 |
| Impact | Temporal | 25 | -0.6669 | 0.511 | 0.715 | 0.0533 | -0.2152 | -0.2685 | 2.0000 | 1.0000 | 1.0000 | 0.0000 | 1.0000 | 1.000 | 0.849 | 0.985 | 1.000 |
| Impact | Other | 25 | 0.0554 | 0.956 | 1.000 | -0.4297 | -0.3855 | 0.0442 | 1.0000 | 2.0000 | 1.0000 | 2.0000 | 3.0000 | 1.000 | 0.983 | 1.000 | 1.000 |
| Post-hit | Frontal | 25 | -0.1639 | 0.871 | 0.650 | 1.6776 | 1.1859 | -0.4917 | 6.0000 | 8.0000 | 4.0000 | 6.0000 | 10.0000 | 0.754 | 0.983 | 0.974 | 1.000 |
| Post-hit | Sensorimotor | 25 | 0.9299 | 0.362 | 0.508 | -1.2908 | 0.1901 | 1.4809 | 6.0000 | 4.0000 | 4.0000 | 2.0000 | 6.0000 | 0.688 | 0.849 | 0.974 | 1.000 |
| Post-hit | Parietal | 25 | -0.9632 | 0.345 | 0.285 | 0.4706 | -0.9120 | -1.3825 | 4.0000 | 6.0000 | 2.0000 | 4.0000 | 6.0000 | 0.688 | 0.849 | 0.854 | 1.000 |
| Post-hit | Posterior | 25 | 0.6618 | 0.514 | 0.953 | 0.1508 | 1.1119 | 0.9611 | 1.0000 | 4.0000 | 1.0000 | 4.0000 | 5.0000 | 0.375 | 0.849 | 1.000 | 1.000 |
| Post-hit | Temporal | 25 | -1.0282 | 0.314 | 0.225 | 0.3797 | -1.0006 | -1.3804 | 2.0000 | 3.0000 | 1.0000 | 2.0000 | 3.0000 | 1.000 | 0.849 | 0.854 | 1.000 |

| Window | Sensor region | Participants | Paired t | Paired t p | Wilcoxon p | Human mean ΔBIC | Machine mean ΔBIC | Machine − human mean ΔBIC | Human: max3 preferred, n | Machine: max3 preferred, n | Human only: max3 preferred, n | Machine only: max3 preferred, n | Discordant preference pairs, n | Exact preference-balance p | Paired t FDR q | Wilcoxon FDR q | Exact preference-balance FDR q |
|---|---|---|---|---|---|---|---|---|---|---|---|---|---|---|---|---|---|
| Post-hit | Other | 25 | -0.1487 | 0.883 | 0.484 | 0.4740 | 0.2607 | -0.2133 | 3.0000 | 4.0000 | 2.0000 | 3.0000 | 5.0000 | 1.000 | 0.983 | 0.974 | 1.000 |

### Section 8. Two-second max6 fixed-family statistics

| Window | Sensor region | Feature | Participants | Human mean | Machine mean | Human median | Machine median | Machine − human mean | Machine − human median | Difference SD | Cohen's dz | Paired t | Paired t p | Wilcoxon p | Machine lower, n | Machine higher, n | Equal, n | Machine lower (%) | Machine higher (%) | Paired t FDR q | Wilcoxon FDR q |
|---|---|---|---|---|---|---|---|---|---|---|---|---|---|---|---|---|---|---|---|---|---|
| Two-second peri-hit | Frontal | Aperiodic exponent | 25 | 1.2864 | 1.3298 | 1.3397 | 1.3142 | 0.0434 | 0.0427 | 0.1572 | 0.2760 | 1.3801 | 0.180 | 0.164 | 9 | 16 | 0 | 36.0000 | 64.0000 | 0.180 | 0.164 |
| Two-second peri-hit | Sensorimotor | Aperiodic exponent | 25 | 0.8735 | 0.9297 | 0.9230 | 0.8809 | 0.0562 | 0.0552 | 0.1470 | 0.3823 | 1.9114 | 0.068 | 0.037 | 10 | 15 | 0 | 40.0000 | 60.0000 | 0.082 | 0.044 |
| Two-second peri-hit | Parietal | Aperiodic exponent | 25 | 0.6591 | 0.7828 | 0.6129 | 0.8006 | 0.1237 | 0.1197 | 0.1474 | 0.8391 | 4.1955 | <0.001 | <0.001 | 6 | 19 | 0 | 24.0000 | 76.0000 | <0.001 | <0.001 |
| Two-second peri-hit | Posterior | Aperiodic exponent | 25 | 0.4517 | 0.6011 | 0.3431 | 0.5686 | 0.1494 | 0.1395 | 0.1147 | 1.3028 | 6.5139 | <0.001 | <0.001 | 2 | 23 | 0 | 8.0000 | 92.0000 | <0.001 | <0.001 |
| Two-second peri-hit | Temporal | Aperiodic exponent | 25 | 0.3132 | 0.4086 | 0.2052 | 0.3546 | 0.0954 | 0.0914 | 0.2115 | 0.4511 | 2.2554 | 0.034 | 0.037 | 7 | 18 | 0 | 28.0000 | 72.0000 | 0.050 | 0.044 |
| Two-second peri-hit | Other | Aperiodic exponent | 25 | 0.6080 | 0.7420 | 0.5734 | 0.7089 | 0.1340 | 0.1084 | 0.1577 | 0.8496 | 4.2478 | <0.001 | <0.001 | 2 | 23 | 0 | 8.0000 | 92.0000 | <0.001 | <0.001 |
| Two-second peri-hit | Frontal | Aperiodic offset | 25 | -10.4043 | -10.4365 | -10.4121 | -10.5078 | -0.0322 | -0.0057 | 0.1987 | -0.1621 | -0.8103 | 0.426 | 0.491 | 13 | 12 | 0 | 52.0000 | 48.0000 | 0.639 | 0.736 |
| Two-second peri-hit | Sensorimotor | Aperiodic offset | 25 | -11.2976 | -11.3105 | -11.2453 | -11.2446 | -0.0129 | 0.0057 | 0.1735 | -0.0745 | -0.3725 | 0.713 | 0.874 | 12 | 13 | 0 | 48.0000 | 52.0000 | 0.727 | 0.874 |
| Two-second peri-hit | Parietal | Aperiodic offset | 25 | -11.2496 | -11.1942 | -11.2682 | -11.1699 | 0.0554 | 0.0599 | 0.1365 | 0.4061 | 2.0307 | 0.054 | 0.085 | 8 | 17 | 0 | 32.0000 | 68.0000 | 0.161 | 0.255 |
| Two-second peri-hit | Posterior | Aperiodic offset | 25 | -11.1019 | -11.0327 | -11.1288 | -11.1096 | 0.0692 | 0.0784 | 0.0951 | 0.7278 | 3.6391 | 0.001 | <0.001 | 4 | 21 | 0 | 16.0000 | 84.0000 | 0.008 | 0.004 |
| Two-second peri-hit | Temporal | Aperiodic offset | 25 | -11.3510 | -11.3320 | -11.3538 | -11.2941 | 0.0190 | 0.0474 | 0.2688 | 0.0705 | 0.3526 | 0.727 | 0.634 | 11 | 14 | 0 | 44.0000 | 56.0000 | 0.727 | 0.761 |
| Two-second peri-hit | Other | Aperiodic offset | 25 | -10.8050 | -10.7514 | -10.7771 | -10.7442 | 0.0535 | 0.0210 | 0.1777 | 0.3013 | 1.5064 | 0.145 | 0.200 | 10 | 15 | 0 | 40.0000 | 60.0000 | 0.290 | 0.400 |

### Section 9. Two-second max6 bootstrap

| Window | Sensor region | Feature | Participants | Bootstrap resamples | Random seed | Observed mean difference | Mean difference 95% CI, lower | Mean difference 95% CI, upper | Observed Cohen's dz | Cohen's dz 95% CI, lower | Cohen's dz 95% CI, upper | Bootstrap fraction mean > 0 |
|---|---|---|---|---|---|---|---|---|---|---|---|---|
| Two-second peri-hit | Parietal | Aperiodic exponent | 25 | 50000 | 20260722 | 0.1237 | 0.0680 | 0.1813 | 0.8391 | 0.4904 | 1.3314 | 1.0000 |

### Section 10. Two-second max6 leave-one-participant-out

| Excluded participant | Window | Sensor region | Feature | Participants | Human mean | Machine mean | Human median | Machine median | Machine − human mean | Machine − human median | Difference SD | Cohen's dz | Paired t | Paired t p | Wilcoxon p | Machine lower, n | Machine higher, n | Equal, n | Machine lower (%) | Machine higher (%) | Paired t FDR q | Wilcoxon FDR q |
|---|---|---|---|---|---|---|---|---|---|---|---|---|---|---|---|---|---|---|---|---|---|---|
| Participant 01 | Two-second peri-hit | Parietal | Aperiodic exponent | 24 | 0.6654 | 0.7816 | 0.6483 | 0.7751 | 0.1162 | 0.1074 | 0.1457 | 0.7979 | 3.9087 | <0.001 | <0.001 | 6 | 18 | 0 | 25.0000 | 75.0000 | 0.001 | <0.001 |
| Participant 02 | Two-second peri-hit | Parietal | Aperiodic exponent | 24 | 0.6627 | 0.7760 | 0.6483 | 0.7751 | 0.1134 | 0.1074 | 0.1410 | 0.8038 | 3.9377 | <0.001 | <0.001 | 6 | 18 | 0 | 25.0000 | 75.0000 | 0.001 | <0.001 |
| Participant 03 | Two-second peri-hit | Parietal | Aperiodic exponent | 24 | 0.6727 | 0.7976 | 0.6483 | 0.8020 | 0.1249 | 0.1216 | 0.1504 | 0.8300 | 4.0664 | <0.001 | <0.001 | 6 | 18 | 0 | 25.0000 | 75.0000 | <0.001 | <0.001 |
| Participant 04 | Two-second peri-hit | Parietal | Aperiodic exponent | 24 | 0.6526 | 0.7785 | 0.5970 | 0.7751 | 0.1259 | 0.1216 | 0.1501 | 0.8386 | 4.1081 | <0.001 | <0.001 | 6 | 18 | 0 | 25.0000 | 75.0000 | <0.001 | <0.001 |
| Participant 05 | Two-second peri-hit | Parietal | Aperiodic exponent | 24 | 0.6530 | 0.7641 | 0.5970 | 0.7751 | 0.1111 | 0.1074 | 0.1362 | 0.8159 | 3.9970 | <0.001 | <0.001 | 6 | 18 | 0 | 25.0000 | 75.0000 | 0.001 | <0.001 |
| Participant 06 | Two-second peri-hit | Parietal | Aperiodic exponent | 24 | 0.6773 | 0.7972 | 0.6483 | 0.8020 | 0.1199 | 0.1074 | 0.1493 | 0.8029 | 3.9334 | <0.001 | <0.001 | 6 | 18 | 0 | 25.0000 | 75.0000 | 0.001 | <0.001 |
| Participant 07 | Two-second peri-hit | Parietal | Aperiodic exponent | 24 | 0.6485 | 0.7821 | 0.5970 | 0.7765 | 0.1336 | 0.1216 | 0.1418 | 0.9420 | 4.6147 | <0.001 | <0.001 | 5 | 19 | 0 | 20.8333 | 79.1667 | <0.001 | <0.001 |
| Participant 08 | Two-second peri-hit | Parietal | Aperiodic exponent | 24 | 0.6794 | 0.8102 | 0.6483 | 0.8020 | 0.1309 | 0.1216 | 0.1460 | 0.8960 | 4.3894 | <0.001 | <0.001 | 5 | 19 | 0 | 20.8333 | 79.1667 | <0.001 | <0.001 |
| Participant 09 | Two-second peri-hit | Parietal | Aperiodic exponent | 24 | 0.6698 | 0.8023 | 0.6483 | 0.8020 | 0.1325 | 0.1216 | 0.1437 | 0.9223 | 4.5181 | <0.001 | <0.001 | 5 | 19 | 0 | 20.8333 | 79.1667 | <0.001 | <0.001 |
| Participant 10 | Two-second peri-hit | Parietal | Aperiodic exponent | 24 | 0.6556 | 0.7848 | 0.5970 | 0.8020 | 0.1292 | 0.1216 | 0.1479 | 0.8731 | 4.2773 | <0.001 | <0.001 | 5 | 19 | 0 | 20.8333 | 79.1667 | <0.001 | <0.001 |
| Participant 11 | Two-second peri-hit | Parietal | Aperiodic exponent | 24 | 0.6525 | 0.7678 | 0.5970 | 0.7751 | 0.1153 | 0.1074 | 0.1443 | 0.7988 | 3.9131 | <0.001 | <0.001 | 6 | 18 | 0 | 25.0000 | 75.0000 | 0.001 | <0.001 |
| Participant 12 | Two-second peri-hit | Parietal | Aperiodic exponent | 24 | 0.6766 | 0.8026 | 0.6483 | 0.8020 | 0.1259 | 0.1216 | 0.1501 | 0.8389 | 4.1097 | <0.001 | <0.001 | 6 | 18 | 0 | 25.0000 | 75.0000 | <0.001 | <0.001 |
| Participant 13 | Two-second peri-hit | Parietal | Aperiodic exponent | 24 | 0.6559 | 0.7747 | 0.5970 | 0.7751 | 0.1189 | 0.1074 | 0.1485 | 0.8002 | 3.9200 | <0.001 | <0.001 | 6 | 18 | 0 | 25.0000 | 75.0000 | 0.001 | <0.001 |
| Participant 14 | Two-second peri-hit | Parietal | Aperiodic exponent | 24 | 0.6624 | 0.7842 | 0.6483 | 0.8020 | 0.1218 | 0.1074 | 0.1503 | 0.8107 | 3.9715 | <0.001 | <0.001 | 6 | 18 | 0 | 25.0000 | 75.0000 | 0.001 | <0.001 |
| Participant 15 | Two-second peri-hit | Parietal | Aperiodic exponent | 24 | 0.6610 | 0.7887 | 0.6324 | 0.8020 | 0.1277 | 0.1216 | 0.1492 | 0.8559 | 4.1930 | <0.001 | <0.001 | 6 | 18 | 0 | 25.0000 | 75.0000 | <0.001 | <0.001 |
| Participant 16 | Two-second peri-hit | Parietal | Aperiodic exponent | 24 | 0.6463 | 0.7779 | 0.5970 | 0.7751 | 0.1316 | 0.1216 | 0.1451 | 0.9070 | 4.4433 | <0.001 | <0.001 | 5 | 19 | 0 | 20.8333 | 79.1667 | <0.001 | <0.001 |
| Participant 17 | Two-second peri-hit | Parietal | Aperiodic exponent | 24 | 0.6627 | 0.7846 | 0.6483 | 0.8020 | 0.1219 | 0.1074 | 0.1503 | 0.8113 | 3.9744 | <0.001 | <0.001 | 6 | 18 | 0 | 25.0000 | 75.0000 | 0.001 | <0.001 |
| Participant 18 | Two-second peri-hit | Parietal | Aperiodic exponent | 24 | 0.6641 | 0.7897 | 0.6483 | 0.8020 | 0.1256 | 0.1216 | 0.1503 | 0.8359 | 4.0953 | <0.001 | <0.001 | 6 | 18 | 0 | 25.0000 | 75.0000 | 0.001 | <0.001 |
| Participant 19 | Two-second peri-hit | Parietal | Aperiodic exponent | 24 | 0.6675 | 0.7911 | 0.6483 | 0.8020 | 0.1237 | 0.1074 | 0.1506 | 0.8213 | 4.0235 | <0.001 | <0.001 | 6 | 18 | 0 | 25.0000 | 75.0000 | 0.001 | <0.001 |
| Participant 20 | Two-second peri-hit | Parietal | Aperiodic exponent | 24 | 0.6482 | 0.7686 | 0.5970 | 0.7751 | 0.1204 | 0.1074 | 0.1496 | 0.8047 | 3.9421 | <0.001 | <0.001 | 6 | 18 | 0 | 25.0000 | 75.0000 | 0.001 | <0.001 |
| Participant 21 | Two-second peri-hit | Parietal | Aperiodic exponent | 24 | 0.6377 | 0.7548 | 0.5970 | 0.7751 | 0.1171 | 0.1074 | 0.1468 | 0.7979 | 3.9088 | <0.001 | <0.001 | 6 | 18 | 0 | 25.0000 | 75.0000 | 0.001 | <0.001 |
| Participant 22 | Two-second peri-hit | Parietal | Aperiodic exponent | 24 | 0.6780 | 0.8096 | 0.6483 | 0.8020 | 0.1316 | 0.1216 | 0.1451 | 0.9066 | 4.4413 | <0.001 | <0.001 | 5 | 19 | 0 | 20.8333 | 79.1667 | <0.001 | <0.001 |
| Participant 23 | Two-second peri-hit | Parietal | Aperiodic exponent | 24 | 0.6331 | 0.7587 | 0.5970 | 0.7751 | 0.1256 | 0.1216 | 0.1502 | 0.8361 | 4.0961 | <0.001 | <0.001 | 6 | 18 | 0 | 25.0000 | 75.0000 | <0.001 | <0.001 |
| Participant 24 | Two-second peri-hit | Parietal | Aperiodic exponent | 24 | 0.6581 | 0.7819 | 0.5970 | 0.7751 | 0.1239 | 0.1092 | 0.1506 | 0.8226 | 4.0297 | <0.001 | <0.001 | 6 | 18 | 0 | 25.0000 | 75.0000 | 0.001 | <0.001 |
| Participant 25 | Two-second peri-hit | Parietal | Aperiodic exponent | 24 | 0.6370 | 0.7607 | 0.5970 | 0.7751 | 0.1237 | 0.1074 | 0.1506 | 0.8215 | 4.0246 | <0.001 | <0.001 | 6 | 18 | 0 | 25.0000 | 75.0000 | 0.001 | <0.001 |

### Section 11. Two-second fit-quality association

| Window | Sensor region | Spearman ρ | Effect difference | Fit-quality difference | Spearman p | Exploratory | Spearman FDR q |
|---|---|---|---|---|---|---|---|
| Two-second peri-hit | Parietal | 0.6369 | machine_minus_human_aperiodic exponent | Machine − human $R^2$ | <0.001 | Yes | 0.001 |
| Two-second peri-hit | Parietal | -0.5162 | machine_minus_human_aperiodic exponent | Machine − human model error | 0.008 | Yes | 0.008 |

### Section 12. Two-second fit-quality influence audit

| Participants | Spearman ρ | Predictor | Predictor variable | Spearman permutation p | OLS intercept at Δquality = 0 | OLS intercept HC3 p | OLS intercept HC3 CI, lower | OLS intercept HC3 CI, upper | OLS slope | OLS slope HC3 p | OLS slope HC3 CI, lower | OLS slope HC3 CI, upper | OLS $R^2$ | Theil–Sen intercept | Theil–Sen slope | Theil–Sen slope CI, lower | Theil–Sen slope CI, upper | Maximum Cook's distance | Cook's distance > 4/n, n | Cook's-distance threshold (4/n) | Interpretation |
|---|---|---|---|---|---|---|---|---|---|---|---|---|---|---|---|---|---|---|---|---|---|
| 25 | 0.6369 | $\Delta R^2$ | Machine − human $R^2$ | <0.001 | 0.1051 | <0.001 | 0.0548 | 0.1554 | 0.8129 | 0.047 | 0.0110 | 1.6147 | 0.2841 | 0.1097 | 1.2428 | 0.5688 | 2.3352 | 1.1376 | 1 | 0.1600 | Exploratory only; fit quality and exponent are outputs of the same spectral model. |
| 25 | -0.5162 | Δ model error | Machine − human model error | 0.009 | 0.0950 | <0.001 | 0.0470 | 0.1429 | -9.3090 | 0.009 | -16.3376 | -2.2804 | 0.2692 | 0.1030 | -11.4044 | -17.6718 | -3.8406 | 0.4776 | 1 | 0.1600 | Exploratory only; fit quality and exponent are outputs of the same spectral model. |

### Section 13. Fit-quality-neutral intercept bootstrap

| Bootstrap resamples | Random seed | Predictor | Intercept at Δquality = 0, 95% CI lower | Intercept at Δquality = 0, 95% CI upper | Bootstrap fraction intercept > 0 | Observed OLS intercept at Δquality = 0 |
|---|---|---|---|---|---|---|
| 50000 | 20260822 | $\Delta R^2$ | 0.0586 | 0.1579 | 1.0000 | 0.1051 |
| 50000 | 20260823 | Δ model error | 0.0531 | 0.1461 | 1.0000 | 0.0950 |

### Section 14. Aperiodic-only focal-model comparison

| Model | Window | Sensor region | Feature | Participants | Human mean | Machine mean | Machine − human mean | Machine − human median | Difference SD | Cohen's dz | Paired t | Paired t p | Wilcoxon p | Machine lower, n | Machine higher, n | Equal, n | Machine higher (%) | Paired t FDR q | Wilcoxon FDR q |
|---|---|---|---|---|---|---|---|---|---|---|---|---|---|---|---|---|---|---|---|
| Two-second max6 | Two-second peri-hit | Parietal | Aperiodic exponent | 25 | 0.6591 | 0.7828 | 0.1237 | 0.1197 | 0.1474 | 0.8391 | 4.1955 | <0.001 | <0.001 | 6 | 19 | 0 | 76.0000 | <0.001 | <0.001 |
| Two-second aperiodic-only | Two-second peri-hit | Parietal | Aperiodic exponent | 25 | 0.6291 | 0.7509 | 0.1218 | 0.1084 | 0.1548 | 0.7869 | 3.9346 | <0.001 | <0.001 | 5 | 20 | 0 | 80.0000 | 0.001 | 0.001 |

### Section 15. Aperiodic-only bootstrap

| Model | Participants | Bootstrap resamples | Random seed | Observed mean difference | Mean difference 95% CI, lower | Mean difference 95% CI, upper | Observed Cohen's dz | Cohen's dz 95% CI, lower | Cohen's dz 95% CI, upper | Bootstrap fraction mean > 0 |
|---|---|---|---|---|---|---|---|---|---|---|
| Two-second max6 | 25 | 50000 | 20262309 | 0.1237 | 0.0680 | 0.1802 | 0.8391 | 0.4955 | 1.3299 | 1.0000 |
| Two-second aperiodic-only | 25 | 50000 | 20263418 | 0.1218 | 0.0633 | 0.1818 | 0.7869 | 0.4250 | 1.2964 | 1.0000 |

### Section 16. Aperiodic-only leave-one-participant-out

| Model | Excluded participant | Window | Sensor region | Feature | Participants | Human mean | Machine mean | Machine − human mean | Machine − human median | Difference SD | Cohen's dz | Paired t | Paired t p | Wilcoxon p | Machine lower, n | Machine higher, n | Equal, n | Machine higher (%) | Paired t FDR q | Wilcoxon FDR q |
|---|---|---|---|---|---|---|---|---|---|---|---|---|---|---|---|---|---|---|---|---|
| Two-second aperiodic-only | Participant 01 | Two-second peri-hit | Parietal | Aperiodic exponent | 24 | 0.6341 | 0.7484 | 0.1142 | 0.1024 | 0.1533 | 0.7451 | 3.6502 | 0.001 | 0.002 | 5 | 19 | 0 | 79.1667 | 0.003 | 0.003 |
| Two-second aperiodic-only | Participant 02 | Two-second peri-hit | Parietal | Aperiodic exponent | 24 | 0.6330 | 0.7436 | 0.1106 | 0.1024 | 0.1473 | 0.7504 | 3.6763 | 0.001 | 0.002 | 5 | 19 | 0 | 79.1667 | 0.002 | 0.003 |
| Two-second aperiodic-only | Participant 03 | Two-second peri-hit | Parietal | Aperiodic exponent | 24 | 0.6399 | 0.7622 | 0.1222 | 0.1024 | 0.1581 | 0.7733 | 3.7882 | <0.001 | 0.001 | 5 | 19 | 0 | 79.1667 | 0.002 | 0.002 |
| Two-second aperiodic-only | Participant 04 | Two-second peri-hit | Parietal | Aperiodic exponent | 24 | 0.6242 | 0.7456 | 0.1214 | 0.1024 | 0.1581 | 0.7679 | 3.7618 | 0.001 | 0.002 | 5 | 19 | 0 | 79.1667 | 0.001 | 0.002 |
| Two-second aperiodic-only | Participant 05 | Two-second peri-hit | Parietal | Aperiodic exponent | 24 | 0.6230 | 0.7330 | 0.1100 | 0.1024 | 0.1461 | 0.7526 | 3.6871 | 0.001 | 0.002 | 5 | 19 | 0 | 79.1667 | 0.002 | 0.003 |
| Two-second aperiodic-only | Participant 06 | Two-second peri-hit | Parietal | Aperiodic exponent | 24 | 0.6492 | 0.7679 | 0.1187 | 0.1024 | 0.1573 | 0.7548 | 3.6976 | 0.001 | 0.002 | 5 | 19 | 0 | 79.1667 | 0.002 | 0.003 |
| Two-second aperiodic-only | Participant 07 | Two-second peri-hit | Parietal | Aperiodic exponent | 24 | 0.6174 | 0.7511 | 0.1337 | 0.1097 | 0.1460 | 0.9158 | 4.4866 | <0.001 | <0.001 | 4 | 20 | 0 | 83.3333 | <0.001 | <0.001 |
| Two-second aperiodic-only | Participant 08 | Two-second peri-hit | Parietal | Aperiodic exponent | 24 | 0.6516 | 0.7800 | 0.1285 | 0.1097 | 0.1544 | 0.8321 | 4.0763 | <0.001 | <0.001 | 4 | 20 | 0 | 83.3333 | <0.001 | 0.001 |
| Two-second aperiodic-only | Participant 09 | Two-second peri-hit | Parietal | Aperiodic exponent | 24 | 0.6435 | 0.7691 | 0.1256 | 0.1097 | 0.1569 | 0.8007 | 3.9228 | <0.001 | 0.001 | 5 | 19 | 0 | 79.1667 | 0.001 | 0.002 |
| Two-second aperiodic-only | Participant 10 | Two-second peri-hit | Parietal | Aperiodic exponent | 24 | 0.6253 | 0.7529 | 0.1276 | 0.1097 | 0.1553 | 0.8220 | 4.0271 | <0.001 | <0.001 | 4 | 20 | 0 | 83.3333 | 0.001 | 0.001 |
| Two-second aperiodic-only | Participant 11 | Two-second peri-hit | Parietal | Aperiodic exponent | 24 | 0.6188 | 0.7343 | 0.1155 | 0.1024 | 0.1548 | 0.7462 | 3.6555 | 0.001 | 0.002 | 5 | 19 | 0 | 79.1667 | 0.003 | 0.003 |
| Two-second aperiodic-only | Participant 12 | Two-second peri-hit | Parietal | Aperiodic exponent | 24 | 0.6501 | 0.7744 | 0.1243 | 0.1097 | 0.1576 | 0.7889 | 3.8650 | <0.001 | 0.001 | 5 | 19 | 0 | 79.1667 | 0.002 | 0.002 |
| Two-second aperiodic-only | Participant 13 | Two-second peri-hit | Parietal | Aperiodic exponent | 24 | 0.6286 | 0.7431 | 0.1145 | 0.1024 | 0.1537 | 0.7452 | 3.6508 | 0.001 | 0.002 | 5 | 19 | 0 | 79.1667 | 0.003 | 0.003 |
| Two-second aperiodic-only | Participant 14 | Two-second peri-hit | Parietal | Aperiodic exponent | 24 | 0.6287 | 0.7535 | 0.1248 | 0.1097 | 0.1573 | 0.7934 | 3.8869 | <0.001 | 0.001 | 5 | 19 | 0 | 79.1667 | 0.001 | 0.002 |
| Two-second aperiodic-only | Participant 15 | Two-second peri-hit | Parietal | Aperiodic exponent | 24 | 0.6263 | 0.7517 | 0.1254 | 0.1097 | 0.1570 | 0.7987 | 3.9130 | <0.001 | 0.001 | 5 | 19 | 0 | 79.1667 | 0.001 | 0.001 |
| Two-second aperiodic-only | Participant 16 | Two-second peri-hit | Parietal | Aperiodic exponent | 24 | 0.6127 | 0.7449 | 0.1322 | 0.1097 | 0.1489 | 0.8876 | 4.3484 | <0.001 | <0.001 | 4 | 20 | 0 | 83.3333 | <0.001 | <0.001 |
| Two-second aperiodic-only | Participant 17 | Two-second peri-hit | Parietal | Aperiodic exponent | 24 | 0.6368 | 0.7586 | 0.1217 | 0.1024 | 0.1581 | 0.7699 | 3.7719 | <0.001 | 0.001 | 5 | 19 | 0 | 79.1667 | 0.002 | 0.002 |
| Two-second aperiodic-only | Participant 18 | Two-second peri-hit | Parietal | Aperiodic exponent | 24 | 0.6295 | 0.7542 | 0.1247 | 0.1097 | 0.1574 | 0.7926 | 3.8828 | <0.001 | 0.001 | 5 | 19 | 0 | 79.1667 | 0.002 | 0.002 |
| Two-second aperiodic-only | Participant 19 | Two-second peri-hit | Parietal | Aperiodic exponent | 24 | 0.6370 | 0.7598 | 0.1228 | 0.1097 | 0.1580 | 0.7775 | 3.8088 | <0.001 | 0.001 | 5 | 19 | 0 | 79.1667 | 0.002 | 0.002 |
| Two-second aperiodic-only | Participant 20 | Two-second peri-hit | Parietal | Aperiodic exponent | 24 | 0.6209 | 0.7365 | 0.1156 | 0.1024 | 0.1549 | 0.7463 | 3.6562 | 0.001 | 0.002 | 5 | 19 | 0 | 79.1667 | 0.002 | 0.002 |
| Two-second aperiodic-only | Participant 21 | Two-second peri-hit | Parietal | Aperiodic exponent | 24 | 0.6066 | 0.7201 | 0.1135 | 0.1024 | 0.1524 | 0.7451 | 3.6503 | 0.001 | 0.002 | 5 | 19 | 0 | 79.1667 | 0.003 | 0.003 |
| Two-second aperiodic-only | Participant 22 | Two-second peri-hit | Parietal | Aperiodic exponent | 24 | 0.6460 | 0.7769 | 0.1309 | 0.1097 | 0.1511 | 0.8660 | 4.2424 | <0.001 | <0.001 | 4 | 20 | 0 | 83.3333 | <0.001 | <0.001 |
| Two-second aperiodic-only | Participant 23 | Two-second peri-hit | Parietal | Aperiodic exponent | 24 | 0.6111 | 0.7344 | 0.1233 | 0.1097 | 0.1579 | 0.7807 | 3.8244 | <0.001 | 0.001 | 5 | 19 | 0 | 79.1667 | 0.002 | 0.002 |
| Two-second aperiodic-only | Participant 24 | Two-second peri-hit | Parietal | Aperiodic exponent | 24 | 0.6263 | 0.7487 | 0.1223 | 0.1038 | 0.1581 | 0.7740 | 3.7919 | <0.001 | 0.001 | 5 | 19 | 0 | 79.1667 | 0.002 | 0.002 |
| Two-second aperiodic-only | Participant 25 | Two-second peri-hit | Parietal | Aperiodic exponent | 24 | 0.6067 | 0.7271 | 0.1204 | 0.1024 | 0.1579 | 0.7624 | 3.7351 | 0.001 | 0.002 | 5 | 19 | 0 | 79.1667 | 0.002 | 0.002 |

### Section 17. Aperiodic-only model concordance

| Participants | Model A | Model B | Pearson r | Spearman ρ | Mean difference between model effects | Mean absolute effect difference | Maximum absolute effect difference | Participants changing direction, n | Direction agreement fraction |
|---|---|---|---|---|---|---|---|---|---|
| 25 | Two-second max6 | Two-second aperiodic-only | 0.9488 | 0.9238 | -0.0019 | 0.0367 | 0.1196 | 1 | 0.9600 |

*Note.* BIC = Bayesian information criterion; lower BIC indicates the preferred model. In BIC sections, max6 is model B1 and max3 is model B2, so ΔBIC = max3 − max6. Positive paired effects are machine minus human. Bootstrap confidence intervals are paired-participant intervals. Leave-one-participant-out sections repeat the analysis after omitting each participant. Fit-quality association and influence analyses are exploratory because exponent and fit quality are outputs of the same spectral model.
